# RNA interference pathways display high rates of adaptive protein evolution across multiple invertebrates

**DOI:** 10.1101/154153

**Authors:** William H. Palmer, Jarrod Hadfield, Darren J. Obbard

## Abstract

Conflict between organisms can lead to reciprocal adaptation that manifests itself as an increased evolutionary rate in the genes mediating the conflict. This adaptive signature has been observed in RNA interference (RNAi) pathway genes involved in the suppression of viruses and transposable elements in *Drosophila melanogaster*, suggesting that a subset of *Drosophila* RNAi genes may be locked into an arms race with these parasites. However, it is not known whether rapid evolution of RNAi genes is a general phenomenon across invertebrates, or which RNAi genes generally evolve adaptively. Here we use population genomic data from eight invertebrate species to infer rates of adaptive sequence evolution, and to test for past and ongoing selective sweeps in RNAi genes. We assess rates of adaptive protein evolution across species by using a formal meta-analytic framework to combine data across species, and by implementing a multispecies generalised linear mixed model of mutation counts. In all species, we find that RNAi genes display a greater rate of adaptive protein substitution than other genes, and that this is primarily mediated by positive selection acting on the subset of genes that are most likely to defend against viruses and transposable elements. In contrast, evidence for recent selective sweeps is broadly spread across functional classes of RNAi genes and differs substantially among species. Finally, we identify genes that exhibit elevated adaptive evolution across the analysed insect species combined, perhaps due to concurrent parasite-mediated arms races.

## Introduction

RNA-interference mechanisms include a diverse group of pathways, united by their use of Argonaute-family proteins complexed with short (20-30 nt) RNA molecules to guide the targeting of longer RNA molecules through sequence complementarity (Carmell, et al., 2002; Meister, 2013). These pathways regulate multiple biological processes that can be divided into three distinct subpathways in arthropods and nematodes, each represented by a characteristic class of small RNAs: the micro-RNA (miRNA), the short-interfering RNA (siRNA), and the piwi-interacting RNA (piRNA) pathways. The miRNA pathway processes endogenously-encoded foldback hairpins which, once in their mature miRNA form, regulate gene expression and coordinate developmental processes (Alvarez-Garcia & Miska, 2005; Chen, et al., 2014; Ha & Kim, 2014). The siRNA pathway has two distinct roles, depending on the endogenous or exogenous origin of its substrate. First, the endo-siRNA pathway processes endogenously encoded dsRNA to regulate processes such as TE defense (Kawamura, et al., 2008; Czech, et al., 2008; Ghildiyal, et al., 2008) chromosomal segregation (Hall, et al., 2003; Huang, et al., 2015), and heterochromatin formation (Deshpande, et al., 2005). Second, the exo-siRNA (or viRNA) functions primarily as a form of antiviral immunity (Wang, et al., 2006; Bronkhorst & van Rij, 2014). The piRNA pathway forms a defence against transposable elements (TEs) in the germ line, and piRNAs are derived from endogenously-encoded piRNA clusters of inactivated TE sequences and from active TEs (Klattenhoff & Theurkauf, 2008; Thomson & Lin, 2009; Czech, et al., 2016).

Nevertheless, within this simple framework there is substantial variation among species, and RNAi-pathway components seem to be evolutionarily labile. For example, in nematodes the mechanism and function of the piRNA pathway is not well conserved: primary piRNA-like small RNAs are encoded by short distinct loci instead of the clusters observed in flies and mammals, and mediate the biogenesis of a separate endo-siRNA population transcribed by an RNA-dependent RNA Polymerase (RdRP) and processed by Dicer (Duchaine, et al., 2006; Das, et al., 2008). Further, only one of the five major clades of nematode have retained Piwi-subfamily proteins — the canonical effector of the piRNA pathway—and instead rely solely on the (RDRP-produced) endo-siRNAs (Sarkies, et al., 2015). The piRNA pathway can also take on entirely new roles, for example, multiple duplications of *piwi* in *Aedes* mosquitoes has allowed the piRNA pathway to adopt an antiviral role in the somatic tissues (Morazzani, et al., 2012), while other *piwi* duplicates maintain the ancestral function (Miesen, et al., 2015; Miesen, et al., 2016).

The role of RNAi pathways in mediating inter-genomic (host-virus) and intra-genomic (host-TE, segregation distortion) (Ferree & Barbash, 2007) conflict suggests that they may be a hotspot of adaptive protein evolution. This has been well studied in *Drosophila*, where RNAi pathway genes show elevated rates of adaptive protein evolution (Obbard, et al., 2006; Obbard, et al., 2009), signatures of selective sweeps (Obbard, et al., 2011; Kolaczkowski, et al., 2011; Lewis, et al., 2016), and sites with elevated protein evolution across the *Drosophila* phylogeny (Vermaak, et al., 2005; Heger & Ponting, 2007; Kolaczkowski, et al., 2011). For example, a comparison of the antiviral RNAi genes *AGO2*, *Dcr-2*, and *r2d2* to their miRNA functional counterparts with no known role in conflict (the paralogs *AGO1*, *Dcr-1*, and *loqs*) shows a striking difference in rates of protein evolution, as well as a greater rate of adaptive amino-acid substitution (Obbard, et al., 2006). In addition, evolutionary rates of piRNA pathway genes involved in transcriptional silencing are elevated and highly correlated with other piRNA pathway genes across the *Drosophila* phylogeny (Blumenstiel, et al., 2016).

Although some antiviral and anti-TE RNAi pathway genes clearly display elevated rates of adaptive protein evolution in *Drosophila*, the generality of this pattern remains to be elucidated. Here we apply both traditional McDonald-Kreitman (McDonald & Kreitman, 1991) and SnIPRE-style (Eilertson, et al., 2012) analyses, and selective sweep-based analyses (Nielsen, et al., 2005; Pavlidis, et al., 2013) to publicly-available genome-scale data from 6 insects and 2 nematodes. By combining estimates across species, we investigate the specific RNAi subpathways that may be the target of elevated positive selection. This allows us to estimate the rates of adaptation across species, thereby improving single gene estimates and allowing us to identify genes that are undergoing parallel adaptation across the taxa analysed. Finally, we summarise the evidence for recently completed and ongoing selective sweeps in RNAi genes across these eight taxa. We conclude that rapid evolution of RNAi genes is a general phenomenon in these invertebrates, although evidence for recent sweeps is highly contingent on the focal species.

## Materials and Methods

### Selection of genes for analysis

Genes implicated in the RNAi pathway of either *Drosophila melanogaster* or *Caenorhabditis elegans* were used to find homologues in six insects and two nematode species (Table S1, Table S2). For the six insect species, these were further classified as miRNA, piRNA, siRNA, or viRNA. Although the viRNA pathway is not widely regarded as separate from siRNA, we make this distinction based on the hypothesis that these genes may be evolving adaptively in response to viruses, as these genes have direct experimental evidence of an antiviral role in *D. melanogaster*. We also split the piRNA pathway genes among three functional categories: post-transcriptional silencing effectors, transcriptional silencing effectors, and biogenesis machinery. A gene was considered a biogenesis factor if piRNA levels decrease upon loss-of-function, an effector if piRNA pathway function is compromised without reducing piRNA levels, and a transcriptional silencing effector if the effector is involved in transcriptional silencing (Table S1). Finally, we selected 65 piRNA genes in *D. melanogaster* with known tissue-specificity to calculate rates of adaptation in the germline versus the somatic follicle cells (Table S3). This gene list contains the core of the piRNA pathway and genes independently validated in two of the three recent screens for piRNA pathway constituents (Handler, et al., 2013; Czech, et al., 2013; Muerdter, et al., 2013).

Homologs of the *D. melanogaster* and *C. elegans* genes were identified using a two-step process. First, a hidden Markov Model (HMMer) (Eddy, 2008)) was used to find best reciprocal best-hits for a gene of interest using predicted protein sets (if available) or UniProtKB. If no hit was found, then Exonerate was used to identify unannotated homologues in the genome using the model ‘protein2genome’ (Slater & Birney, 2005). If exonerate was unable to model a homologue, then this gene was classified as missing, either due to gene loss or an incomplete genome assembly. We defined genes as duplicates (paralogues) if multiple regions of a genome shared a best hit to a reference gene, and these regions showed substantial sequence divergence between them (i.e. they were not obviously a mis-assembly duplicate or allelic). Because of the large divergence times between insects and nematodes and the complexity of RNAi pathways in nematodes, and hence the associated difficulty in assigning homology in the two nematodes, we restricted our gene-level analyses to only the insect species.

### Population genomic data

We utilised previously published population genomic data for *Drosophila melanogaster* (Lack et al, 2015), *Drosophila pseudoobscura* (Pseudobase) (McGaugh, et al., 2012), *Anopheles gambiae* (The *Anopheles gambiae* 1000 Genomes Consortium (2014): Ag1000G phase 1 AR2 data release. MalariaGEN.), *Heliconius melpomene* (Kronforst, et al., 2013), *Bombyx mandarina* (Xia, et al., 2009), *Apis mellifera* (Harpur, et al., 2014), *Pristionchus pacificus* (Rödelsperger, et al., 2014), and *Caenorhabditis briggsae* (Thomas, et al., 2015) for our analyses (Table S4). For both *Drosophila* species, we used previously-published haplotype data (haploid sequencing of *D. melanogaster*, inbred lines of *D. pseudoobscura*). For the other taxa we obtained raw sequencing reads from EBI ENA (identifiers provided in Table S4) and mapped them to the most recent reference genome for each species using Bowtie2 (Langmead & Salzberg, 2012) with default settings. We used GATK’s HaplotypeCaller on each individual separately (DePristo, et al., 2011) to call variants in a 200 kb region surrounding each gene of interest. For high coverage datasets (*A. mellifera, H. melpomene, C. briggsae, A. gambiae,* and *P. pacificus*) we excluded sites with a read depth lower than 5, but we reduced this threshold to 2 for the low-coverage *B. mandarina*. After mapping and filtering sites we created two randomly resolved pseudohaplotype sequences per individual (i.e. without any linkage information) from the sites that remained, and these were used for downstream analyses (none of which depend on linkage information). Only one haplotype was sampled from each *C. briggsae* and *P. pacificus* individual as the sequenced individuals were reported to be highly homozygous. In *H. melpomene*, we occasionally observed long stretches of high divergence shared by multiple individuals. We assumed these to be possible cases of either contamination, inversions that have recently risen to a high frequency, or introgression (Pardo-Diaz, et al., 2012), and removed these haplotypes.

To calculate divergence between genes, and to polarise mutations for sweep analyses, we used the outgroup species *Drosophila simulans, Drosophila miranda, Heliconius hecale, Bombyx huttoni, Anopheles christyi* and *Anopheles melas, Apis cerana, Caenorhabditis nigoni,* and *Pristionchus exspectatus*, respectively (Table S4). Outgroups were chosen based on their divergence from the ingroup species (*ca.* 1-10% divergence of all sites) and on the availability of genomic data. For *A. gambiae* we tested outgroups with low (*An. melas*) and high (*An. christyi*) divergence times, as most *Anopheles* species are too close or too divergent to provide a robust outgroup for MK tests (Obbard, et al., 2007), and our results remain qualitatively the same for both outgroups (*A. melas* used for the presented analyses). For *D. simulans* (FlyBase, r2.02), *D. miranda* (Pseudobase, MSH22 strain), *A. melas* (VectorBase, CM1001059 strain, AmelC1 assembly), *A. christyi* (VectorBase, ACHKN1017 strain, AchrA1 assembly), *B. huttoni* (Sackton, et al., 2014) (BioProject PRJNA198873), and *P. exspectatus* (WormBase, Bioproject PRJEB6009), the outgroup reference assemblies were publicly available and used as provided. However, the *Caenorhabditis nigoni* reference assembly sequence (caenorhabditis.bio.ed.ac.uk/home/download) is contaminated with the more divergent nematode *Caenorhabditis afra* (Thomas, et al., 2015), and *Caenorhabditis nigoni* is the only current suitable outgroup for C. briggsae. We therefore applied a sliding window across the alignments between *C. nigoni* and *C. afra*, and excluded regions that were greater than 6 standard deviations from the mean divergence. Published reference assemblies were not available for *Apis cerana* and *Heliconius hecale*. To generate outgroup sequences for these species we iteratively remapped reads (*H. hecale:* ERR260306; *A. cerana:* SRR957079) to the respective *Apis mellifera* and *Heliconius melpomene* references, each time updating the previous reference with homozygous nonreference calls. These reads were mapped with Bowtie2 and then remapped with the divergent alignment software, Stampy (Lunter & Goodson, 2011). Homozygous nonreference calls (enriched for sites divergent between the ingroup and outgroup) were made with GATK’s HaplotypeCaller, with the heterozygosity parameter set to the expected divergence between species. Such sequences will not perfectly reflect the true outgroup sequence, and are expected to be biased toward the ingroup, downwardly biasing estimates of divergence in high-divergence regions. However, we confirmed that this approach works well by iteratively mapping *D. simulans* to *D. melanogaster*, and comparing the result with the known *D. simulans* assemblies (*K_S_*= 0.10 for iterative mapping vs *K_S_*=0.12 for the true assembly), and while bias probably remains, it is unlikely to spuriously elevate the inferred rates of one class of genes relative to the other. More generally, our approach to mapping, filtering, and variant calling may be prone to such biases, but they are unlikely to differentially affect gene classes of different function.

For MK analyses, target sequences were aligned as amino acids using MUSCLE (Edgar, 2004), and then each examined by eye to remove putative mis-alignments. Within-species data was aligned first, and then a consensus sequence of this alignment used to align against the outgroup sequence. Synonymous and nonsynonymous substitutions between species were inferred using codeml from the PAML package using the YN00 model (Yang & Nielsen, 2000), which estimates substitution rates using an approximation to maximum likelihood methods, while accounting for base composition differences between codon positions and differences in transition/transversion rates.

### Rates of adaptive evolution by pathway

To estimate the rate of adaptive protein evolution in different functional classes of gene, and to test for differences in rate between classes, we used two different approaches derived from the McDonald-Kreitman test (‘MK framework’) (McDonald & Kreitman, 1991). The MK framework combines polymorphism and divergence data from putatively neutral (synonymous) and potentially selected (nonsynonymous) variants to infer an excess of nonsynonymous fixations—beyond that expected under model of neutrality and constraint—that can be attributed to positive selection. We first used an explicit population-genetic model to estimate the number of adaptive nonsynonymous substitutions per site (DFE-alpha) (Eyre-Walker & Keightley, 2009). This approach has the advantage that it provides direct estimates of the parameters of interest, and explicitly models changes in population size and the distribution of deleterious fitness effects, which might otherwise bias estimates (Keightley & Eyre-Walker, 2007; Eyre-Walker & Keightley, 2009). However, as currently implemented, this method does not allow data to be directly combined between species. Therefore, to obtain more precise homologue- and pathway-based estimates we combined per-gene point estimates from DFE-alpha using a linear mixed model (including their estimated uncertainty; i.e. a meta-analysis). Our second approach used an extension of the SnIPRE model (Eilertson, et al., 2012), which re-frames the MK test as a linear model in which polymorphism and substitution counts are predicted by synonymous or nonsynonymous state. Although this model does not explicitly consider the same underlying population-genetic processes, it does permit a straightforward extension to natively include gene, homologue, pathway, and host species as predictors in the model, and therefore provides a direct test of the questions of interest (although at a cost of potentially less accurate or arbitrarily-scaled parameter estimates). We have re-implemented the SnIPRE model using the Bayesian Generalised Linear Mixed Modelling R package MCMCglmm (Hadfield, 2010), and the code is provided in S1 text.

#### DFE-alpha analyses

DFE-alpha (Eyre-Walker & Keightley, 2009) infers ω_A_ (the number of adaptive nonsynonymous substitutions per nonsynonymous site, relative to the number of synonymous substitutions per synonymous site), while simultaneously modelling the distribution of deleterious fitness effects and population size changes (Keightley & Eyre-Walker, 2007; Eyre-Walker & Keightley, 2009). The ω_A_ statistic is closely related to the more widely reported α statistic (the proportion of nonsynonymous substitutions that are adaptive (Charlesworth, 1994; Fay, et al., 2001; Smith & Eyre-Walker, 2002; Bierne & Eyre-Walker, 2004; Welch, 2006), but differs in that ω_A_ is expected to be less dependent on effective population size and therefore better for cross-species comparisons (because the denominator, dS, should be less affected by the efficacy of selection, and thus effective population size (Gossmann, et al., 2010; Gossmann, et al., 2012; Kousathanas, et al., 2014). DFE-alpha utilises the observed site frequency spectrum (SFS) for putatively neutral synonymous sites and potentially selected nonsynonymous sites, and maximises the likelihood of observing these spectra given the distribution of deleterious fitness effects (DFE) for nonsynonymous sites and a step-change in effective population size (Eyre-Walker & Keightley, 2009). The ‘excess’ nonsynonymous divergence attributable to adaptive substitution is then inferred, given the maximum likelihood estimate of the DFE and the observed divergence (Eyre-Walker & Keightley, 2009). We inferred ω_A_ for: (i) each RNAi gene and each position-matched control gene (i.e. those with no known RNAi-pathway role falling within the same 200 Kbp interval); (ii) each RNAi subpathway and their set of control genes, and; (iii) all RNAi pathway genes together, by pooling polymorphism and divergence data across genes within classes. We then compared this grouped polymorphism and divergence data in pathways of interest against control genes. We estimated the parameters of the nominal change in population size (the relative population size change parameter N_2_, and the time of the population size change, t_2_) for all genes treated together within species, and then fixed these estimates for pathway and individual gene estimates. Conditional on this species-wide estimate of demographic history, the DFE was estimated separately for RNAi and control genes. We obtained confidence intervals for estimates of α and ω_A_ by bootstrapping genes within classes (1000 draws), and we tested for differences in rate between gene classes by randomly permuting genes 1000 times between classes. To test for differences in the DFE between RNAi and control genes we performed a likelihood ratio test between a model in which parameters of the DFE were estimated for all genes together, and one in which we allowed the DFE parameters to be estimated separately for RNAi and control genes.

Pooling polymorphism and divergence data across genes allows calculation of pathway-specific ω_A_ within a species, but cannot readily give cross-species estimates. Therefore, we also calculated ω_A_ for individual genes in each species, and analysed these estimates across species. In general, such estimates are extremely poor unless samples sizes are extremely large (e.g. hundreds of alleles are sampled, or genes are very large) (Keightley & Eyre-Walker, 2010). However, if the selective pressure acting on genes is consistent across species, for example as is assumed by many phylogenetic approaches to detecting selection (Yang, 2007), we can acquire more accurate estimates of the relative rate of adaptive evolution by combining information across species. We therefore used a formal meta-analytic approach to combine small-group and single-gene estimates across species using MCMCglmm (Hadfield, 2010), by constructing linear mixed models. These models were used to estimate average gene-level ω_A_ of various pathways and homologues, and variation among gene-level ω_A_ estimates.

The first three models took the same form, only distinguished by the pathways among which genes were divided. In Model 1A the genes were classified as either ‘control’ or ‘RNAi’, in Model 1B the RNAi class was expanded into four levels: ‘miRNA’, ‘siRNA’, ‘viRNA’, and ‘piRNA’ and in Model 1C the piRNA class was further split into three functional categories: ‘effectors of transcriptional silencing’, ‘effectors of post-transcriptional silencing’, and ‘biogenesis factors’. The model for the estimate of *ω*_A_ (i.e. 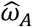) for homologue *k* in gene class *l* in species *m* had the form:
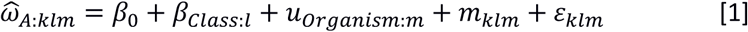

where *β*_0_ is the intercept, *β_Class_*_:*l*_ is a fixed effect associated with gene class *l*, *u_Organism_*_:*m*_ is a random effect associated with species *m*, *m_klm_* is the sampling error associated with each estimate, and *ε_klm_* is the between observation error after accounting for measurement error, which was allowed to vary by gene class (i.e. pathway). The variance of the sampling errors was obtained by bootstrapping genes by codon, and this sampling error variance was fixed at that value in the analysis. All species effects were assumed to come from a single normal distribution but the errors were assumed to come from independent normal distributions with different variances for each gene class.

Model 2 extended Model 1 by including homologue as a random effect (*u_Hom_*_:*kl*_) in order to identify homologues with elevated adaptation across lineages:
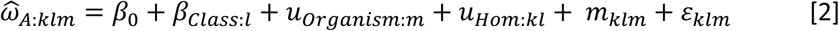

where the homologue effects were assumed to come from independent normal distributions with different variances for each gene class. In this model the cross-species average *ω*_A_ for a homologue *k* in gene class *l* is given by 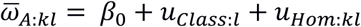. However, if genes are misclassified with respect to the gene class they belong, then 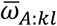 is likely to biased in general, and particularly so for misclassified genes. An arguably more conservative approach is to only use information from homologous genes to estimate the cross-species (i.e. remove the class effects from the model; this approach is provided as Model 2B in S1 text) and have 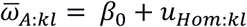. See S1 text for R code and a full description of the models used.

#### SnIPRE-like analysis

The meta-analytic approach to cross-species analysis above has the advantage of utilising DFE-alpha estimates that are inferred under an explicit population-genetic model. However, it has the disadvantage that it conditions on point estimates from a model, rather than using the available data directly. We have therefore taken advantage of the Poisson linear mixed model approach to MK analyses ‘SnIPRE’ proposed by Eilertson et al. (2012), which models the counts of mutations in four classes: synonymous within-species polymorphisms, nonsynonymous within-species polymorphisms, between-species synonymous differences (divergence) and between-species nonsynonymous differences. By fitting ‘nonsynonymous’ and ‘divergent’ as main effects, selection can be inferred from their interaction, which records the excess contribution of nonsynonymous mutations to between-species divergence. This excess can be assessed at the level of individual genes (by treating gene identity as a random effect) or can be expressed as a function of other fixed or random effects such as gene class and species. Although this approach does not directly provide parameter estimates that are interpretable in simple population-genetic terms, such as ω_A_, it has the advantage of extending naturally to provide comparisons between species and gene classes while still using raw count data directly. Here we combine polymorphism and divergence data from several species to test whether RNAi genes have higher rates of adaptive substitution than our set of control genes, whether these rates vary between different subclasses of RNAi gene, and whether these rates vary between different homologues. We fit these models with the R package MCMCglmm (Hadfield, 2010) and the code is provided in the S1 text. In their single-species and single-class analysis Eilertson et al. (2012) used the generalised linear mixed model with the fixed effect part of the model as:
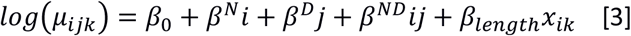

where *μ_ik_* is the expected number of mutations in gene *k* in one of the four classes indexed by *i* = 0,1 and *j* = 0,1 where *i* = 1 indicates nonsynonymous (*N*) and *k* = 1 divergent (*D*). This model estimates the intercept *β*_0_ (the density of synonymous polymorphisms), *β^N^* (the genome-wide difference between a mutation being nonsynonymous versus synonymous), *β^D^* (the genome-wide difference between a mutation being a substitution versus a polymorphism), and *β^ND^* (the interaction effect describing any genome-wide excess or dearth of nonsynonymous substitutions). *x_ik_* is the logarithm of the number of sites in gene *k* where a synonymous (*i* = 0) or a nonsynonymous (*i* = 1) mutation could occur and the fixed effect *β_length_* models how the number of observed mutations changes as a function of the number of sites. Eilertson et al. (2012) also fitted a random effect structure that models between-gene mutation patterns after accounting for the fixed effects:
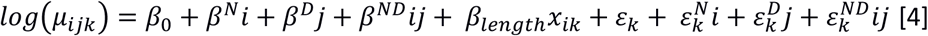

where the additional terms denoted *ε* are the gene-specific random deviations from each of the first four fixed effect terms described above. The four gene-specific random deviations were assumed to come from a multivariate normal distribution with estimated (co)variance matrix. Eilertson et al. (2012) define the selection effect of gene *k* as 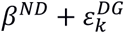, where a positive effect is evidence for positive selection, and (in Bayesian terms) the posterior probability that the effect exceeds zero can be directly assessed.

Here we extend the *SnIPRE*-like model of Eilertson et al. (2012) to accommodate multiple species and to allow the evolutionary parameters to differ between different classes of gene. To this end we allowed the four fixed effects to vary by species and by gene class (control, piRNA, siRNA, miRNA and viRNA) to give the fixed effect model:
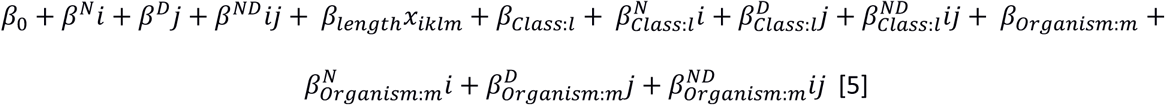

From this, we calculated the estimated selection effect for a specific pathway as 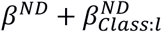. The random effect portion of the model included homologue-specific effects and gene-specific effects and had the form
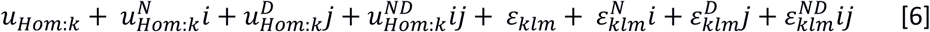

In addition to the four gene effects, the four homologue effects were also assumed to come from a multivariate normal distribution with estimated (co)variance matrix. We used this model to calculate the selection effect for homologue *k* in gene class *l* as 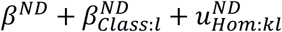 and each gene as 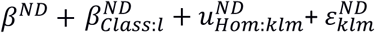. We estimated *β_length_* rather than fixing it at one, as in Eilertson et al. (2012), although the posterior mean of *β_length_* was close to one, supporting the assumption of Eilertson et al. (2012). In addition, we also fitted the SnIPRE model without assuming genes belong to known pathways, analogous to model 2. The code to fit these models is provided in the S1 text.

### Selective sweep analysis

The recent spread of a positively selected allele leaves characteristic patterns of diversity and allele frequencies in the genomic region surrounding the selected site, and these can be used to detect recent adaptive substitutions (e.g. Maynard Smith & Haigh, 1974; Barton, 1998; Nielsen, et al., 2005). We used SweeD (Pavlidis, et al., 2013; derived from Sweepfinder, Nielsen, et al., 2005) to search for evidence of recent selective sweeps in the regions surrounding RNAi genes. The algorithm scans the genome and at a user-defined interval calculates the composite likelihood of the observed site frequency spectrum (SFS) under a model of a selective sweep centred on that site, versus a standard neutral model. The ratio of the two composite likelihoods (CLR) is then used as a test statistic, with significance assessed by coalescent simulation (see Figure S1 and Text S2). We used this method to scan 200 kb (or less if the reference genome contig was less than 200 kb) surrounding each gene of interest in each species. For each focal region, we polarised the SFS by parsimony between the outgroup reference genome and the ingroup consensus sequence, which we aligned with LastZ ungapped alignment (Harris, 2007). We did not assume an ancestral state for fixed differences that were invariant in our ingroup (i.e. these sites were folded). This will make the analysis more robust to possible errors during contig alignment, because misalignment would manifest itself as regions of increased divergence between species. We included invariant sites in the analysis, as a characteristic signature of a recent sweep is a lack of diversity, and so including invariant sites in Sweepfinder analyses can greatly improve statistical power (Nielsen, et al., 2005). This comes with a risk of increased false positives (Huber et al, 2016), but including these sites should not differentially affect RNAi and control genes, unless there is a consistent difference in mutation rates between these two classes of genes. The SweeD analysis provides CLR values for equidistant points across the genome, with CLR values forming a “peak” in areas with high support for a sweep. To assess whether RNAi genes have experienced more sweeps than control genes in 6 of our 8 species (*B. mandarina* and *P. pacificus* were not tested because the published genome assemblies are unannotated), we counted the number of RNAi and control genes that overlapped significant peaks in the CLR statistic (based on the significance threshold provided by coalescent simulation, Figure S1, S2 text). If consecutive peaks occurred within 1 kb of each other, we classified them as a single broad peak, such that the contig was split into “sweep-positive” and “sweep-negative” areas. We then classified all genes along the contig as to whether they overlapped a “sweep-positive” area or not, and whether or not they were an RNAi gene. We used a binomial test to assess whether RNAi or control classes had more sweep-positive genes than expected given the summed gene length for each class.

To test whether sweeps were enriched in any particular subpathway, we normalised the maximum CLR statistic in a gene by the expected significance threshold from coalescent simulations and modelled these values 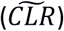 using the following linear mixed model:
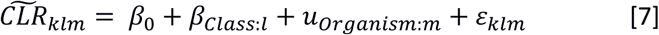

Here, *β_Class_*_:*l*_ is a fixed effect for the pathway each gene is assigned (miRNA, siRNA, piRNA or viRNA), *u_Organism:m_* is a random effect for species *m* and *ε_klm_* is the error term.

In the four organisms for which we have haplotype information (*D. melanogaster, D.pseudoobscura, P. pacificus, C. briggsae*), we additionally tested for ongoing or soft sweeps using the haplotype-based nSL statistic (Ferrer-Admetlla, et al., 2014). The nSL statistic is similar to the more widely used iHS statistic (Voight, et al., 2006), except that distance is measured in polymorphic sites rather than the genetic map distance (Ferrer-Admetlla, et al., 2014). This genome scan calculates the average number of consecutive polymorphisms associated with either the ancestral or derived allele at each polymorphic site along the contig across all pairwise comparisons. Areas with long range linkage disequilibrium will therefore be identified through SNPs with extreme nSL values.

## Results

### Evidence of genome-wide adaptive substitution in insects, but not nematodes

The position-matched ‘control’ genes (that lack RNAi-related function) included in our analyses allowed us to estimate the average genome-wide rate of adaptation, assuming that proximity to RNAi gene has no effect on their rate of adaptive evolution. Our analysis broadly agrees with previous ones, suggesting a substantial fraction of amino-acid substitution is adaptive across insect species (Figure 1). All insect species shared similar estimates (ω_A_ from 0.02 to 0.05) except for *D. pseudoobscura,* which exhibited an extremely high ω_A_ value of 0.16 [0.05,0.32] (95% bootstrap confidence interval) adaptive nonsynonymous substitutions per synonymous substitution per site. Although we only sampled two nematode lineages, it is notable that both ω_A_ estimates were negative (*C. briggsae*: -0.20 [-0.25, -0.15]; *P. pacificus:* -0.24 [-0.27, -0.21]. This is consistent with the previously noted high ratio of nonsynonymous to synonymous polymorphism (π_A_/π_S_) ratio in these species, and perhaps suggests population structure and local adaptation (Rödelsperger, et al., 2014; Thomas, et al., 2015). We also calculated α, or the proportion of adaptive substitutions for each species, which reflect the same patterns observed for ω_A_ (Figure S2).

**Figure 1:**
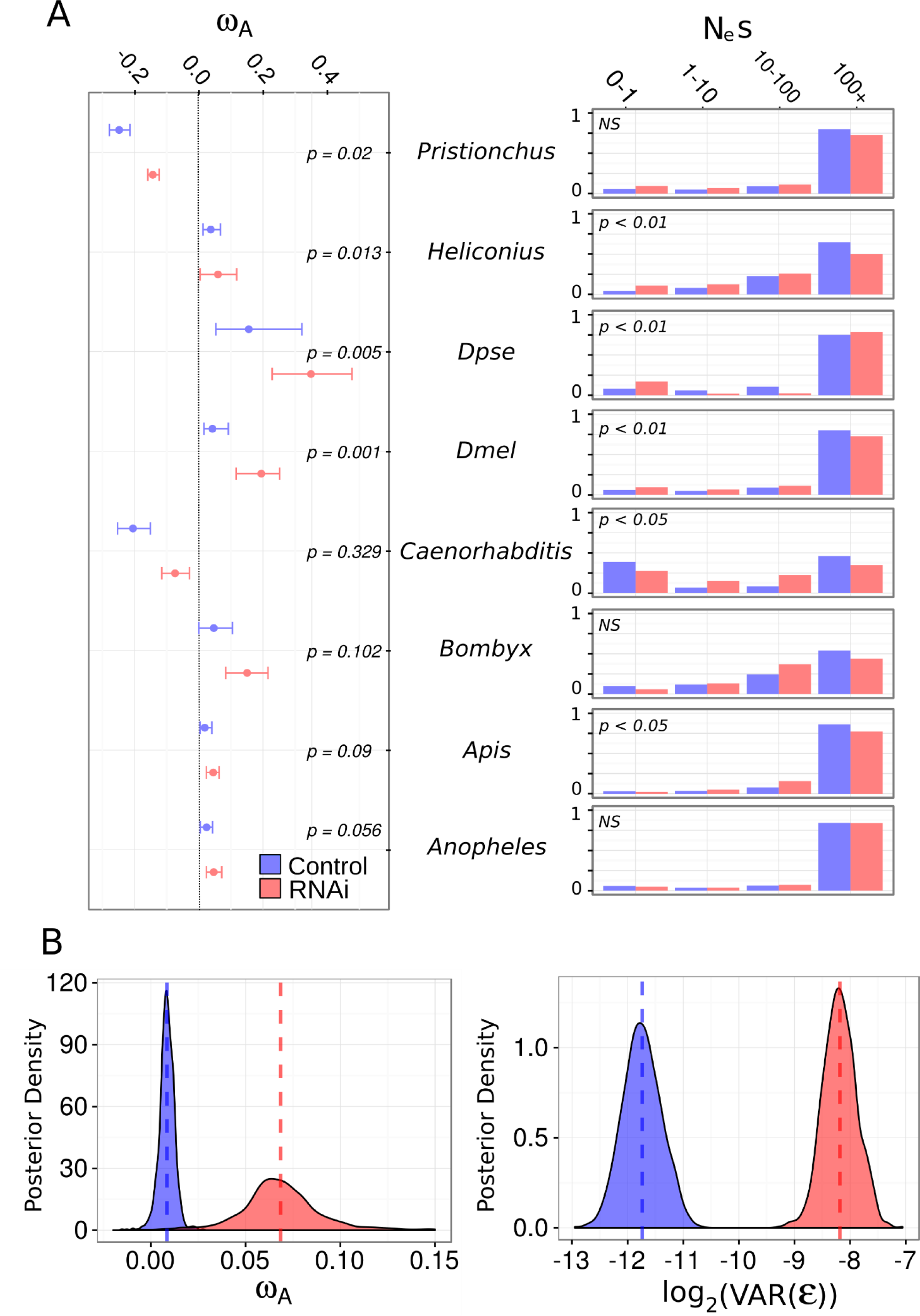
ω_A_ the DFE differ between RNAi genes and other genes. (A) Left: For each species, ω_A_ estimates and bootstrap confidence intervals for control (i.e. non-RNAi; blue) and RNAi (red) genes are plotted, with 95% bootstrap confidence intervals. Significance was determined by permutation. Right: The estimated discretised DFE for each species, with the proportion of mutation with *N_e_s* values in each category given for non-RNAi (blue) and RNAi (red) genes. (B) The posterior distribution of estimated ω_A_ for RNAi (red) versus control (blue) genes, showing that RNAi genes have much great ω_A_ estimates (left) and greater residual gene-level variation (right), indicating RNAi genes display higher rates adaptive amino-acid substitution, but are more variable.

The cross-species SnIPRE-like model provides a formal comparison of adaptive divergence in the insect species. The structure of the model forces comparison relative to one species, for which we chose *D. melanogaster*. *Anopheles gambiae* and *Bombyx mandarina* had levels of putatively adaptive nonsynonymous divergence that were indistinguishable from those of *D. melanogaster* (MCMCp = 0.489 and MCMCp=0.616, respectively). Consistent with the DFE-alpha estimates of ω_A_, *A. mellifera* and *H. melpomene* had significantly less adaptive nonsynonymous divergence than *D. melanogaster* (MCMCp = 0.04 and MCMCp < 3x10^-4^, respectively), whereas *D. pseudoobscura* had an increased excess of nonsynonymous divergence (MCMCp = 0.0005). Other species-specific SnIPRE parameters can be found in S1 text.

### RNAi genes consistently display more adaptive protein substitution than other genes

For each focal species we estimated the distribution of fitness effects of new mutations using DFE-alpha for RNAi pathway and non-RNAi (‘control’) genes, by pooling polymorphism and divergence data for each gene class. We fitted two models, one in which RNAi and control genes share a single DFE, and second in which each class of gene had a separate DFE. We then compared these models using a likelihood ratio test. In *D. melanogaster, D. pseudoobscura*, *H. melpomene, A. mellifera,* and *C. briggsae*, models in which control and RNAi genes have separate DFE parameters fitted the data significantly better than a model in which the two classes share a single DFE (Figure 1). Although there is no clear or universal trend, the DFE of control genes generally seemed slightly shifted towards more deleterious mutations than RNAi genes. For example, in most lineages (not *D. pseudoobscura* or *A. gambiae*), the estimated DFEs had a higher proportion of strongly deleterious mutations in control genes than RNAi genes, which suggests less constraint in RNAi genes. However, the overall shape of the DFE is quite different between species, either indicating that in these species gene function may play a smaller role than other factors in patterns of polymorphism, or that the DFE is estimated with low precision.

We then compared rates of adaptive amino acid substitution in RNAi genes to those in the non-RNAi control genes in each lineage, by pooling polymorphism and divergence data for the two classes as input to DFE-alpha (Figure 1). In every species tested, the point-estimate of class-wide ω_A_ was greater in RNAi genes than control genes. Although the effect was often small, the difference was individually significant in *D. melanogaster, D. pseudoobscura, H. melpomene,* and *P. pacificus.* To quantify the overall difference, we analysed individual gene estimates of ω_A_ in a linear mixed model framework (i.e. a meta-analysis) to estimate across-species rates of adaptive evolution in control and RNAi genes (model 1 in S1 text, Figure 1). We found the cross-species ω_A_ was significantly greater for RNAi genes than control genes, estimated as ω_A_ = 0.062 [0.049, 0.078] (95% Highest posterior density) versus ω_A_ = 0.01 [0.0009, 0.019] (*p* < 0.001). In addition, the residual gene-level variance was also much greater (MCMCp <0.001) for RNAi genes (0.0037, [0.0022, 0.0051]) than control genes (0.0003, [0.0001, 0.0004]), implying that ω_A_ is more variable in this class than among genes in general and consistent with a subset of RNAi genes or pathways undergoing extreme rates of adaptive amino acid substitution (Figure 1).

### Adaptive rates are high in piRNA and viRNA pathways

The higher rate of adaptive substitution seen in RNAi genes as a whole could result from slightly elevated positive selection across all components, or to a subset of the genes or pathway being substantially elevated. The higher gene-level variance seen in RNAi genes (above) suggests the latter, and to test this we pooled polymorphism and divergence data by sub-pathway for each insect species to calculate rates of adaptation in miRNA, siRNA, viRNA (i.e. confirmed antiviral siRNA in *D. melanogaster*), and piRNA pathways (Figure 2). In each species, the piRNA pathway exhibited a significantly greater rate of adaptive amino acid substitution than control genes, and miRNA pathway genes showed similar rates to control genes. Rates of adaptation for the siRNA (both endo-siRNA and viRNA) pathway were greater in only a subset of lineages. The magnitude of rates and proportion of adaptive lineages increased upon removing endo-siRNA genes and restricting the analysis to viRNA genes only. For all subsequent analyses, we analysed these pathways separately to test the hypothesis that the core antiviral RNAi genes have elevated rates of adaptive evolution.

**Figure 2:**
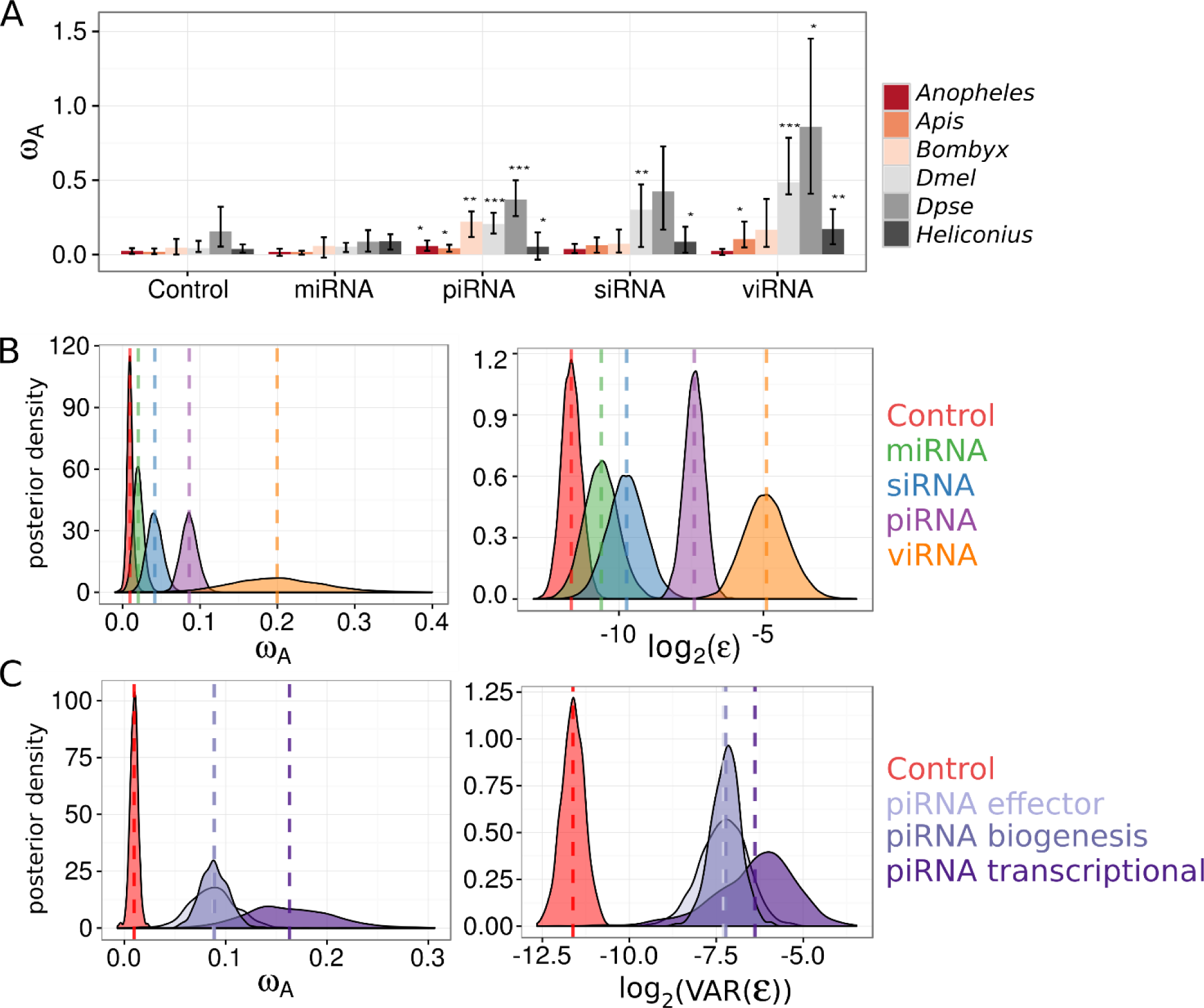
DFE-alpha estimates of ω_A_ for each subpathway. (A) ω_A_ estimates from pooled polymorphism and divergence data across insect RNAi subpathways using DFE-alpha. ω_A_ was estimated for each subpathway in each organism and confidence intervals obtained by bootstrapping across genes. Significance was assessed by permutation tests between sub-pathway and control genes for each organism (p < 0.05*, p<0.01**, p<0.001***). (B) Individual-gene DFE-alpha ω_A_ estimates were analysed using a linear mixed model in MCMCglmm, and show that (left) the viRNA pathway exhibits the fastest rate of adaptive protein substitution, followed by the piRNA pathway, and that among-gene variance shows the same pattern (right). (C) Individual gene DFE-alpha ω_A_ estimates were analysed in MCMCglmm, except that the piRNA pathway was further split into genes involved in transcriptional silencing, piRNA biogenesis, or piRNA-mediated effectors of silencing. The posterior distributions of these three effect sizes versus control genes are plotted. All three piRNA functions are targets of elevated positive selection and have large residual variances, although genes mediating transcriptional silencing have greater point estimates for both.

To formalise the effect of pathway (miRNA, piRNA, non-antiviral endo-siRNA, viRNA) while accounting for variability in adaptation across species (model 2 in S1 text, Figure 2), we performed a meta-analysis of ω_A_ estimates in individual genes from DFE-alpha, fitting pathway as a fixed effect. The piRNA, viRNA, and endo-siRNA pathways were each significantly different from control genes (control ω_A_ =0.01 [0.002,0.018]; piRNA MCMCp < 0.001; viRNA MCMCp = 0.002; siRNA MCMCp = 0.004; for MCMCp value calculation, see the S1 text), with cross-species estimates of ω_A_ of 0.08 [0.06,0.10], 0.18 [0.06, 0.30] and 0.03 [0.01,0.05], respectively. The viRNA pathway ω_A_ estimate was not significantly greater than the piRNA pathway (MCMCp = 0.07), but was greater than the endo-siRNA pathway (MCMCp = 0.01), and the miRNA pathway (MCMCp < 0.001). The ω_A_ estimate for the piRNA pathway was significantly greater than the endo-siRNA (MCMCp = 0.002) and the miRNA pathways (MCMCp < 0.001). Consistent with our analysis of pooled polymorphism and divergence data, the rate of adaptive evolution in the miRNA pathway (ω_A_ = 0.01 [-0.001, 0.02]; MCMCp=0.09) was not significantly different from control genes. Our linear models included pathway-specific error variances, which were lower for control genes (3 [2,4] x10^-4^) and miRNA pathway genes (7 [2,12] x10^-4^) than for endo-siRNA (13 [4,22] x10^-4^), piRNA (66 [37,97] x10^-4^), and viRNA pathway genes (0.04 [0.007, 0.86]), consistent with a great variation in adaptive rates in these pathways.

We repeated the subpathway-level analysis using a SnIPRE-like model (Eilertson, et al., 2012) to estimate the average selection effect within subpathways across organisms, without making any explicit assumptions about the DFE. Although SnIPRE can be used to provide estimates of population genetic parameters, we limit our discussion to the “selection effect” statistic, where negative values are consistent with constraint and positive values with adaptive protein evolution, and magnitude reflects the strength of positive or negative selection. Consistent with our analysis of DFE-alpha estimates, the SnIPRE model identified a mean positive selective effect estimated across species (selective effect=0.25 [0.02, 0.46] 95% HPD interval, MCMCp = 0.03), with large variance among genes (Figure 3). Again, viRNA, endo-siRNA, and piRNA pathway-level selection effects were significantly elevated compared to control genes (viRNA: 1.10 [0.63, 1.57] MCMCp < 5x10^-4^, non-antiviral siRNA: 0.96 [0.44, 1.52] MCMCp = 0.02, piRNA: 0.63 [0.44, 0.84] MCMCp < 3x10^-4^), with the viRNA pathway exhibiting a significantly larger effect than the piRNA (MCMCp = 0.006), but not the endo-siRNA (MCMCp = 0.66). In agreement with the DFE alpha analysis, the miRNA pathway was not significantly different from control genes (MCMCp = 0.07), and had a selection effect of 0.53 [0.20, 0.86].

**Figure 3.**
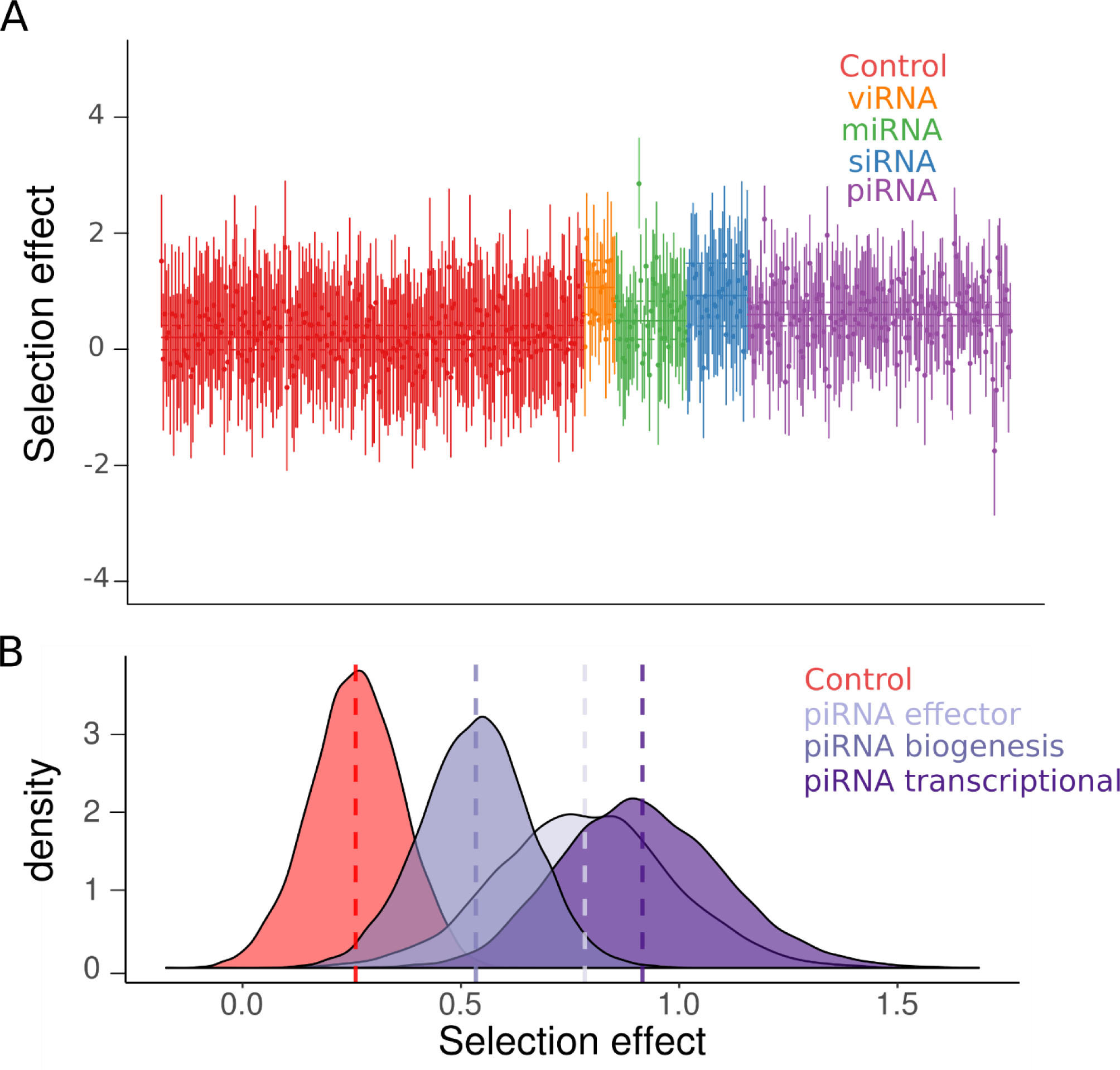
SnIPRE-like selection effects. (A) SnIPRE ‘selection effect’ with 95% confidence intervals (species-level effects removed) are plotted for each gene in each species, coloured according to the gene’s role in the RNAi pathway. Solid horizontal lines signify the mean selection effect for each RNAi subpathway (or control genes) with dotted lines signifying the 95% confidence intervals for the subpathway mean. SnIPRE and DFE-alpha analyses are consistent in suggesting that the viRNA, endo-siRNA, and piRNA pathway have more adaptive amino-acid substitutions than control genes. (B) We also performed a SnIPRE analysis after dividing the piRNA pathway into three functional classes, as in Figure 2. The posterior distribution of selection effects associated with each piRNA function are plotted. Similar to DFE-alpha, SnIPRE identifies all three pathways as significantly elevated relative to control genes, however in the SnIPRE analysis transcriptional silencing genes have a significantly greater adaptive rate than biogenesis factors.

### Adaptation is elevated in all major piRNA pathway functions, but is most enriched in transcriptional silencing

Rapid adaptation in *Drosophila* piRNA pathway genes has been hypothesized to be the result of fluctuating selection for increased TE defence and decreased off-target genic silencing (Blumenstiel et al 2016). A prediction of this hypothesis is that genes involved in transcriptional silencing would be under increased positive selection. We tested this prediction by further dividing the piRNA pathway into effectors (e.g. PIWIs), biogenesis factors (e.g. adapter proteins), and transcriptional silencing factors, and using single-gene polymorphism and divergence data to estimate ω_A_ and the selection effect for each piRNA functional category (Model 3). We found all piRNA functional groups are significantly greater than control genes (MCMCp < 0.001) (Figure 2C), and that transcriptional silencing genes (ω_A_ = 0.16 [0.08-0.25]) have greater adaptive rates than effectors (MCMCp = 0.04, ω_A_ = 0.08 [0.04-0.13]) and biogenesis factors (MCMCp = 0.03, ω_A_ = 0.08 [0.05-0.11]). This result holds when excluding *Drosophila* transcriptional silencing factors *rhino*, *deadlock*, and *cutoff*, which are products of recent gene duplication or *de novo* formation (Figure S3), and may not have evolutionary rates that are directly comparable to other genes.

We also estimated the average selection effect for each functional process of the piRNA pathway using the SnIPRE approach. Similar to the DFE-alpha meta-analysis, we find that all piRNA functional categories have elevated positive selection relative to control genes (biogenesis: MCMCp=0.018, effector: MCMCp=0.012, transcriptional silencing: MCMCp=0.0004), that transcriptional silencing factors had the largest average selection effect of 0.92 [0.58, 1.31], and that genes involved in transcriptional silencing were significantly greater than biogenesis factors (selection effect: 0.53, [0.29, 0.78], MCMCp = 0.027) (Figure 3B). In contrast to the DFE-alpha meta-analysis, however, genes involved in transcriptional silencing were not significantly greater than effector genes (0.78 [0.40, 1.19], MCMCp = 0.68), and pathway-level point estimates of these selection effects were much closer (Figure 2C, Figure 3B).

### Individual genes in the piRNA and viRNA pathway show elevated adaptation

The higher overall rates of adaptive protein substitution seen in RNAi genes may result from the engagement of some genes in an evolutionary arms race (e.g. with viral suppressors of RNAi), a response to the selection imposed by the invasion of novel parasites (e.g. transposable elements), or a trade-off between the specificity and sensitivity of genome defense (Aravin, et al., 2007; Obbard, et al., 2006; Blumenstiel, et al., 2016). We used a linear mixed model to combine single-gene estimates of ω_A_ from DFE-alpha across multiple species to identify candidate arms race genes in the RNAi pathways, fitting subpathway as a fixed effect, with homologue and organism as random effects, and subpathway-specific error variances. We found little variation among genes in a subpathway after accounting for subpathway, and in most cases there was not enough information to differentiate individual genes from the pathway mean (Figure 4A). Although a model that accounts for pathway is statistically preferable if pathways are meaningful, any errors in assigning ‘pathway’ membership would introduce bias to the estimates for misclassified genes. We therefore also estimated homologue-specific effects in a model that excludes the subpathway effect (model 2B in the S1 text). This model finds significant evidence for positive selection in fewer genes (Figure S4A) including 13 of 22 piRNA genes, 2 of 3 viRNA genes, and no genes in the siRNA or miRNA pathway.

**Figure 4.**
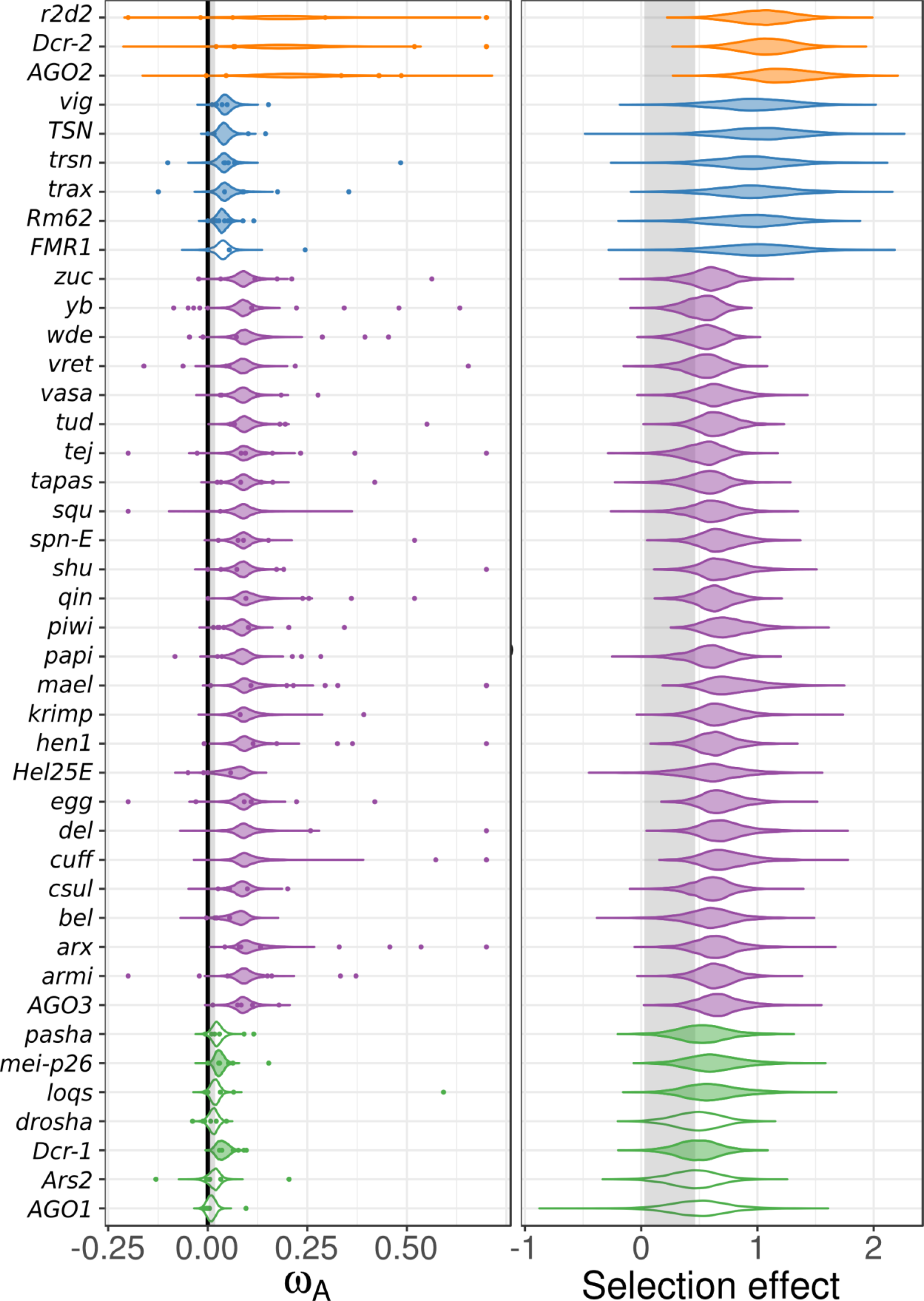
Cross-species homologue-level estimates of ω_A_ and selection effects. (Left) Individual homologue ω_A_ estimates (coloured points) were calculated using DFE-alpha and analysed using a linear mixed model with subpathway as fixed effect and species and homologue as a random effect (estimate uncertainty was included by incorporating bootstrap intervals as measurement error variance). The posterior distributions of the cross-species estimate for ω_A_ for each homologue are plotted, and shaded if significantly different from the control gene distribution (region shaded grey). Single-gene estimates of ω_A_ > 0.75 are plotted at 0.75 for clarity. (Right) The analogous analysis performed using SnIPRE, with the posterior distribution of homologue-level selection effects plotted. Both analyses find little variation among homologues after accounting for subpathway, and homologue-level analyses generally mirror pathway-specific analyses. See Figure S4 for the equivalent models that exclude the fixed effect of pathway.

We also performed this homologue-level analyses using the SnIPRE approach. Similar to the DFE-alpha meta-analysis, we found very little information after accounting for subpathway (Figure 4B), resulting in low among-gene variation within RNAi subpathways. When we excluded subpathway effects, we found a similar result to the homologue-level DFE-alpha meta-analysis without subpathway, except fewer piRNA pathway genes are nominally significant (6/22 genes). Notably, *maelstrom, eggless, piwi* (including *aub*), *AGO2,* and *Dcr-2* were found to have significantly elevated positive selection across all four homologue-level analyses (i.e. with or without imposing a subpathway classification).

MK tests are commonly used to test for positive selection in individual genes. SnIPRE selection effects can be used to perform an analogous test for selection, except the approach can gain power by taking in the genome-wide distribution of polymorphism and divergence patterns by fitting gene as a random effect (Eilertson et al, 2010). We find that 36% of RNAi genes show nominally ‘significant’ positive selection. In contrast, only 5% of selection effects in control genes are significantly positive (Table S5). At the pathway level, 40% of piRNA genes, 44% of viRNA genes, 26% of non-antiviral siRNA pathway genes, and 25% of miRNA pathway genes have significantly positive selection effects (Table S5). No gene had positive selection effects in every lineage, although *armitage, capsuleen, cutoff, tudor, vasa, vretano,* and *Yb* homologs were identified in over half the lineages.

### Selective sweeps are detectable across functional classes of RNAi genes

Recent positive selection is expected to leave a characteristic mark in the genome, including a SFS skewed towards low and high frequency alleles and a local reduction in polymorphism (Maynard Smith & Haigh, 1974; Barton, 1998; Nielsen, et al., 2005). As RNAi genes show elevated rates of adaptive evolution, we speculated that they may also exhibit more evidence of recent selective sweeps. Using SweeD, we found that many of the insect lineages do show evidence for sweeps in a subset of RNAi genes (Figure 5, Figures S5-S12). We tested whether RNAi genes have undergone more recent sweeps than surrounding genes by classifying nominally significant peaks as either occurring near (within 1 KB) an RNAi gene or not, and using a binomial test to determine whether more sweeps than expected occur in RNAi genes (given their length). In four of the six species tested (*D. melanogaster, D. pseudoobscura, A. mellifera,* and *A. gambiae*) there were significantly more detectable sweep signals in RNAi genes than in surrounding non-RNAi genes (*D. melanogaster p* = 0.0006; *A. mellifera p* = 0.015*; A. gambiae p* = 0.0001*; D. pseudoobscura p* = 7x10^-5^). However, we find no difference among subpathways in the frequency with which we detected recent sweeps. In addition, none of the genes exhibited a significant CLR peak across all organisms tested, although *spn-E* and *vig* display significant evidence of recent sweeps in five of the six insect lineages. It was notable that 34% of the variation in the per-gene maximum CLR test statistic was attributable to species, consistent with either sample size or demographic history playing a substantial role in our power to detect sweeps.

**Figure 5.**
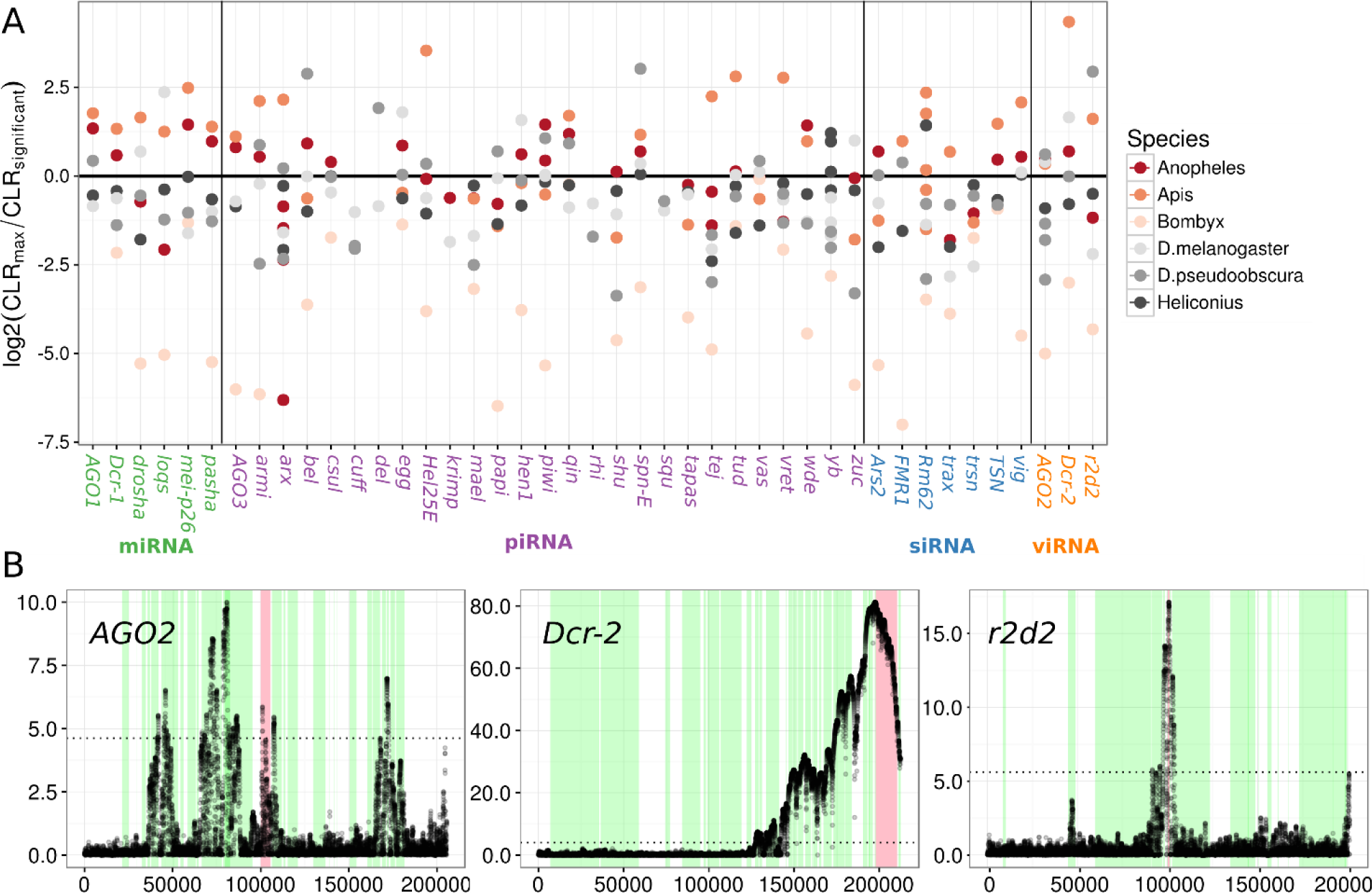
Selective sweeps in RNAi genes and example SweeD plots. (A) Points indicate the log_2_ ratio of the maximum observed CLR value (from SweeD) in the named gene to the CLR 95% significance threshold inferred from simulation. Values above 0 indicate there was a ‘significant’ CLR peak in a genic region and colours indicate species. (B) The viRNA pathway in *Apis mellifera* shows strong evidence for recent sweeps. For each of the three viRNA pathway genes the CLR statistic is plotted across a 200 kb region. The dotted line is the significance threshold estimated through neutral simulations under a published demographic history. Red regions denote the focal gene and green regions highlight surrounding genes. In Apis, both *Dcr2* and *R2D2* show strong evidence for sweeps with the surrounding region of *Dcr2* being devoid of polymorphism, indicating this sweep was recent and rapid. *AGO2* also shows a significant peak, but this is narrow and only marginally significant.

Sweep signatures were the most pronounced in *A. mellifera*, in both the CLR magnitude and breadth of the genomic region affected (Figure 5, Figure S10). These were associated with large regions devoid of any polymorphism, despite the high rate of recombination seen in honeybees (Beye, et al., 2006), which is expected to narrow the region affected by a nearby sweep. We also searched for evidence of haplotype structure, as would be expected during an ongoing or soft selective sweeps using the nSL statistic (data not shown). However, there were no strong signals in any of the RNAi genes for which we had haplotype information.

## Discussion

Using both DFE-alpha and SnIPRE-like McDonald-Kreitman framework analyses we identify elevated rates of adaptive evolution in RNAi-pathway genes across six insects and two nematodes. In most species, the RNAi-pathway genes are also more likely to display evidence of a recent selective sweep. These results generalise the findings of previous analyses in *Drosophila*, and are consistent with these genes being engaged in an arms race across the invertebrates. Across species, we find that genes involved in the suppression of viruses and transposable elements show the highest rates of adaptive evolution, and those in the miRNA pathway the lowest (not significantly different from non-RNAi-genes). There is substantial variation in rates among RNAi but the antiviral genes *AGO2* and *Dcr-2* and the piwi-pathway genes *maelstrom, eggless, piwi*, *aub, armitage, capsuleen, cutoff, tudor, vasa, vretano, spn-E, vig* and *Yb* show consistently strong signatures of long-term and/or recent positive selection.

### Identification of rapidly evolving pathways by DFE-alpha and SnIPRE

Estimating rates of adaptive protein evolution in an MK-framework (McDonald and Kreitman, 1991) can be biased by past population size changes and slightly deleterious mutations that segregate at low frequencies. We compare adaptive rates between different classes of RNAi genes, accounting for these biases by explicitly modelling the DFE and demographic history using DFE-alpha (Eyre-Walker and Keightley, 2009), or by modelling the genome-wide patterns of polymorphism and divergence with SnIPRE (Eilertson, et al., 2012). Most of the qualitative results of each of these analyses are in agreement, however, SnIPRE and DFE-alpha analyses disagree on the relative differences in the rate of adaptive evolution among subpathways. For example, the DFE-alpha meta-analysis provides low point estimates for the endo-siRNA and miRNA pathways relative to the piRNA and viRNA, but SnIPRE identifies the endo-siRNA selection effect as higher than the piRNA, and piRNA genes closer to the miRNA. This incongruence could reflect differences in the DFE between subpathways. For example, genes in the miRNA and endo-siRNA pathways are highly conserved and have low rates of protein evolution, while mechanisms of piRNA pathway function are surprisingly diverse across animals (e.g. Morazzani, et al., 2012; Sarkies, et al., 2015). These differences in constraint could lead to an underestimation of miRNA and endo-siRNA pathway adaptation and overestimation of piRNA adaptation in the DFE-alpha analyses, and indicate that estimating the DFE separately for each subpathway may improve estimates.

### Adaptive protein evolution across species is enriched in specific functional pathways

We found large differences in rates of adaptative protein substitution between insects and nematodes, but less variation among insect species. In an analysis of variance, we find that species explained only 11% of the variation in gene-level estimates of ω_A_. In contrast, gene and pathway explained 42% of the variation in gene-level ω_A_ estimates. The elevated rate and among-gene variation seen in piRNA and viRNA pathway genes across species could be caused by rapid adaptation in the same subset of genes in a pathway, or in a random selection of genes in a pathway. Homologue-level analysis of ω_A_ and selection effects (Figure 4, Figure S4) indicates it is probably both, as subsets of homologues within pathways show consistent evidence for elevated adaptive protein evolution, but homologous genes also exhibit high variances across species.

### Potential Drivers of Adaptation in the viRNA pathway

It seems likely that the elevated rates of adaptive protein evolution we detect in the viRNA and piRNA pathways are a result of recurrent selection mediated by viruses and/or TEs. First, it is well established that defensive pathways show high rates of adaptive evolution, presumably as a consequence of antagonistic coevolution with parasites (Stenseth & Maynard Smith, 1984; Buckling & Rainey, 2002; Paterson, et al., 2010; Brockhurst, et al., 2014). For example, a recent analysis of virus-interacting proteins estimated that 30% of adaptive protein changes in mammals are driven by viruses (Enard, et al., 2016). Second, for the viRNA pathway genes at least, viral suppressors of RNAi are strong candidates to be the driving agent. Many RNA and DNA viruses of invertebrates are known to have proteins or structural RNAs which actively block RNAi function (Li, et al., 2002; Van Rij, et al., 2006; Nayak, et al., 2010; van Mierlo, et al., 2012; Bronkhorst, et al., 2014), and these can evolve rapidly and can be highly host-specific, consistent with an arms-race scenario (van Mierlo, et al., 2014). We find that *AGO2* and *Dcr-2* display consistently elevated rates of adaptive protein substitution across insect species, with additional limited evidence of elevated adaptation in *hen1*, all of which have previously been identified as targets of active suppression by viral proteins (viral suppressors of RNAi; VSRs) (Van Rij, et al., 2006; Vogler, et al., 2007; Nayak, et al., 2010; van Mierlo, et al., 2012; van Cleef, et al., 2014), lending credibility to the hypothesis that viruses may play a major role in driving the observed rapid evolution in RNAi genes.

### Potential Drivers of Adaptation in the piRNA pathway

Whereas an arms-race between antiviral RNAi genes and viral suppressors of RNAi is intuitive, the observed rapid adaptive evolution of piRNA pathway genes is currently harder to explain. Similar to viruses, TEs are costly for their hosts and could in principle select for increased suppression (Charlesworth, et al., 1994). However, piRNA-generating clusters ostensibly provide an adaptive defence that can arise on much faster time scales than fixation of advantageous mutations, reminiscent of acquired immunity (Brennecke, et al., 2007; Khurana, et al., 2011; Mohn, et al., 2015; Han, et al., 2015). The adaptive response in piRNA genes could be mediated by at least three non-exclusive mechanisms: (i) direct piRNA pathway suppression by TEs or by off-target VSRs, (ii) recurrent “retuning” of piRNA machinery after a novel TE invasion (Lee and Langley et al, 2012; Yi et al, 2014), or (iii) fluctuating selection on the sensitivity to detect transposon sequences and specificity to exclude off-target genic silencing (i.e. the “genomic auto-immune hypothesis”) (Blumenstiel, et al., 2016). Besides the global de-repression of transposons upon invasion of the Penelope retroelement in *D. virilis* (Petrov, et al., 1995; Evgen’ev, et al., 1997; Rozhkov, et al., 2010; Blumenstiel, et al., 2016), there is limited evidence for (i), and the mechanism underlying this phenomenon still awaits elucidation. The latter two hypotheses are not mutually exclusive, and both posit that piRNA adaptation occurs in response to recurrent horizontal transfer of new TEs into the genome, a common occurrence in insects (Peccoud, et al., 2017). In (ii), the piRNA pathway evolves to optimise defence against the current suite of transposons, becoming “less adapted” for dealing with historic, obsolete ones. This would result in a Red Queen-like scenario, but instead of antagonistic coevolution with one parasite, the piRNA pathway must defend against a constant recycling of TE lineages. As the germline cells face a higher TE diversity than somatic tissues, this is broadly supported by our observation that piRNA pathway genes with primarily germline function (Handler, et al., 2013; Czech, et al., 2013; Muerdter, et al., 2013) have higher rates of adaptive protein evolution than those functioning in the somatic layer of cells surrounding the *Drosophila* ovary (Figure S13),. The genomic autoimmunity hypothesis (iii) goes further, and proposes piRNA pathway adaptation to TE invasions results in increased piRNA function and associated off-target genic effects, which are then selected against after the TE is supressed (Blumenstiel, et al., 2016). It could be argued that our analysis of adaptive rates in piRNA functions lends broad support for this, in that genes mediating transcriptional silencing show the greatest adaptive rates across species in the piRNA pathway, with additional evidence for rapid adaptation in biogenesis factors, whose rates are expected to be correlated with the transcriptional machinery (Blumenstiel, et al., 2016). However, our pathway-level and homologue-level analyses also find signals of elevated adaptation in effector genes, which have rates that covary to a lesser degree with other piRNA factors (Blumenstiel, et al., 2016). This does not refute the genomic autoimmunity hypothesis, but may suggest additional selective forces acting on the piRNA pathway independent of genes underlying a trade-off between sensitivity and specificity. Nevertheless, our results would also fit within the context of (ii), in a scenario where the transcriptional machinery has a greater evolutionary potential than the rest of the piRNA pathway.

## Acknowledgements

We thank the authors of the previous studies who made their data publicly available. We thank Rob Ness and Peter Keightley for helpful advice on the analysis.

### Funding

WHP is supported by the Darwin Trust of Edinburgh and this work in DJO’s laboratory was partly supported by a Wellcome Trust strategic award to the Centre for Immunity, Infection and Evolution (WT095831). JDH is supported by a Royal Society Research Fellowship (UF150696).

---

**Table S1:** Insect genes analysed, FlyBase identifiers, and subpathway involvement

| Gene | Flybase ID | Pathway | References |
| --- | --- | --- | --- |
| AGO1 | FBgn0262739 | miRNA | Okamura et al, 2004 |
| AGO2 | FBgn0087035 | viRNA | Wang et al, 2006; van Rij et al, 2006 |
| AGO3 | FBgn0250816 | piRNA - effector | Brennecke et al, 2007; Gunawardane et al, 2007 |
| armi | FBgn0041164 | piRNA - biogenesis | Saito et al, 2010 |
| Ars2 | FBgn0033062 | miRNA | Gruber et al, 2009 |
| arx | FBgn0036826 | piRNA - effector | Ohtani et al, 2013; Donertas et al, 2013 |
| bel | FBgn0263231 | piRNA | Lo et al, 2016 |
| csul | FBgn0015925 | piRNA - effector | Kirino et al, 2009 |
| cuff | FBgn0260932 | piRNA - transcriptional | Pane et al, 2011; Mohn et al, 2014 |
| Dcr-1 | FBgn0039016 | miRNA | Jiang et al, 2005; Saito et al, 2005 |
| Dcr-2 | FBgn0034246 | viRNA | Wang et al, 2006; Galiana-Arnoux et al, 2006 |
| del | FBgn0086251 | piRNA - transcriptional | Mohn et al, 2014 |
| drosha | FBgn0026722 | miRNA | Lee et al, 2003 |
| egg | FBgn0086908 | piRNA - transcriptional | Rangan et al, 2011 |
| FMR1 | FBgn0086908 | siRNA | Ishizuka et al, 2002 |
| Hel25E | FBgn0014189 | piRNA - biogenesis | Zhang et al, 2012 |
| hen1 | FBgn0033686 | piRNA - biogenesis | Saito et al, 2007; Horwich et al, 2007 |
| krimp | FBgn0034098 | piRNA | Sato et al, 2015 |
| loqs | FBgn0032515 | miRNA | Jiang et al, 2005; Saito et al, 2005 |
| mael | FBgn0016034 | piRNA - transcriptional | Sienski et al, 2012 |
| Mei-p26 | FBgn0026206 | miRNA | Neumuller et al, 2008 |
| papi | FBgn0031401 | piRNA - biogenesis | Saxe et al, 2013 |
| pasha | FBgn0039861 | miRNA | Yeom et al, 2006 |
| piwi | FBgn0004872/FBgn0000146 | piRNA - effector | Brennecke et al, 2007 |
| qin | FBgn0263974 | piRNA - biogenesis | Zhang et al, 2011 |
| r2d2 | FBgn0031951 | viRNA | Liu et al, 2003 |
| rhi | FBgn0004400 | piRNA - transcriptional | Klatenhoff et al, 2009; Mohn et al, 2014 |
| rm62 | FBgn0003261 | siRNA | Csink et al, 1994 |
| shu | FBgn0003401 | piRNA - biogenesis | Olivieri et al, 2012; Preall et al, 2012 |
| spn-E | FBgn0003483 | piRNA - biogenesis | Malone et al, 2009 |
| squ | FBgn0267347 | piRNA - effector | Haase et al, 2010 |
| tapas | FBgn0027529 | piRNA - biogenesis | Patil et al, 2014 |
| tejas | FBgn0033921 | piRNA - biogenesis | Patil et al, 2010 |
| trax | FBgn0038327 | siRNA | Liu et al, 2009 |
| trsn | FBgn0033528 | siRNA | Liu et al, 2009 |
| TSN | FBgn0035121 | siRNA | Caudy et al, 2003 |
| tudor | FBgn0003891 | piRNA - biogenesis | Nishida et al, 2009 |
| vasa | FBgn0283442 | piRNA - biogenesis | Xiol et al, 2014 |
| vig | FBgn0024183 | siRNA | Caudy et al, 2002 |
| vret | FBgn0263143 | piRNA - biogenesis | Zamparini et al, 2011 |
| wde | FBgn0027499 | piRNA - transcriptional | Koch et al, 2009 |
| Yb | FBgn0000928,FBgn0051755,FBgn0037205 | piRNA - biogenesis | Saito et al, 2010 |
| zuc | FBgn0261266 | piRNA - biogenesis | Mohn et al, 2015; Han et al, 2015 |

**Table S2.** Nematode Genes and WormBase identifiers

| Gene | Wormbase ID | Gene | Wormbase ID |
| --- | --- | --- | --- |
| ACR-11 | WBGene00000050 | WAGO-4 | WBGene00010263 |
| AIN-2 | WBGene00015007 | LAM-2 | WBGene00016913 |
| C04F12.1 | WBGene00007297 | C35E7.8 | WBGene00016460 |
| CDC-25.1 | WBGene00000386 | EGO-1 | WBGene00001214 |
| DRH-3 | WBGene00008400 | EKL-1 | WBGene00009052 |
| DRSH-1 | WBGene00009163 | PPW-1 | WBGene00004093 |
| ERI-6 | WBGene00016561 | PPW-2 | WBGene00004094 |
| ERI-7 | WBGene00016566 | PRG-1 | WBGene00004178 |
| F57C9.7 | WBGene00019013 | PRG-2 | WBGene00004179 |
| GEI-11 | WBGene00001568 | RDE-2 | WBGene00004324 |
| HAF-6 | WBGene00001816 | SAGO-1 | WBGene00019666 |
| HPO-24 | WBGene00011945 | SAGO-2 | WBGene00018921 |
| MEL-26 | WBGene00003209 | SMG-2 | WBGene00004880 |
| MUT-16 | WBGene00003508 | WAGO-1 | WBGene00011061 |
| MUT-2 | WBGene00003499 | WAGO-2 | WBGene00018862 |
| PTR-2 | WBGene00004217 | WAGO-5 | WBGene00022877 |
| RDE-10 | WBGene00021634 | Y23H5A.3 | WBGene00021270 |
| RRF-1 | WBGene00004508 | ERGO-1 | WBGene00019971 |
| RRF-2 | WBGene00004509 | NRDE-2 | WBGene00011333 |
| RSD-6 | WBGene00004684 | PHO-1 | WBGene00004020 |
| ULP-5 | WBGene00006740 | PID-1 | WBGene00017549 |
| W01A8.5 | WBGene00012167 | PIR-1 | WBGene00011967 |
| PASH-1 | WBGene00011908 | RRF-3 | WBGene00004510 |
| MEL-47 | WBGene00017132 | VIG-1 | WBGene00006924 |
| TSN-1 | WBGene00006626 | WAGO-11 | WBGene00021711 |
| XRN-2 | WBGene00006964 | ALG-3 | WBGene00011910 |
| CUL-2 | WBGene00000837 | ALG-4 | WBGene00006449 |
| DCR-1 | WBGene00000939 | CGH-1 | WBGene00000479 |
| ERI-9 | WBGene00016143 | MUT-7 | WBGene00003504 |
| HENN-1 | WBGene00015349 | TBP-1 | WBGene00006542 |
| HPL-2 | WBGene00001996 | Y37D8A.16 | WBGene00012554 |
| NHL-2 | WBGene00003598 | COGC-2 | WBGene00000585 |
| NRDE-1 | WBGene00007577 | ERI-3 | WBGene00021103 |
| RDE-4 | WBGene00004326 | RDE-11 | WBGene00023421 |
| SID-2 | WBGene00004796 | C14B1.7 | WBGene00007578 |
| ZK418.8 | WBGene00022737 | C27H6.3 | WBGene00007785 |
| CSR-1 | WBGene00017641 | DCS-1 | WBGene00000940 |
| DRH-1 | WBGene00001090 | HRDE-1 | WBGene00007624 |
| EPI-1 | WBGene00001328 | MUT-14 | WBGene00003507 |
| ERI-1 | WBGene00001332 | NRDE-3 | WBGene00019862 |
| ERI-5 | WBGene00021419 | PRDE-1 | WBGene00008995 |
| LAM-1 | WBGene00002247 | RDE-12 | WBGene00010280 |
| RSD-2 | WBGene00004681 | WAGO-10 | WBGene00020707 |
| F20A1.9 | WBGene00017620 | AIN-1 | WBGene00015547 |
| MUT-15 | WBGene00011323 | ALG-1 | WBGene00000105 |
| RDE-1 | WBGene00004323 | ALG-2 | WBGene00000106 |
| SID-1 | WBGene00004795 |  |  |

**Table S3:**
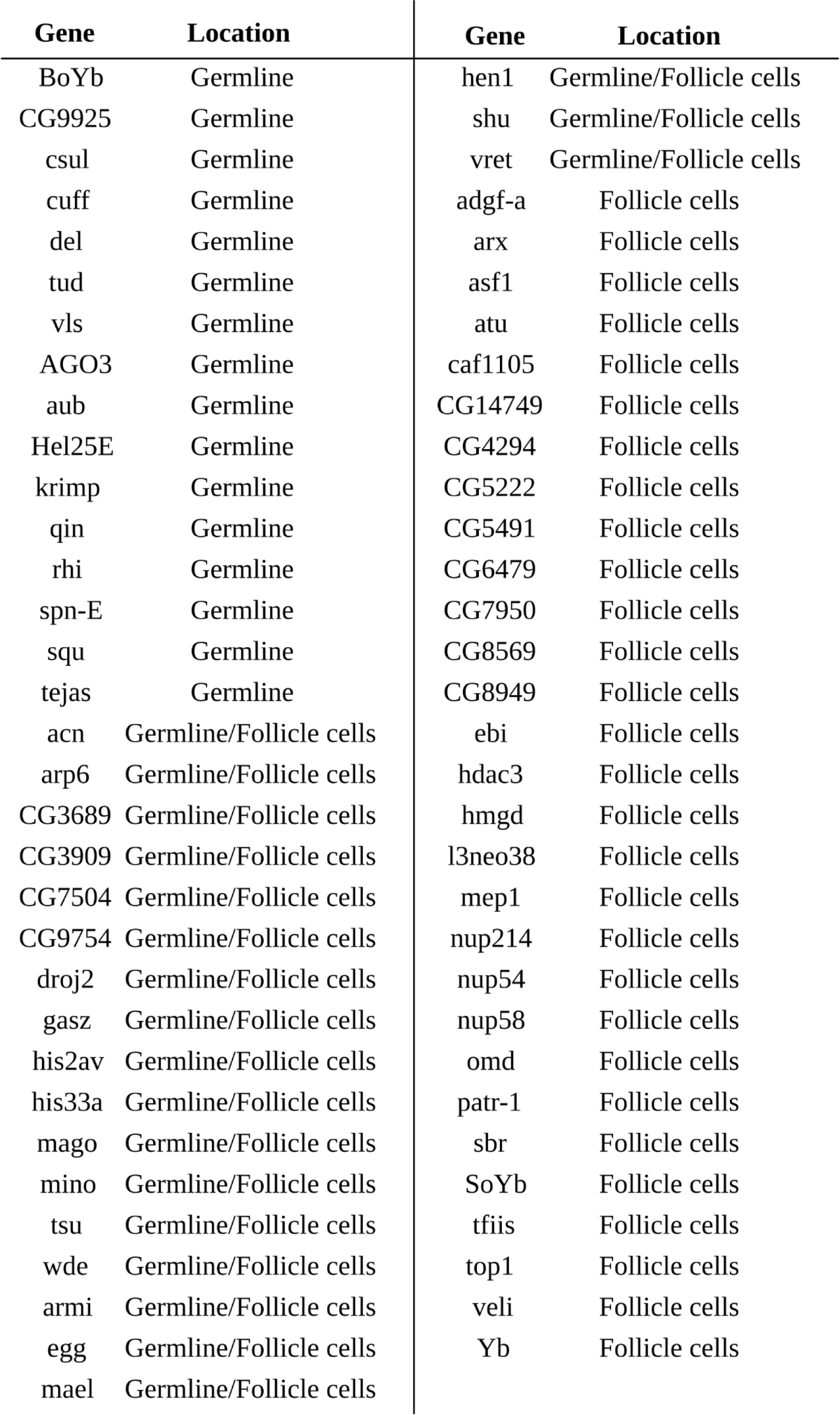
Larger set of piRNA-implicated genes used to calculate rates of adaptive evolution in the germline and soma of *D. melanogaster*

| Gene | Location | Gene | Location |
| --- | --- | --- | --- |
| BoYb | Germline | hen1 | Germline/Follicle cells |
| CG9925 | Germline | shu | Germline/Follicle cells |
| csul | Germline | vret | Germline/Follicle cells |
| cuff | Germline | adgf-a | Follicle cells |
| del | Germline | arx | Follicle cells |
| tud | Germline | asf1 | Follicle cells |
| vls | Germline | atu | Follicle cells |
| AGO3 | Germline | caf1105 | Follicle cells |
| aub | Germline | CG14749 | Follicle cells |
| Hel25E | Germline | CG4294 | Follicle cells |
| krimp | Germline | CG5222 | Follicle cells |
| qin | Germline | CG5491 | Follicle cells |
| rhi | Germline | CG6479 | Follicle cells |
| spn-E | Germline | CG7950 | Follicle cells |
| squ | Germline | CG8569 | Follicle cells |
| tejas | Germline | CG8949 | Follicle cells |
| acn | Germline/Follicle cells | ebi | Follicle cells |
| arp6 | Germline/Follicle cells | hdac3 | Follicle cells |
| CG3689 | Germline/Follicle cells | hmgd | Follicle cells |
| CG3909 | Germline/Follicle cells | l3neo38 | Follicle cells |
| CG7504 | Germline/Follicle cells | mep1 | Follicle cells |
| CG9754 | Germline/Follicle cells | nup214 | Follicle cells |
| droj2 | Germline/Follicle cells | nup54 | Follicle cells |
| gasz | Germline/Follicle cells | nup58 | Follicle cells |
| his2av | Germline/Follicle cells | omd | Follicle cells |
| his33a | Germline/Follicle cells | patr-1 | Follicle cells |
| mago | Germline/Follicle cells | sbr | Follicle cells |
| mino | Germline/Follicle cells | SoYb | Follicle cells |
| tsu | Germline/Follicle cells | tfiis | Follicle cells |
| wde | Germline/Follicle cells | top1 | Follicle cells |
| armi | Germline/Follicle cells | veli | Follicle cells |
| egg | Germline/Follicle cells | Yb | Follicle cells |
| mael | Germline/Follicle cells |  |  |

**Table S4.**
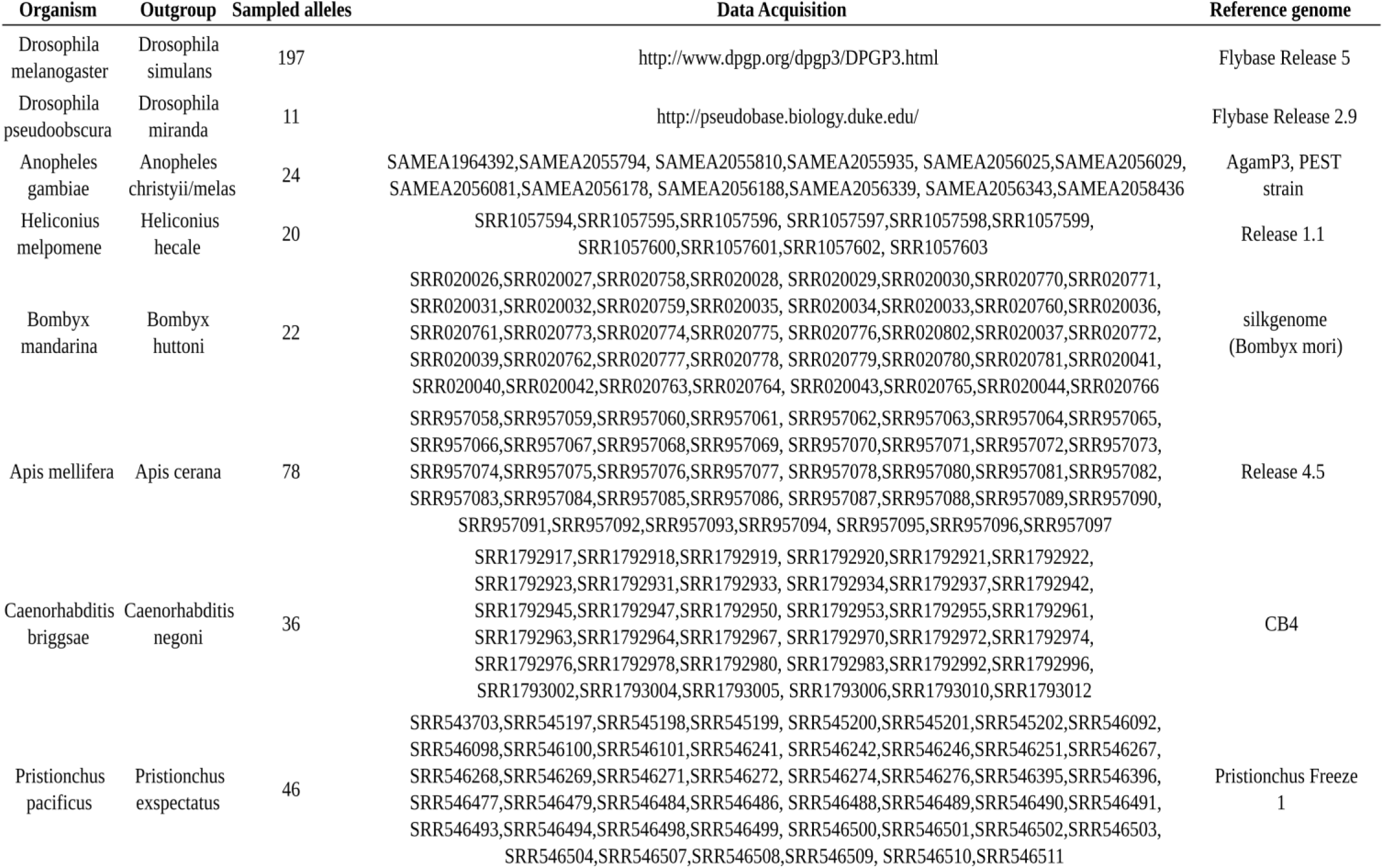
Accession numbers for public data and genome assembly used for each species

**Table S5.** Genes with significantly elevated selection effects

| Species | Gene | Duplicate | Selection Effect | 95% HPD interval | Species | Gene | Duplicate | Selection Effect | 95% HPD interval |
| --- | --- | --- | --- | --- | --- | --- | --- | --- | --- |
| *D. melanogaster* | arx | A | 1.16 | [0.157-2.09] | *D. pseudoobscura* | vas | A | 1.16 | [0.28-2.04] |
| *D. melanogaster* | piwi | A | 1.11 | [0.294-1.94] | *D. pseudoobscura* | vig | A | 1.49 | [0.337-2.65] |
| *D. melanogaster* | csul | A | 1.53 | [0.693-2.33] | *D. pseudoobscura* | vret | B | 1.48 | [0.703-2.3] |
| *D. melanogaster* | Dcr-1 | A | 1.21 | [0.441-2] | *D. pseudoobscura* | wde | A | 0.876 | [0.251-1.5] |
| *D. melanogaster* | Dcr-2 | A | 1.03 | [0.151-1.89] | *D. pseudoobscura* | Yb | A | 1.75 | [0.871-2.74] |
| *D. melanogaster* | krimp | A | 1.12 | [0.174-1.94] | *D. pseudoobscura* | Yb | B | 0.737 | [0.144-1.31] |
| *D. melanogaster* | qin | A | 1.04 | [0.421-1.61] | *D. pseudoobscura* | zuc | A | 1.43 | [0.649-2.15] |
| *D. melanogaster* | r2d2 | A | 1.59 | [0.625-2.66] | *Heliconius* | shu | A | 0.756 | [0.0191-1.51] |
| *D. melanogaster* | tapas | A | 1.35 | [0.639-2.03] | *Heliconius* | Yb | A | 1.03 | [0.483-1.57] |
| *D. melanogaster* | tej | A | 0.826 | [0.21-1.4] | *Apis* | AGO2 | A | 1.55 | [0.76-2.37] |
| *D. melanogaster* | tud | A | 0.816 | [0.0691-1.51] | *Apis* | piwi | A | 1.04 | [0.306-1.79] |
| *D. melanogaster* | vas | A | 1.63 | [0.975-2.33] | *Apis* | Dcr-2 | A | 0.965 | [0.195-1.72] |
| *D. melanogaster* | vret | A | 0.995 | [0.211-1.72] | *Apis* | rm62 | C | 1.12 | [0.02-2.13] |
| *D. melanogaster* | Yb | C | 1.2 | [0.472-1.93] | *Apis* | tud | A | 0.941 | [0.253-1.61] |
| *D. melanogaster* | SoYb | B | 0.78 | [0.262-1.3] | *Apis* | vas | A | 1.46 | [0.467-2.48] |
| *D. pseudoobscura* | AGO2 | A | 1.07 | [0.307-1.86] | *Anopheles* | AGO3 | A | 0.934 | [0.103-1.85] |
| *D. pseudoobscura* | armi | B | 2.66 | [2.09-3.25] | *Anopheles* | armi | A | 0.82 | [0.0614-1.57] |
| *D. pseudoobscura* | armi | A | 1.22 | [0.622-1.82] | *Anopheles* | arx | A | 1.39 | [0.376-2.33] |
| *D. pseudoobscura* | arx | B | 1.01 | [0.0294-2.11] | *Anopheles* | piwi | B | 1.26 | [0.569-1.92] |
| *D. pseudoobscura* | piwi | A | 0.952 | [0.043-1.81] | *Anopheles* | csul | A | 1.04 | [0.141-1.91] |
| *D. pseudoobscura* | csul | A | 0.921 | [0.00777-1.83] | *Anopheles* | Dcr-1 | A | 0.772 | [0.101-1.43] |
| *D. pseudoobscura* | cuff | A | 1.26 | [0.41-2.12] | *Anopheles* | drosha | A | 1.54 | [0.721-2.37] |
| *D. pseudoobscura* | cuff | B | 1.33 | [0.458-2.16] | *Anopheles* | loqs | A | 1.68 | [0.794-2.54] |
| *D. pseudoobscura* | Dcr-2 | A | 1.48 | [0.762-2.27] | *Anopheles* | mei-p26 | A | 1.08 | [0.234-1.89] |
| *D. pseudoobscura* | del | A | 1.33 | [0.413-2.17] | *Anopheles* | papi | A | 0.98 | [0.203-1.79] |
| *D. pseudoobscura* | drosha | A | 1.62 | [0.788-2.47] | *Anopheles* | tej | A | 0.868 | [0.169-1.53] |
| *D. pseudoobscura* | egg | A | 0.949 | [0.249-1.63] | *Anopheles* | trax | A | 1.48 | [0.433-2.48] |
| *D. pseudoobscura* | loqs | A | 1.33 | [0.301-2.42] | *Anopheles* | tud | A | 1.37 | [0.716-2.03] |
| *D. pseudoobscura* | mael | A | 1.22 | [0.0958-2.36] | *Anopheles* | vig | A | 1.29 | [0.196-2.4] |
| *D. pseudoobscura* | mael | B | 1.22 | [0.0897-2.36] | *Anopheles* | wde | A | 0.957 | [0.338-1.58] |
| *D. pseudoobscura* | mei-p26 | A | 1.66 | [0.852-2.38] | *Anopheles* | zuc | A | 1.42 | [0.535-2.31] |
| *D. pseudoobscura* | papi | A | 0.79 | [0.0114-1.56] | *Bombyx* | AGO2 | A | 1.41 | [0.633-2.18] |
| *D. pseudoobscura* | pasha | A | 1.2 | [0.247-2.14] | *Bombyx* | Dcr-1 | A | 2.95 | [2.14-3.75] |
| *D. pseudoobscura* | qin | A | 1.56 | [0.975-2.16] | *Bombyx* | egg | A | 2.06 | [1.26-2.92] |
| *D. pseudoobscura* | r2d2 | A | 1.29 | [0.22-2.37] | *Bombyx* | Hel25E | A | 1.18 | [0.0422-2.35] |
| *D. pseudoobscura* | rm62 | B | 1.33 | [0.247-2.29] | *Bombyx* | mael | A | 1.04 | [0.324-1.75] |
| *D. pseudoobscura* | rm62 | A | 1.09 | [0.0839-2.01] | *Bombyx* | papi | A | 1.15 | [0.11-2.18] |
| *D. pseudoobscura* | shu | A | 1.57 | [0.667-2.48] | *Bombyx* | shu | A | 1.31 | [0.64-1.97] |
| *D. pseudoobscura* | spn-E | A | 1.97 | [1.31-2.59] | *Bombyx* | tapas | A | 0.989 | [0.233-1.74] |
| *D. pseudoobscura* | squ | A | 1.19 | [0.26-2.09] | *Bombyx* | trsn | A | 1.73 | [0.595-2.94] |
| *D. pseudoobscura* | tej | A | 1.13 | [0.171-2] | *Bombyx* | vig | A | 1.72 | [0.432-2.92] |
| *D. pseudoobscura* | trsn | A | 1.54 | [0.622-2.56] | *Bombyx* | vret | A | 0.714 | [0.0928-1.33] |
| *D. pseudoobscura* | tud | A | 0.912 | [0.343-1.53] | *Bombyx* | Yb | A | 1.34 | [0.778-1.87] |

**Figure S1.**
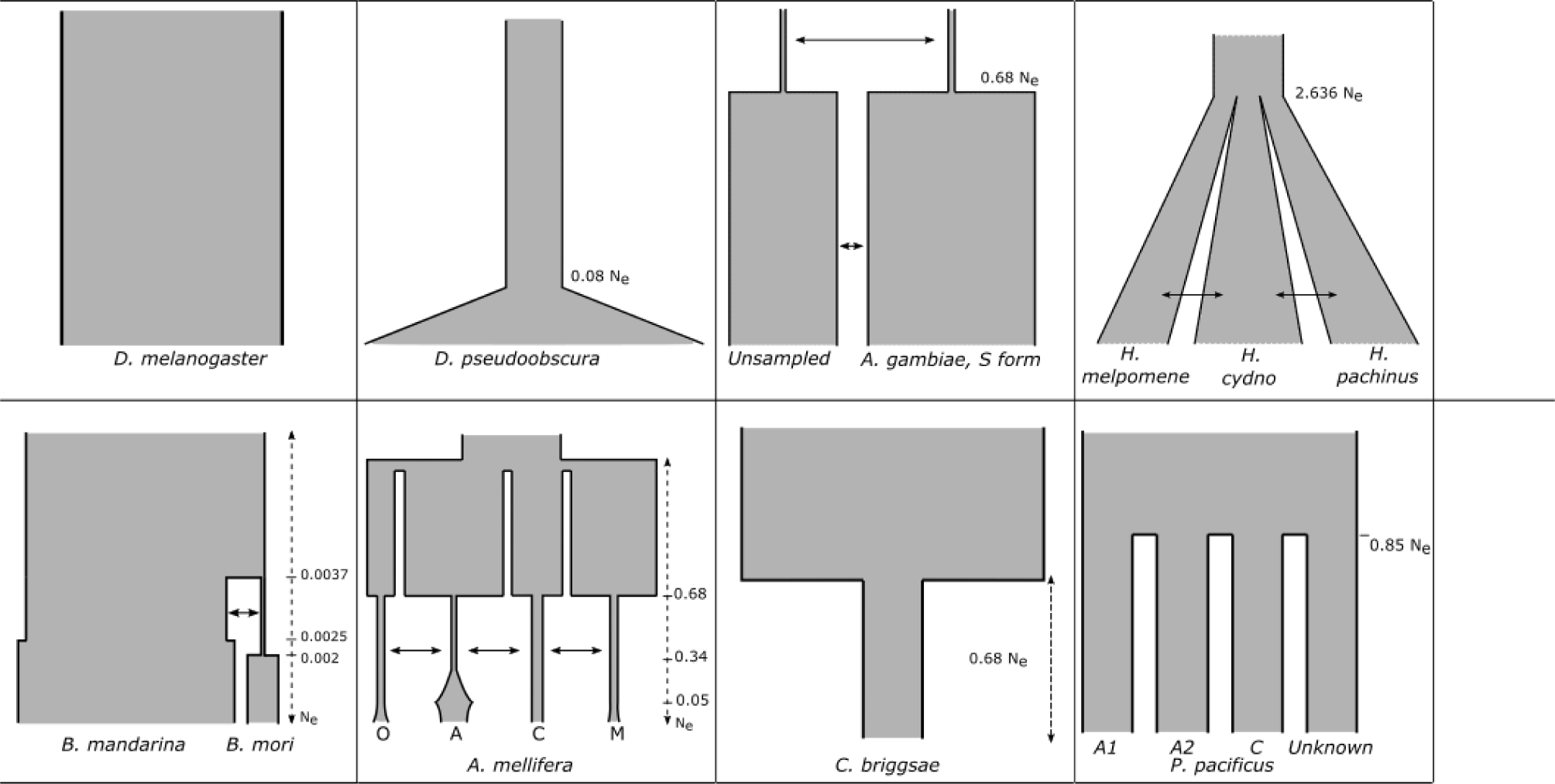
Demographic scenarios simulated for SweeD analysis. Coalescent simulations were performed using ms for demographic scenarios for each species which are supported by other studies. The African (Zambia) *D. melanogaster* were assumed to have a constant population size. *D. pseudoobscura* has recently undergone a population expansion 0.08 Ne generations ago. *A. gambiae* shares migrants with some other unknown, unsampled subpopulation which split 0.68 Ne generations ago. *Heliconius* species in Costa Rica split 2.636 Ne generations ago and have shared migrants since. *Bombyx mandarina* went through a small bottleneck when *B. mori* split, and shared migrants during that bottleneck (but not after). *Apis mellifera* have four subpopulations which have gone through multiple population expansions and bottlenecks, with all subpopulations sharing migrants until they join 0.68 Ne generations ago. *Caenorhabditis briggsae “*tropical samples” have undergone a population bottleneck 0.68 Ne generations ago. Finally, *Pristionchus pacificus* were sampled from four subpopulations, which split 0.85 Ne generations ago.

**Figure S2:**
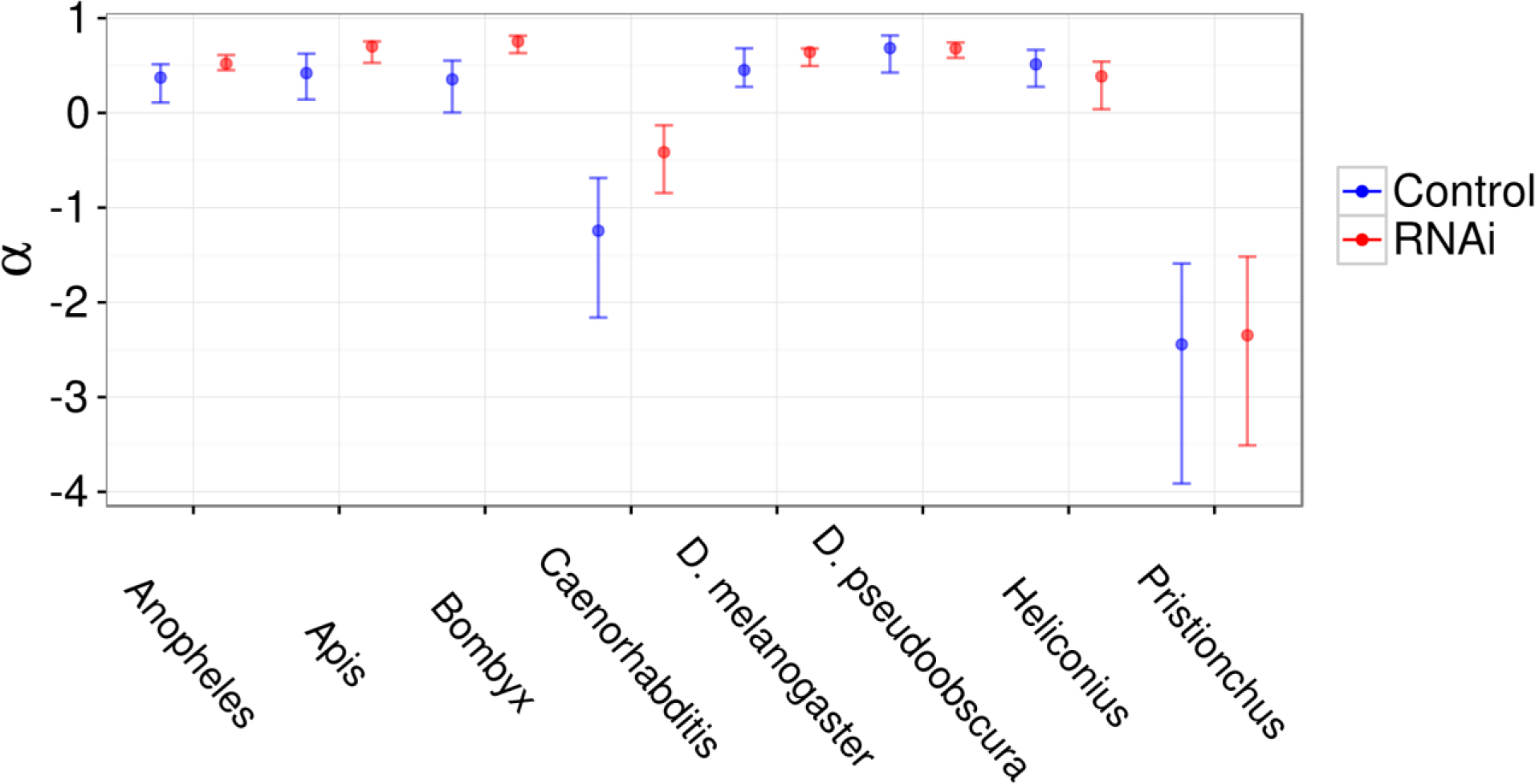
Alpha values for RNAi genes. For each species, α, or the proportion of adaptive substitutions was estimated from pooled polymorphism and divergence data using DFE-alpha for RNAi genes and position-matched control genes. α estimates for control genes are fairly constant across insect species, but are negative in the two nematode species. In all species except *H. melpomene*, the RNAi gene estimates are greater in RNAi genes than control genes.

**Figure S3.**
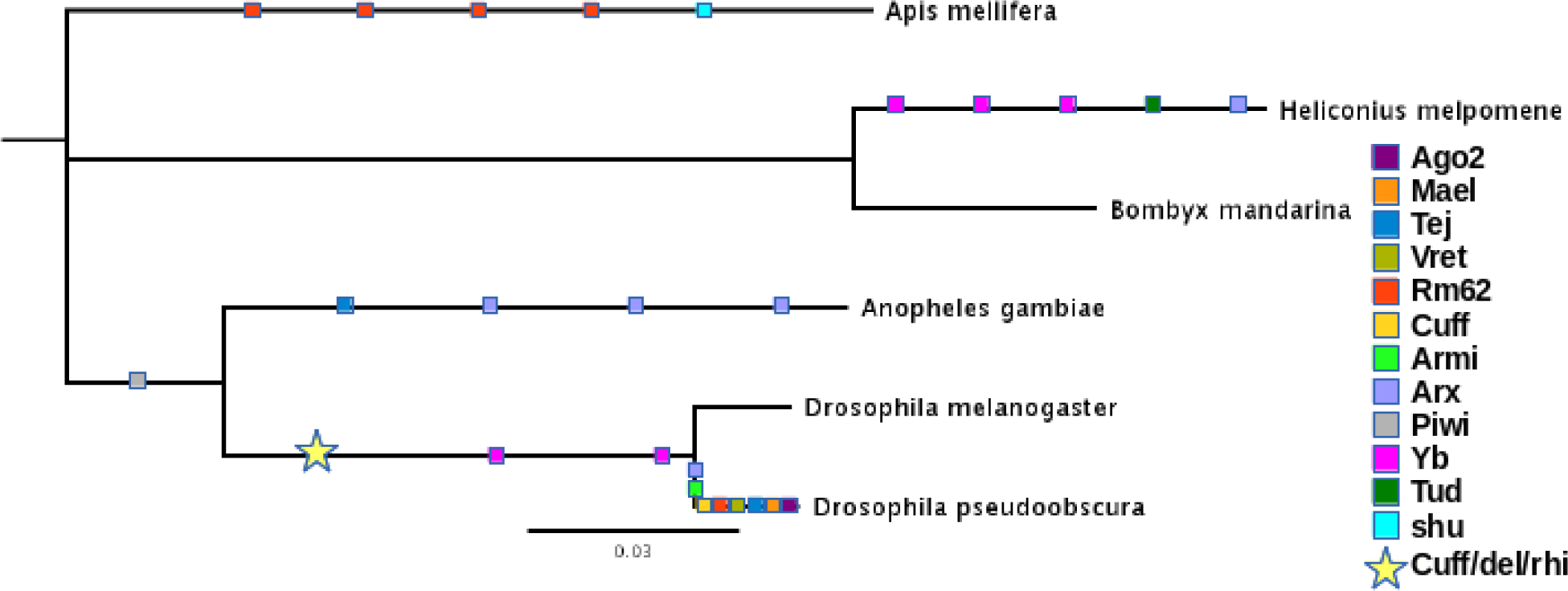
Possible duplications in RNAi pathway. Relationships of the insect species sampled, including coloured squares where possible gene duplications have occurred. Our search for RNAi genes in insect species other than *D. melanogaster* identified numerous duplications, and also some genes which were specific to *Drosophila*. Of note, *D. pseudoobscura* harboured duplications in *asterix, armitage, cutoff, rm62, vretano, tejas, maelstrom,* in addition to the multiple *AGO2* duplications reported previously (Lewis et al, 2016; Lewis et al, 2016), perhaps indicating an extensive addition to RNAi related pathways. *Asterix* was further duplicated three times in *Anopheles* and once in *Heliconius*, and *A. mellifera* also has five distinct copies of *rm62*. Furthermore, *yb* duplications have occurred independently in the lineage leading to *H. melpomene* and the one leading to the *Drosophila* species. The piRNA cluster transcriptional complex composed of cutoff, deadlock, and rhino were only observed in the two *Drosophila* species (represented by a star), and thus have likely either been lost in the other species or have evolved in since the split between *Anopheles* and *Drosophila*.

**Figure S4.**
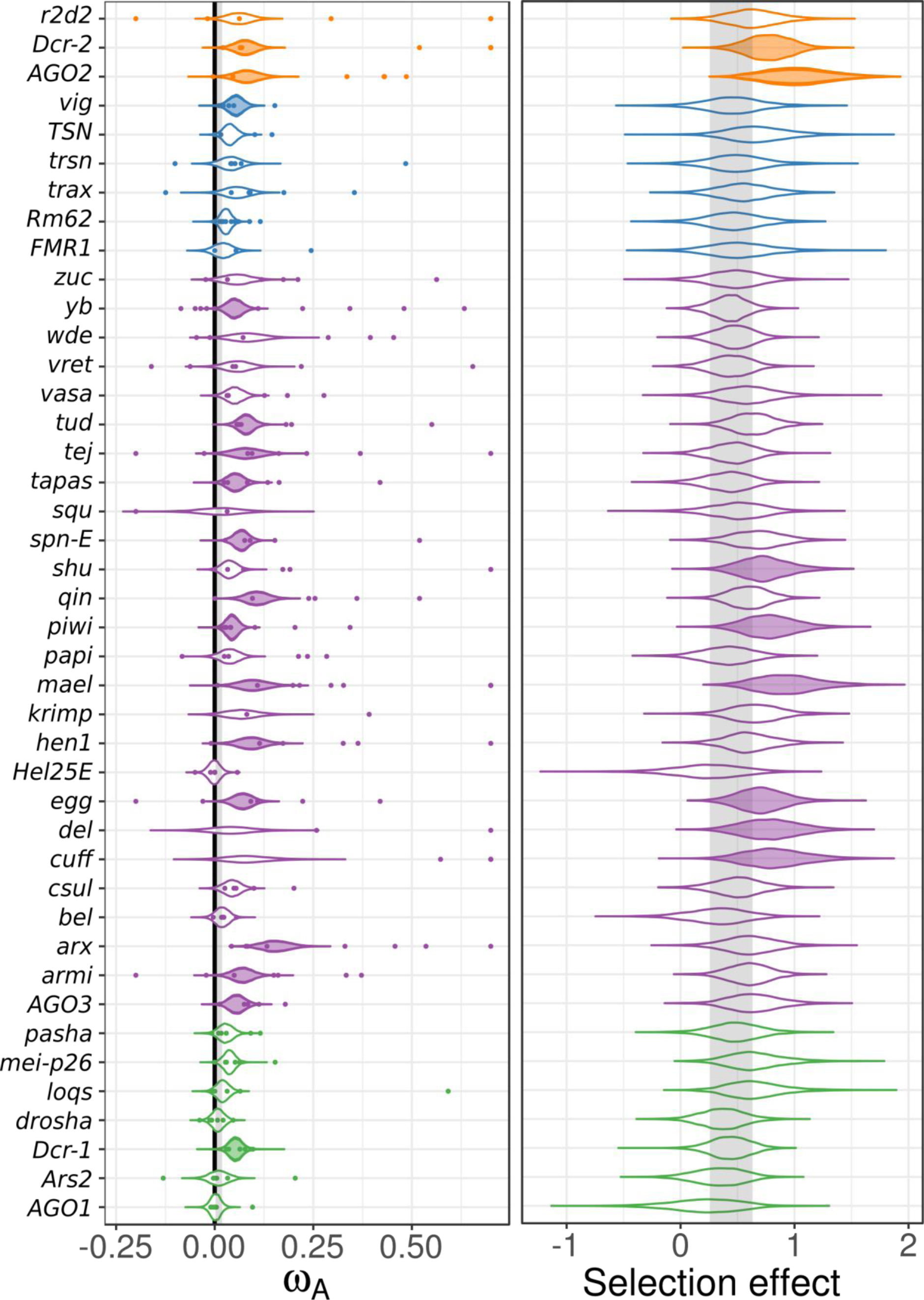
Cross-species homologue-level estimates of ω_A_ and selection effects without pathway assumptions. (Left) Individual gene ω_A_ estimates (coloured points) were calculated using DFE-alpha and analyses using a linear mixed model with species and gene as random effects (estimate uncertainty was included by incorporating bootstrap intervals as measurement error variance), but without subpathway as fixed effect (see Figure 4). The posterior distributions of the cross-species estimate for ω_A_ for each gene are plotted, and shaded if the MCMCp < 0.05 when tested against the control gene distribution (shaded grey region). Single-gene estimates of ω_A_ > 0.75 are plotted at 0.75 for clarity. (Right) The analogous analysis, except performed using SnIPRE, with the posterior distribution of homologue-level selection effects plotted. Both analyses find *AGO2, Dicer-2, piwi, maelstrom,* and *eggless* as having elevated protein substitution.

**Figure S5:**
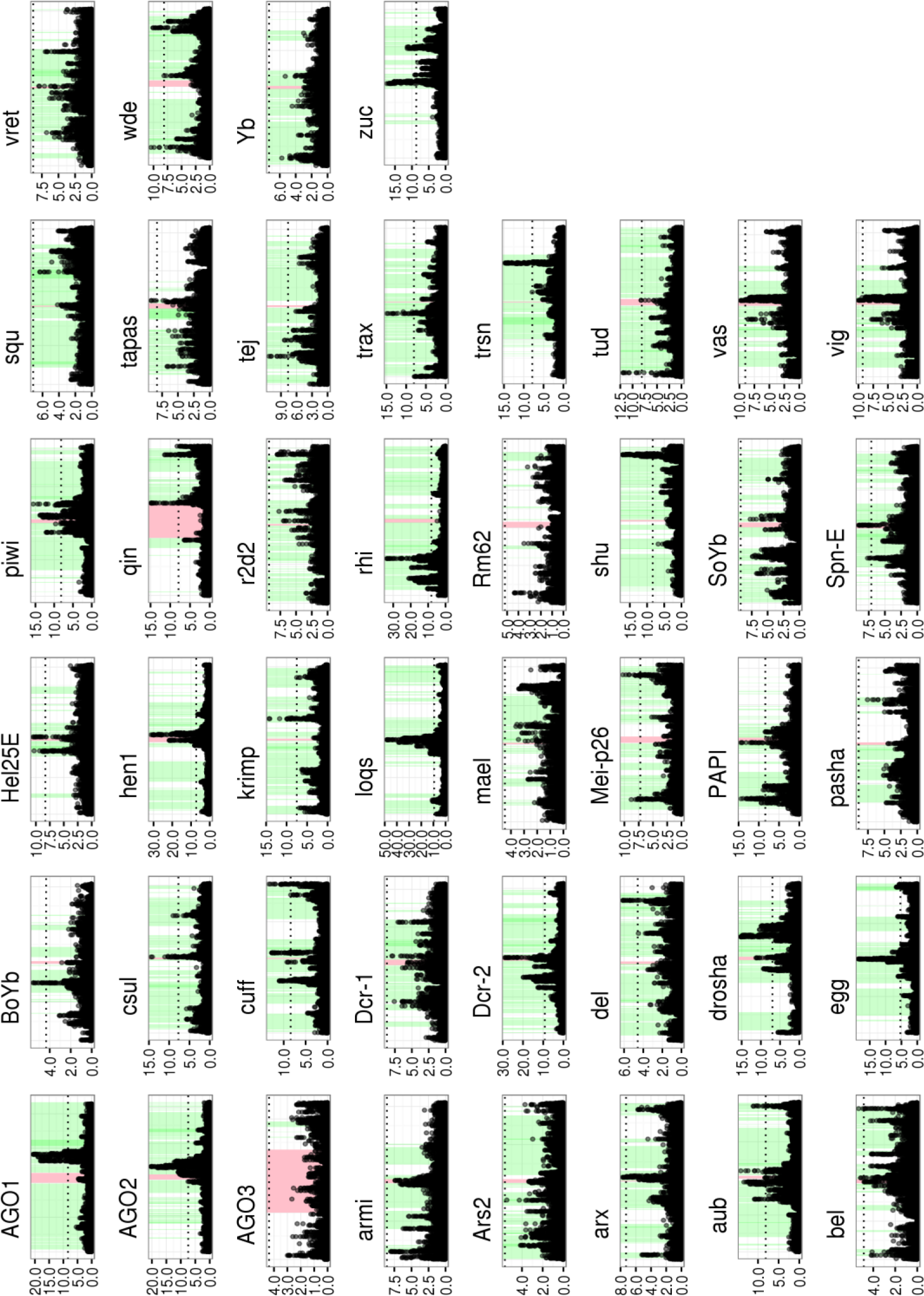
Drosophila melanogaster sweeps. For each *D. melanogaster* gene, the CLR statistic was plotted across a 200 kb region including the gene of interest. Each panel represents a region of the *D. melanogaster* genome, with red-shaded regions being the gene of interest and green-shaded regions being other genes along the chromosome. The horizontal dotted lines in each panel are significance thresholds calculated through neutral coalescent simulations.

**Figure S6:**
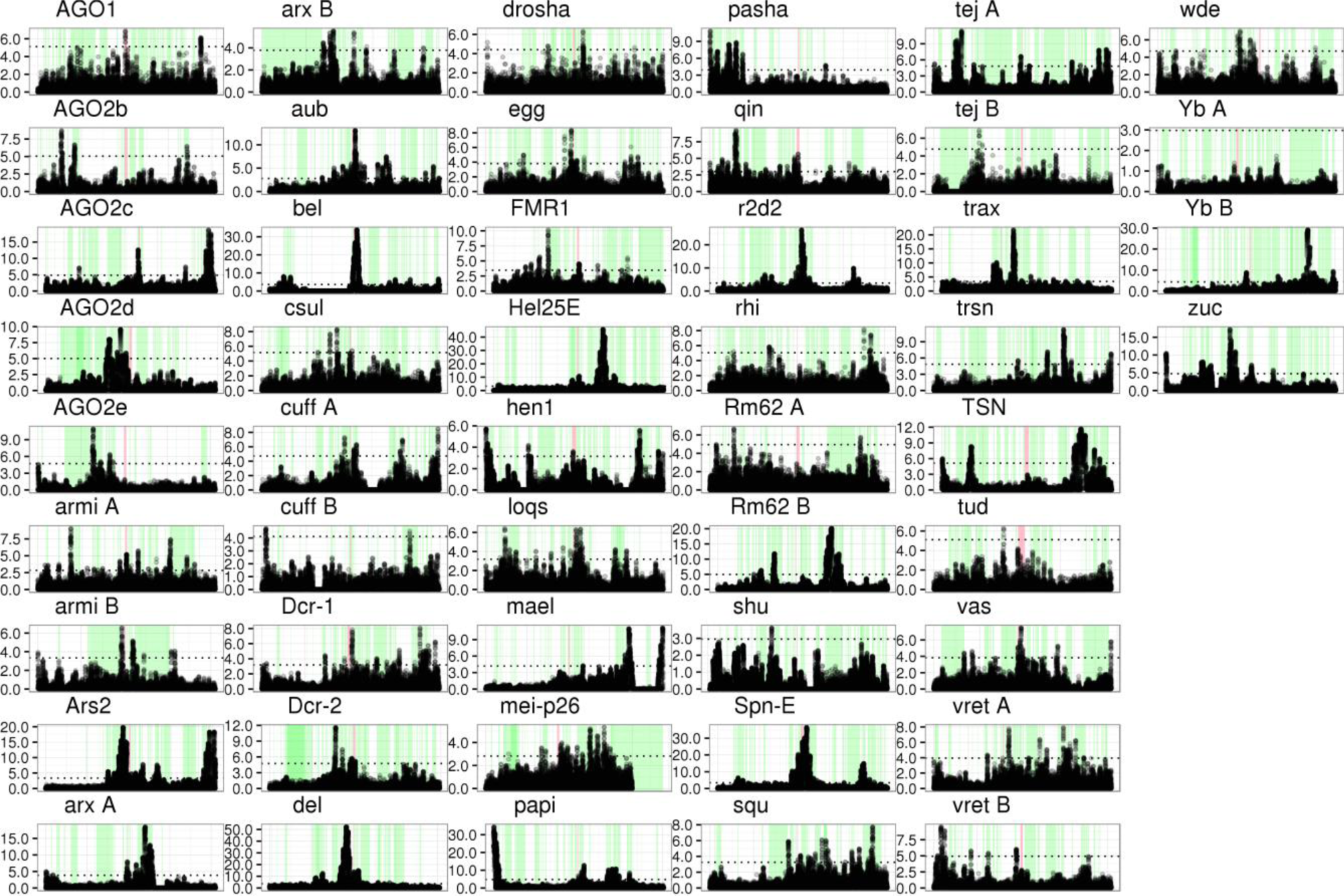
Drosophila pseudoobscura sweeps. For each *D. pseudoobscura* gene, the CLR statistic was plotted across a 200 kb region including the gene of interest. Each panel represents a region of the *D. pseudoobscura* genome, with red-shaded regions being the gene of interest and green-shaded regions being other genes along the chromosome. The horizontal dotted lines in each panel are significance thresholds calculated through neutral coalescent simulations.

**Figure S7:**
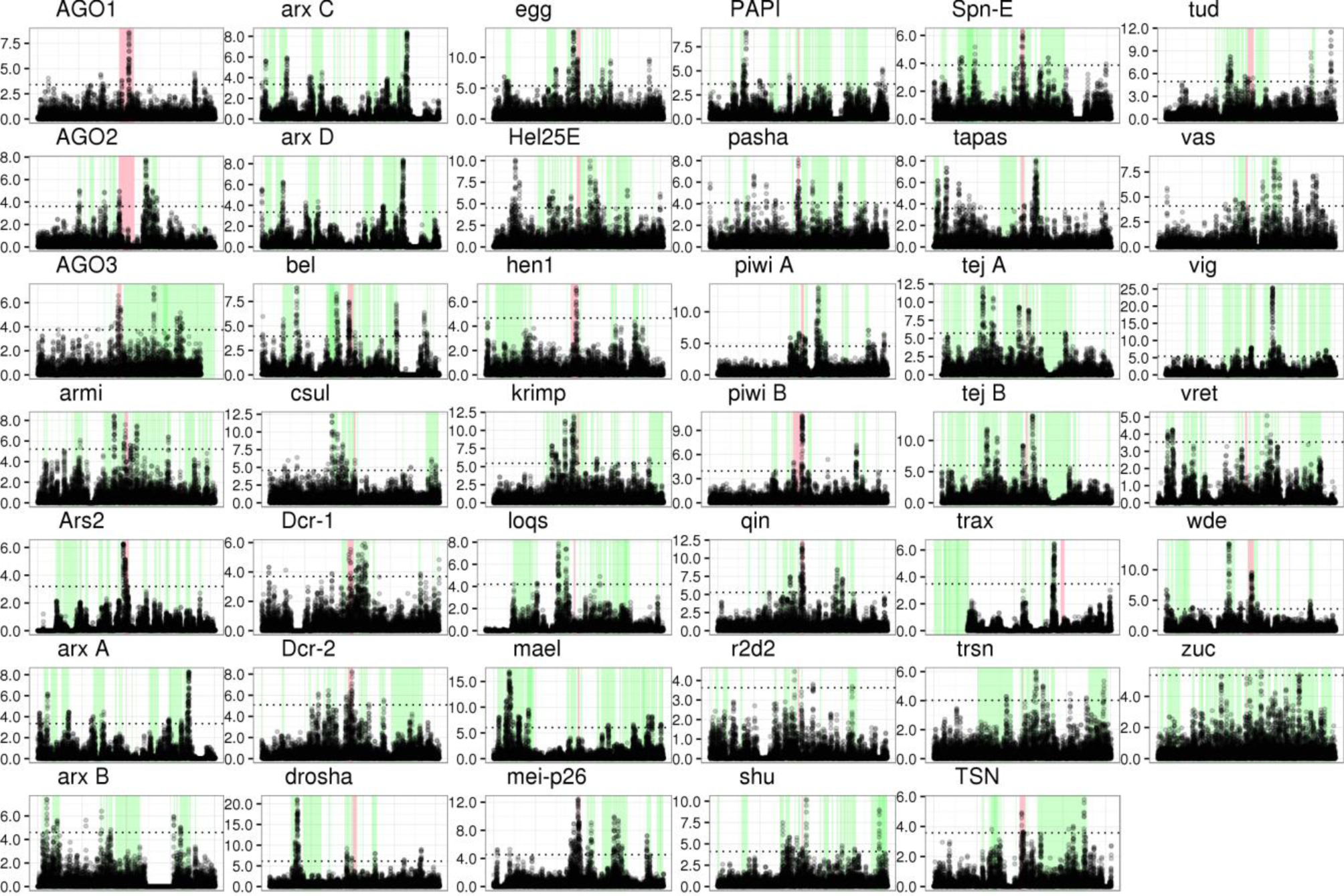
Anopheles gambiae sweeps. For each *A. gambiae* gene, the CLR statistic was plotted across a 200 kb region including the gene of interest. Each panel represents a region of the *A. gambiae* genome, with red-shaded regions being the gene of interest and green-shaded regions being other genes along the chromosome. The horizontal dotted lines in each panel are significance thresholds calculated through neutral coalescent simulations.

**Figure S8:**
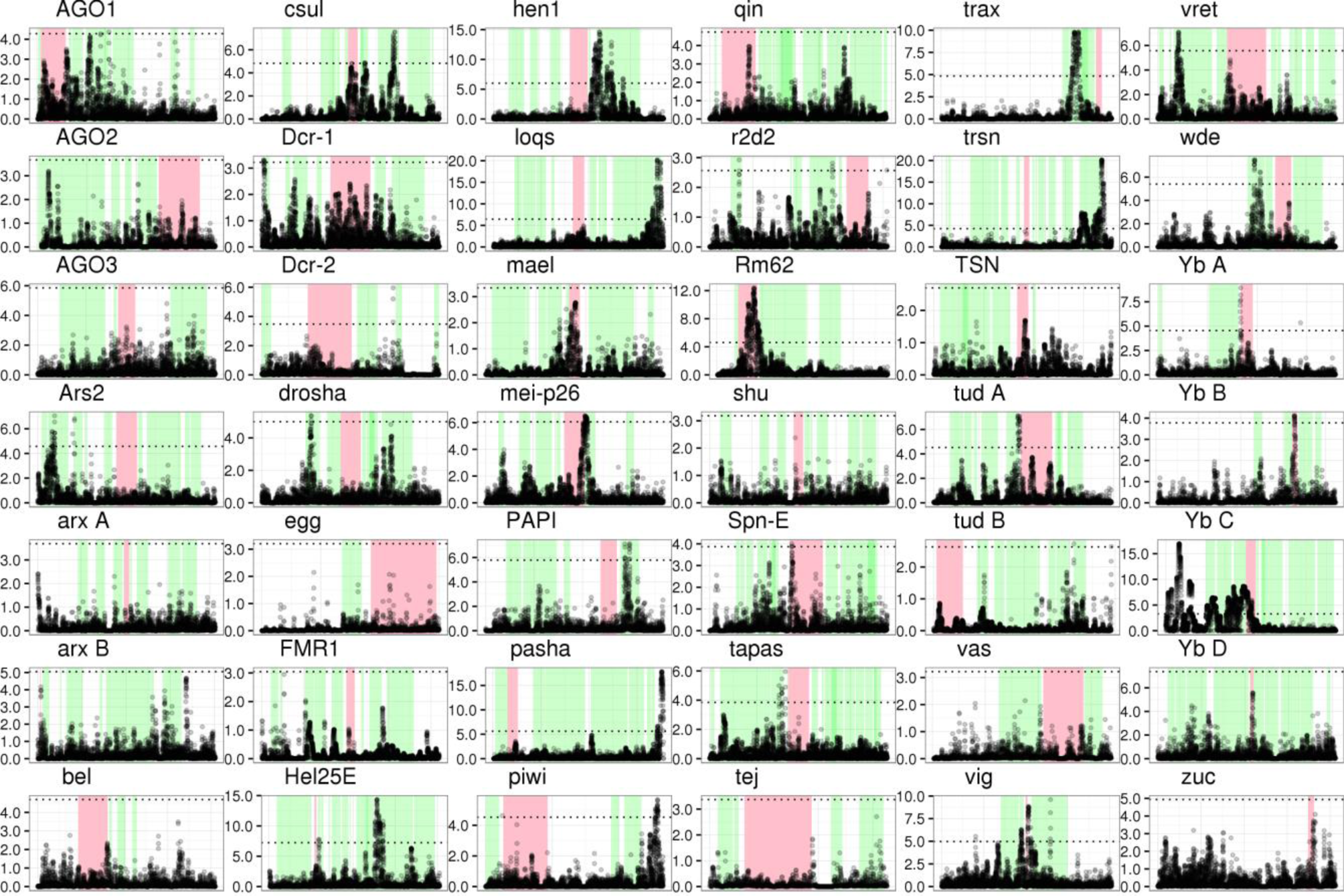
Heliconius melpomene sweeps. For each *H. melpomene* gene, the CLR statistic was plotted across a 200 kb region including the gene of interest. Each panel represents a region of the *H. melpomene* genome, with red-shaded regions being the gene of interest and green-shaded regions being other genes along the chromosome. The horizontal dotted lines in each panel are significance thresholds calculated through neutral coalescent simulations.

**Figure S9:**
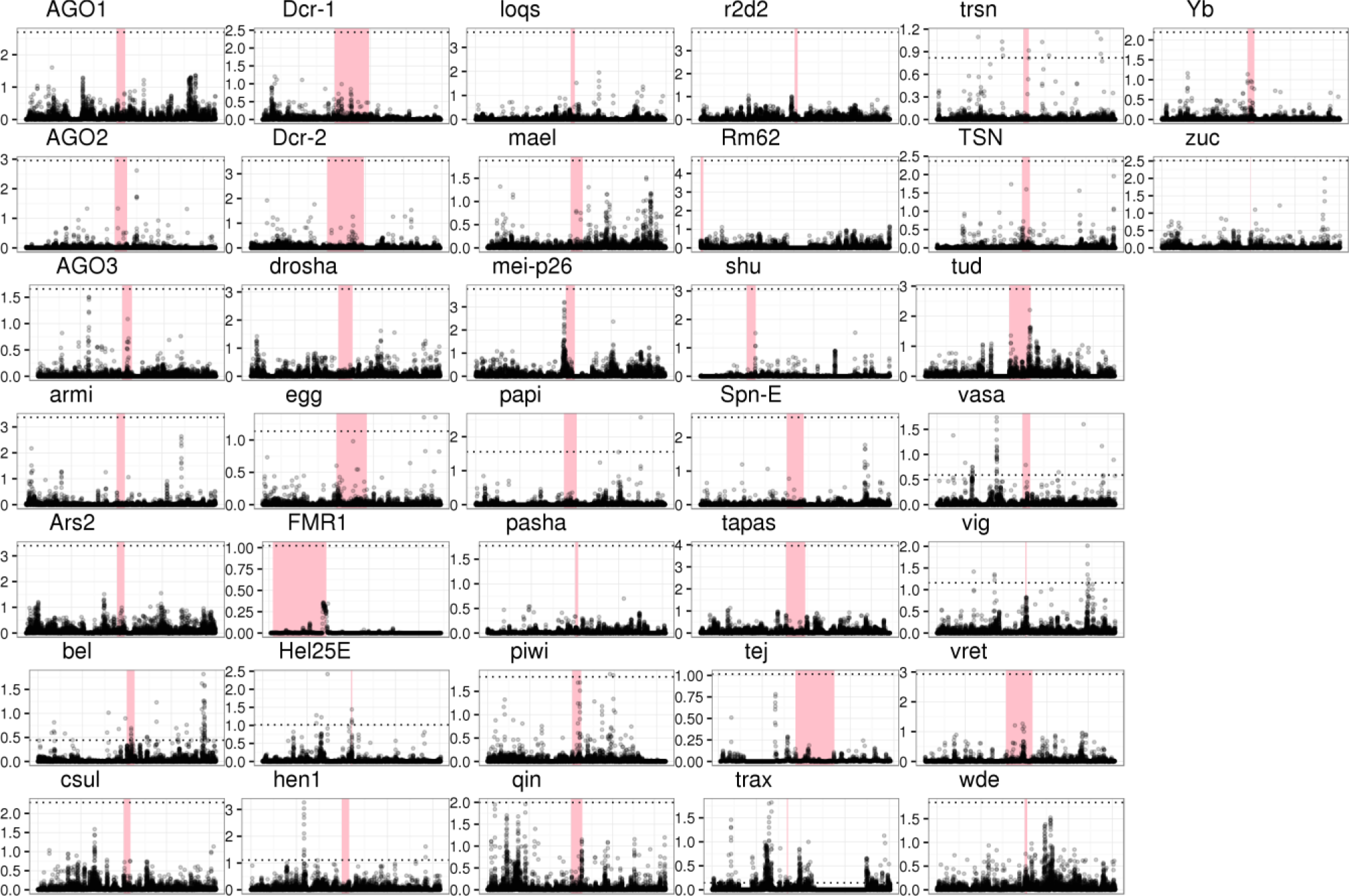
Bombyx mandarina sweeps. For each *B. mandarina* gene, the CLR statistic was plotted across a 200 kb region including the gene of interest. Each panel represents a region of the *B. mandarina* genome, with red-shaded regions being the gene of interest. The horizontal dotted lines in each panel are significance thresholds calculated through neutral coalescent simulations. The *Bombyx* genome used did not have an associated gff file, and so positions of nearby genes were not included.

**Figure S10:**
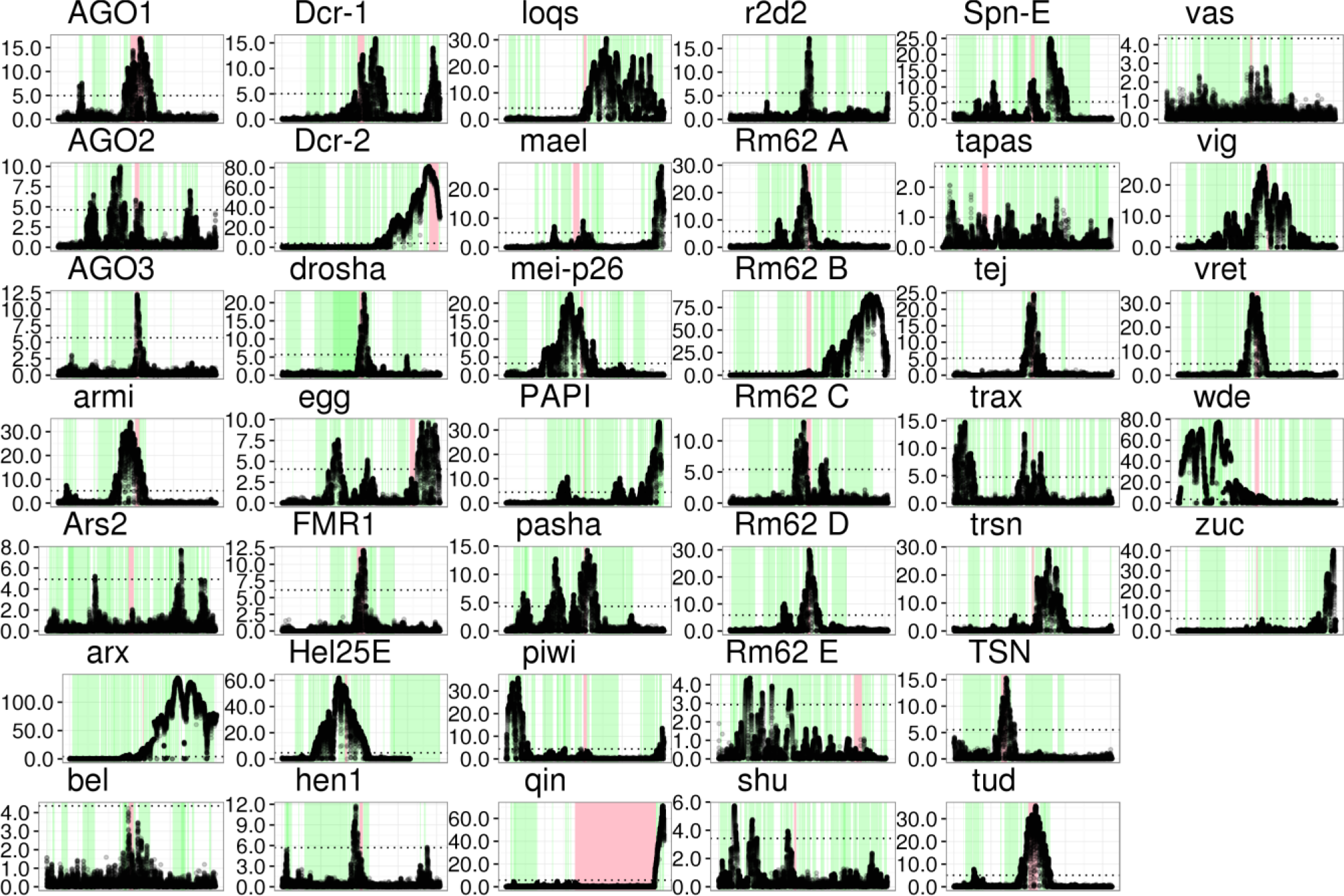
Apis mellifera sweeps. For each *A. mellifera* gene, the CLR statistic was plotted across a 200 kb region including the gene of interest. Each panel represents a region of the *A. mellifera* genome, with red-shaded regions being the gene of interest. The horizontal dotted lines in each panel are significance thresholds calculated through neutral coalescent simulations.

**Figure S11:**
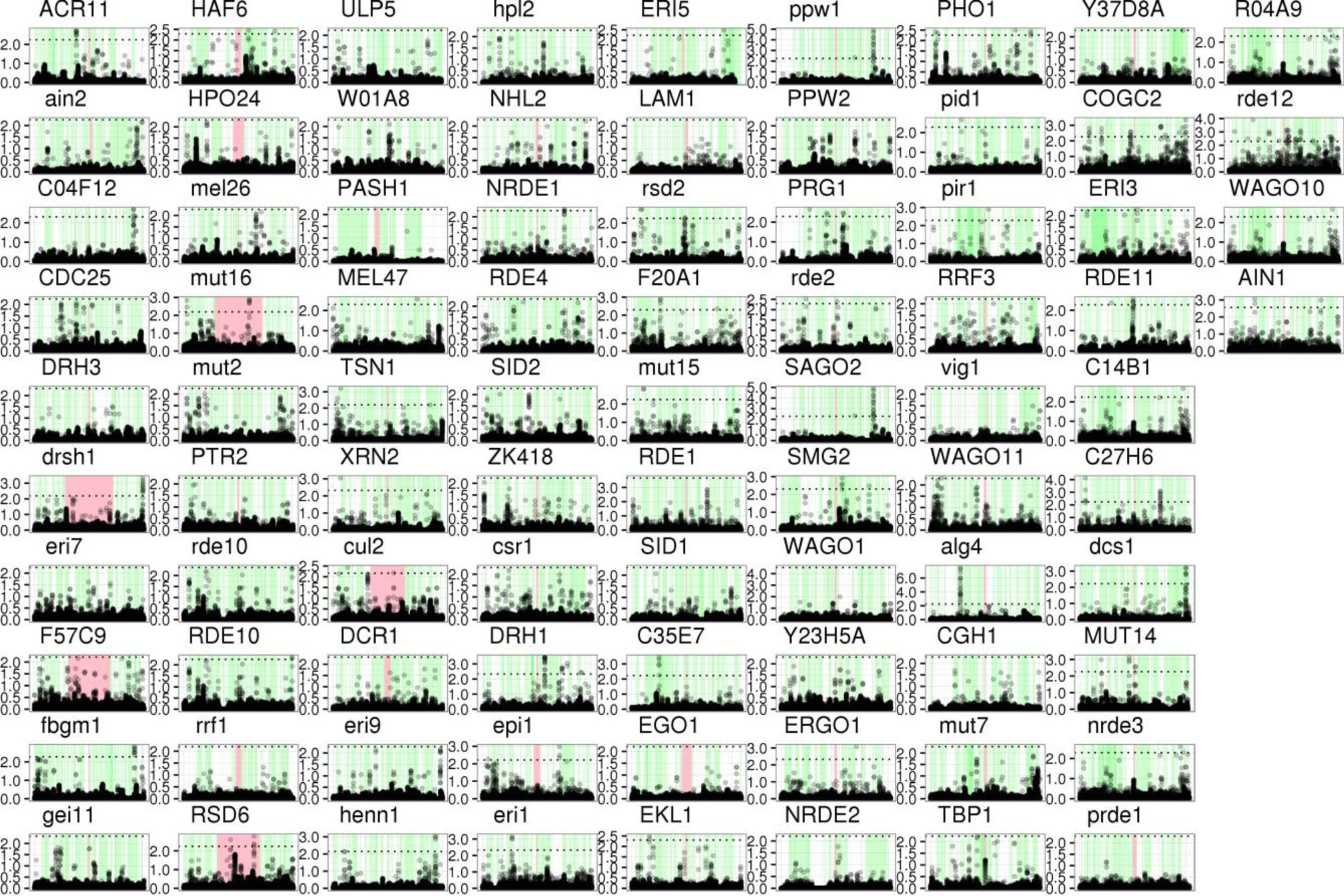
Caenorhabditis briggsae sweeps. For each *C. briggsae* gene, the CLR statistic was plotted across a 200 kb region including the gene of interest. Each panel represents a region of the *C. briggsae* genome, with red-shaded regions being the gene of interest. The horizontal dotted lines in each panel are significance thresholds calculated through neutral coalescent simulations.

**Figure S12:**
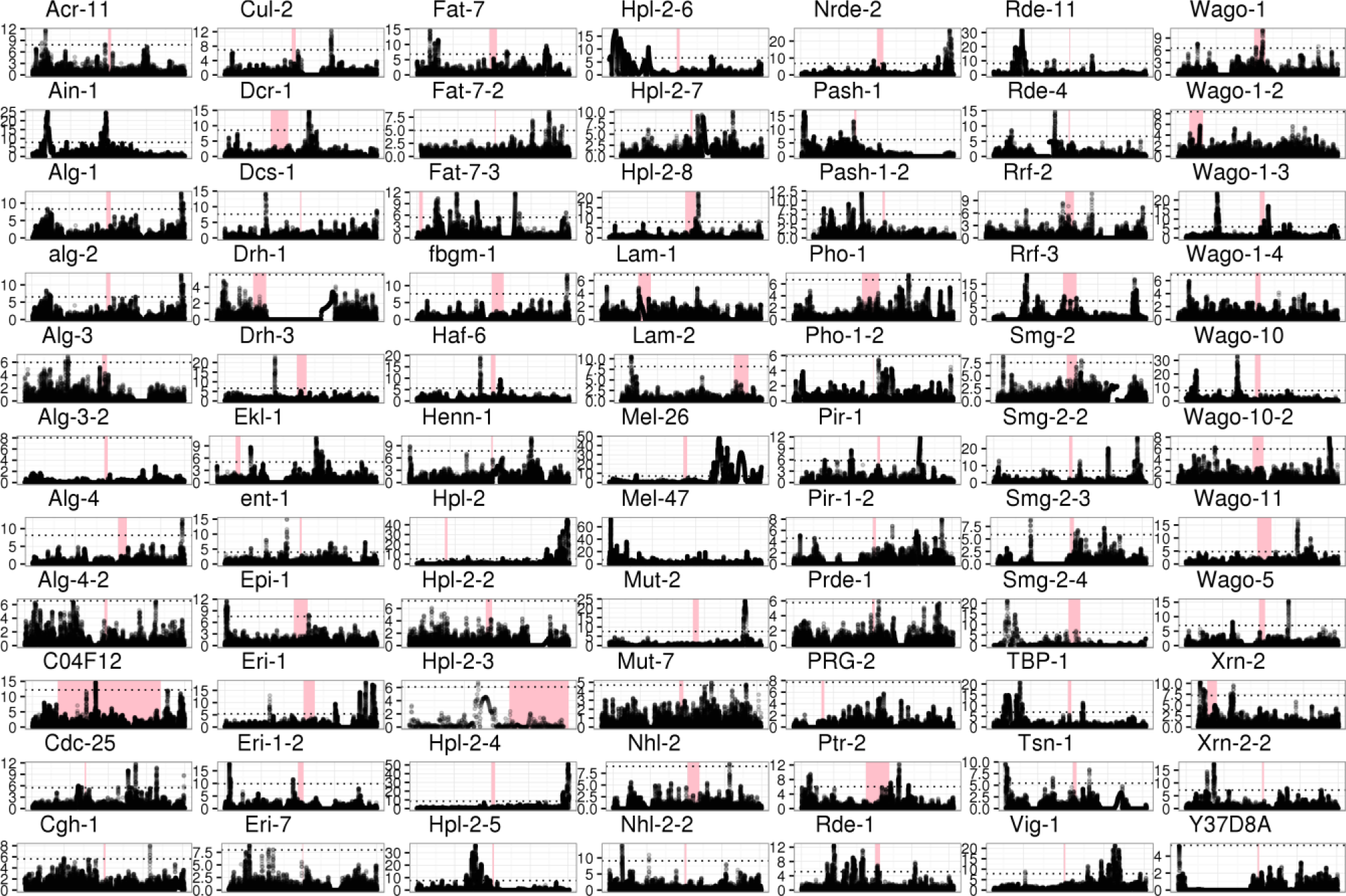
*Pristionchus pacificus* sweeps. For each *P. pacificus* gene, the CLR statistic was plotted across a 200 kb region including the gene of interest. Each panel represents a region of the *P. pacificus* genome, with red-shaded regions being the gene of interest. The horizontal dotted lines in each panel are significance thresholds calculated through neutral coalescent simulations. The *Pristionchus* genome used did not have an associated gff file, so positions of nearby genes were not included.

**Supp Figure 13:**
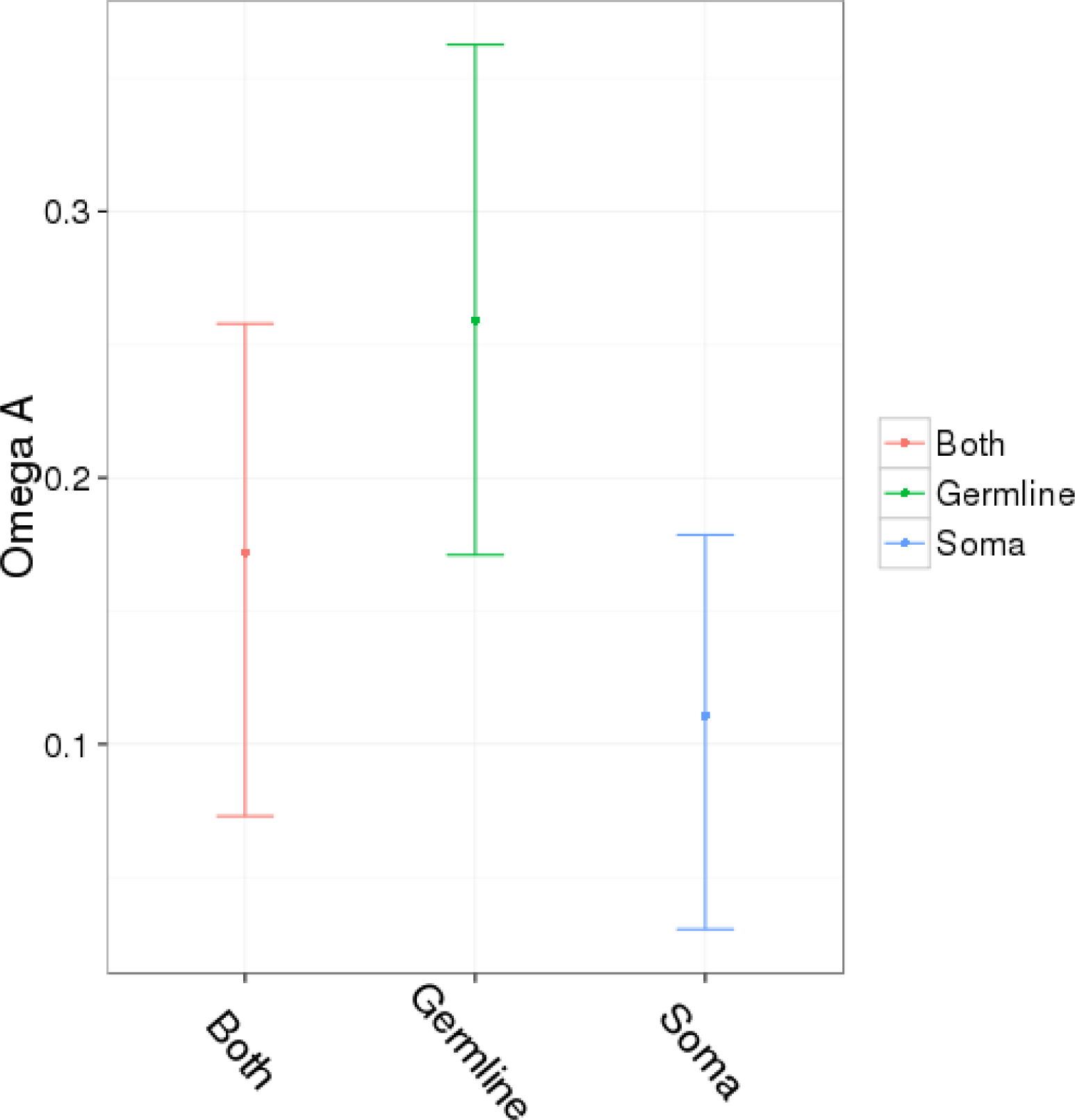
Germline and somatic piRNA pathway genes. Polymorphism and divergence from a larger set of piRNA pathway genes (those identified in two of three recent piRNA pathway screens, plus the core piRNA pathway) (Handler et al, 2013; Czech et al, 2013; Muerdter et al, 2013) in *D. melanogaster* were pooled based on whether they are active in the germline, soma, or both and used to calculate ω_A_. Confidence intervals were obtained by bootstrapping by gene 1000 times. Genes active in germline TE suppression show higher rates of adaptive protein evolution than those active in the somatic follicle cells, with genes active in both having an intermediate adaptive rate.

## S1 Text: Supplementary R code for models

DFE-alpha meta-analysis

Data set up:

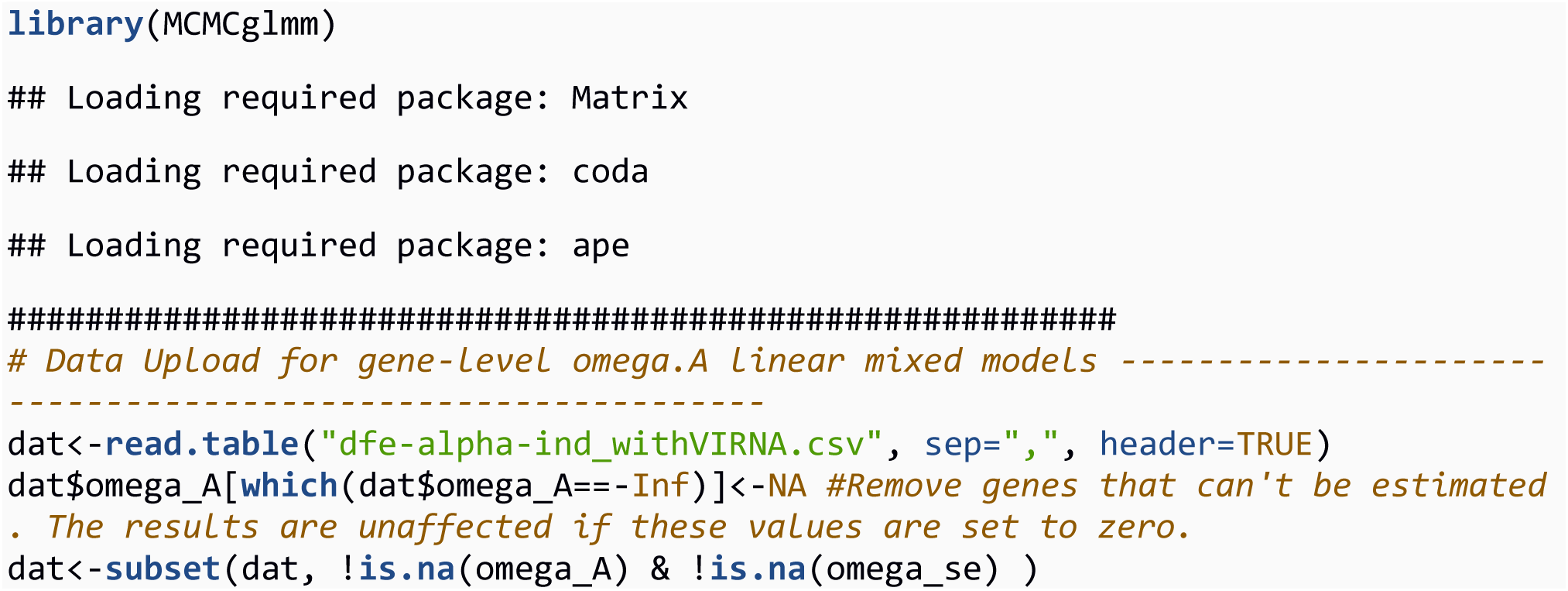

Model 1A: Comparison of RNAi and control genes

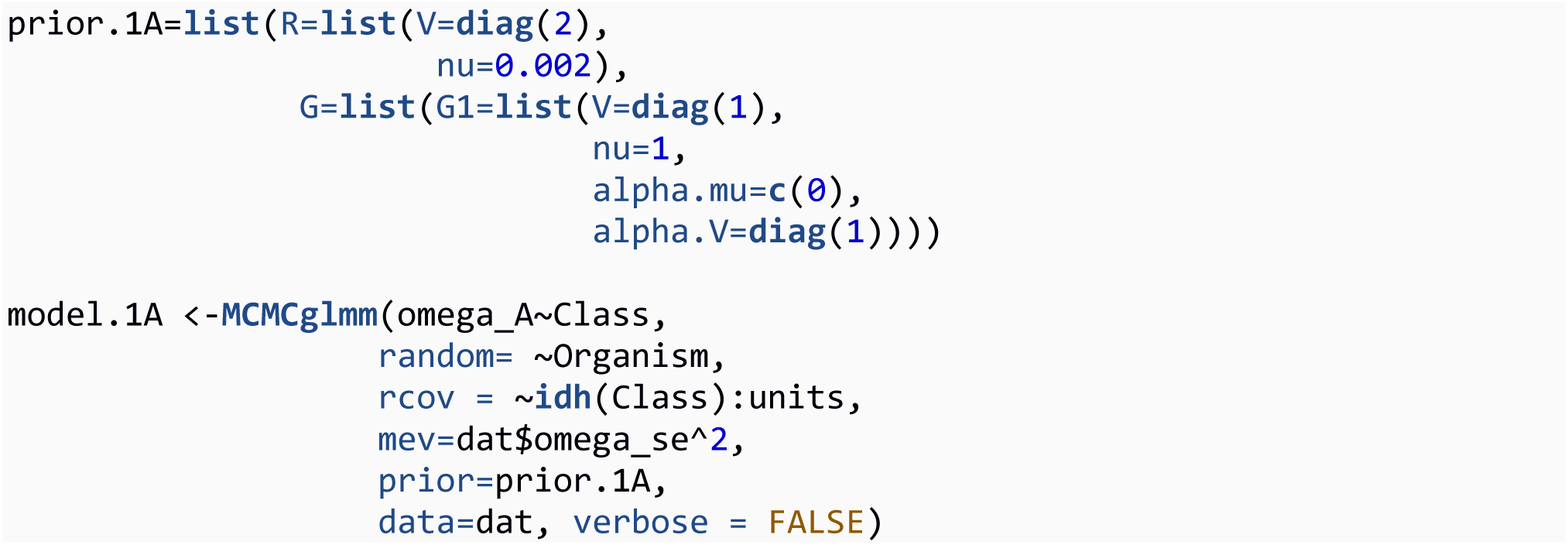

Model 1A models gene-level ωA estimates as a gaussian response with gene class as a fixed effect and species as a random effect. For the random effects (random= ~Organism), we assume all (co)variances among organisms are equal. We also estimate separate error variances for each gene class (rcov = ~idh(Class):units), allowing us to test whether the variance of the adaptive rate of RNAi and control genes differ. The idh() function specifies that the residual variance associated with each class of genes is independent, and sets the off-diagonals of the covariance matrix to zero. We specify the sampling error associated with each estimate of ωA (mev=dat$omega_se^2) obtained by bootstrapping by codon and rerunning DFE-alpha on the new codon set.

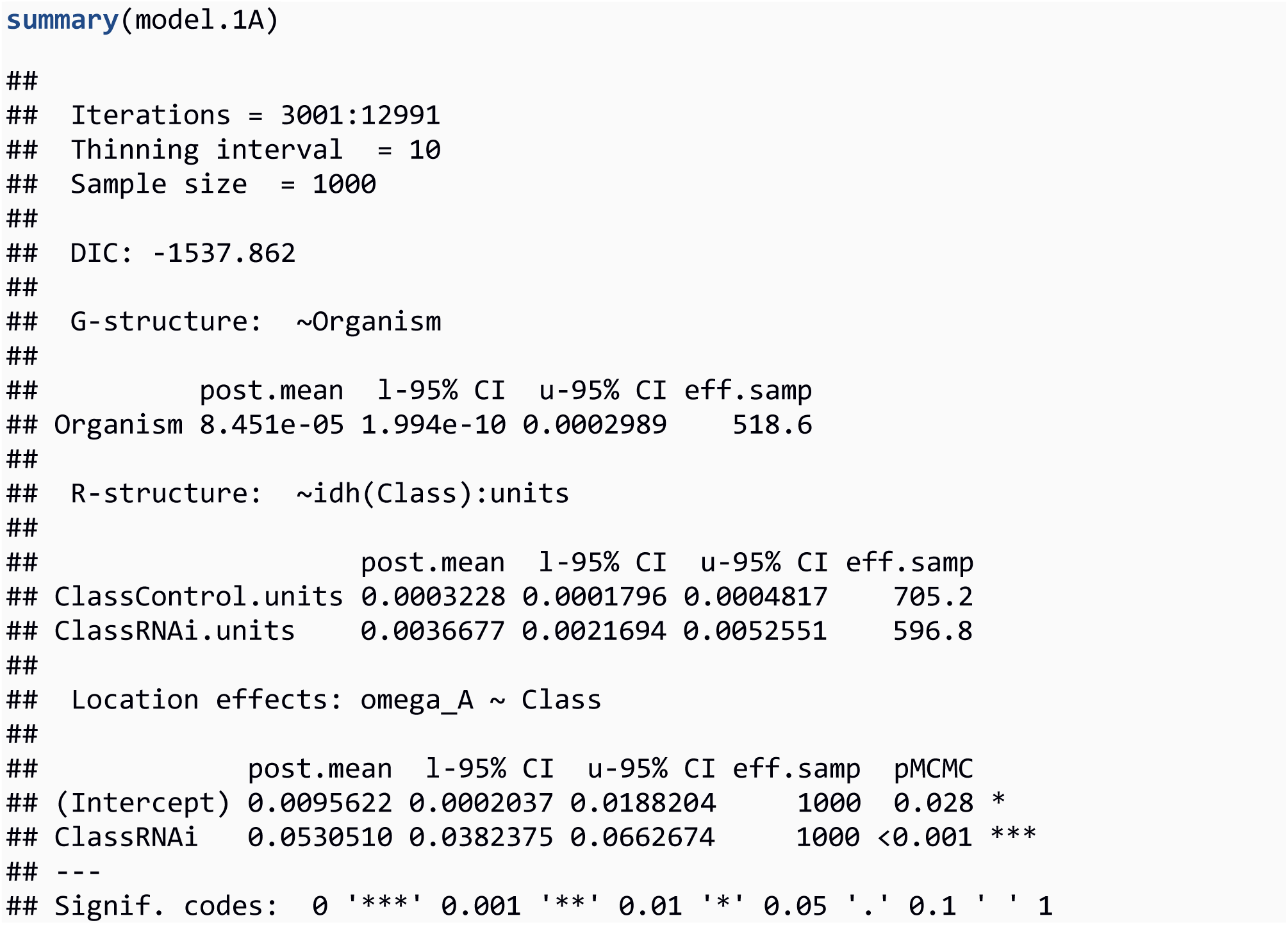

The command summary(model.1A) prints some aspects of the MCMC chain, the DIC score, the variance components for the G-structure (random effects), the error variance estimates, and the estimates for the fixed effects. The RNAi class of genes is estimated to be 0.05 greater than control genes, and this is signficant (pMCMC < 0.001) We also test whether the variance is significantly greater for RNAi genes using the posterior distributions for the error variances saved in the VCV object.

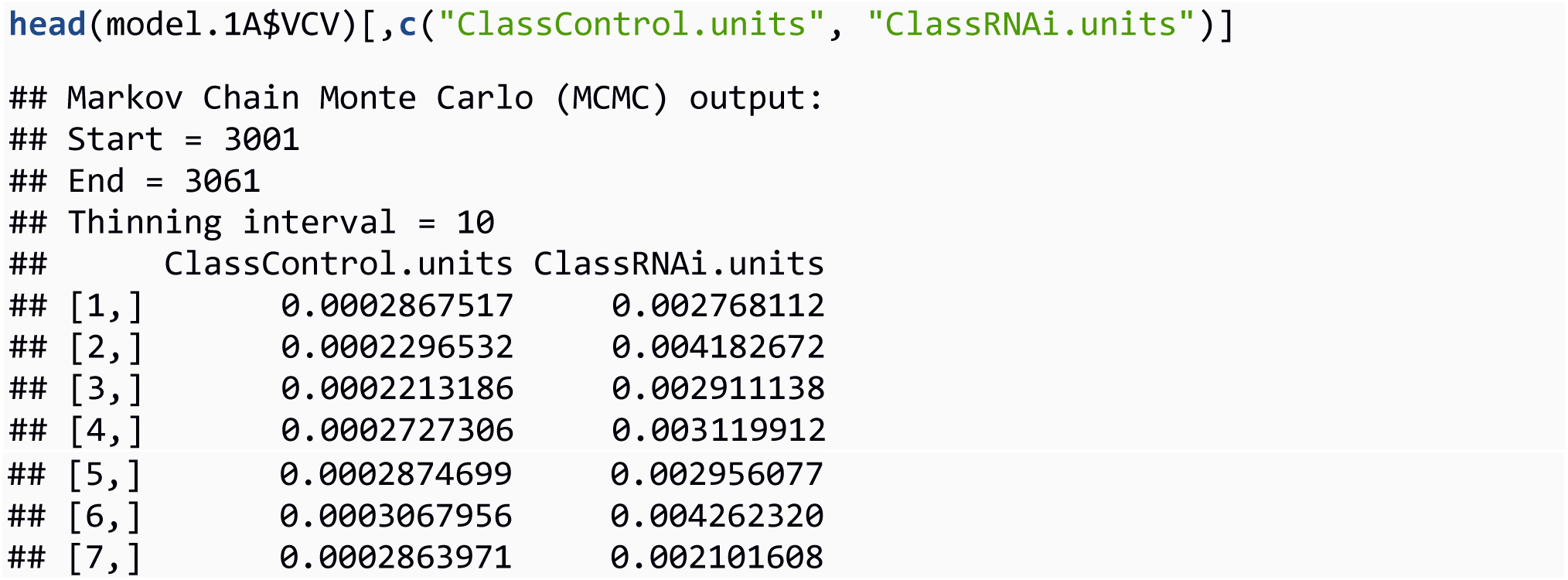

To test for significantly different variances, we subtract one posterior distribution from the other and ask what proportion overlaps zero.

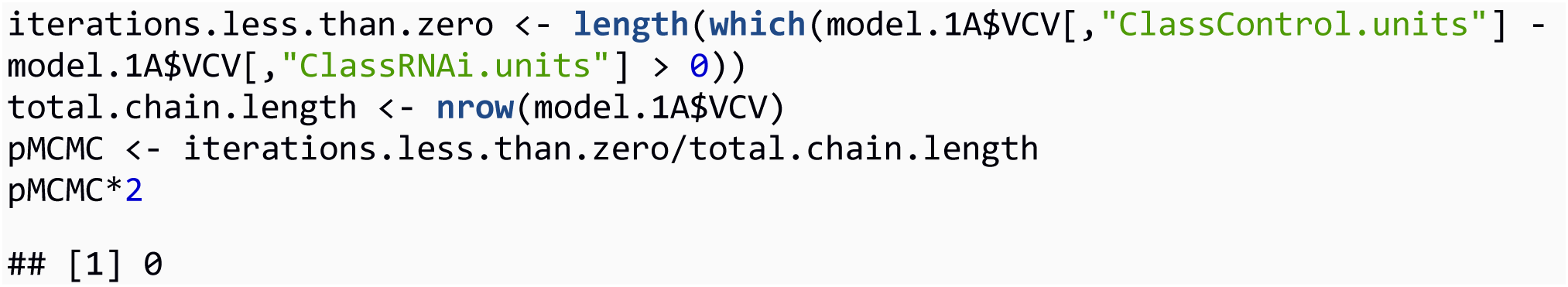

Therefore, in every iteration of the chain (1000 sampled iterations, thinning interval of 10), the error variance associated with RNAi genes was greater than control genes (MCMCp < 0.001).

Model 1B: Comparison of RNAi subpathways (piRNA, siRNA, viRNA, miRNA)

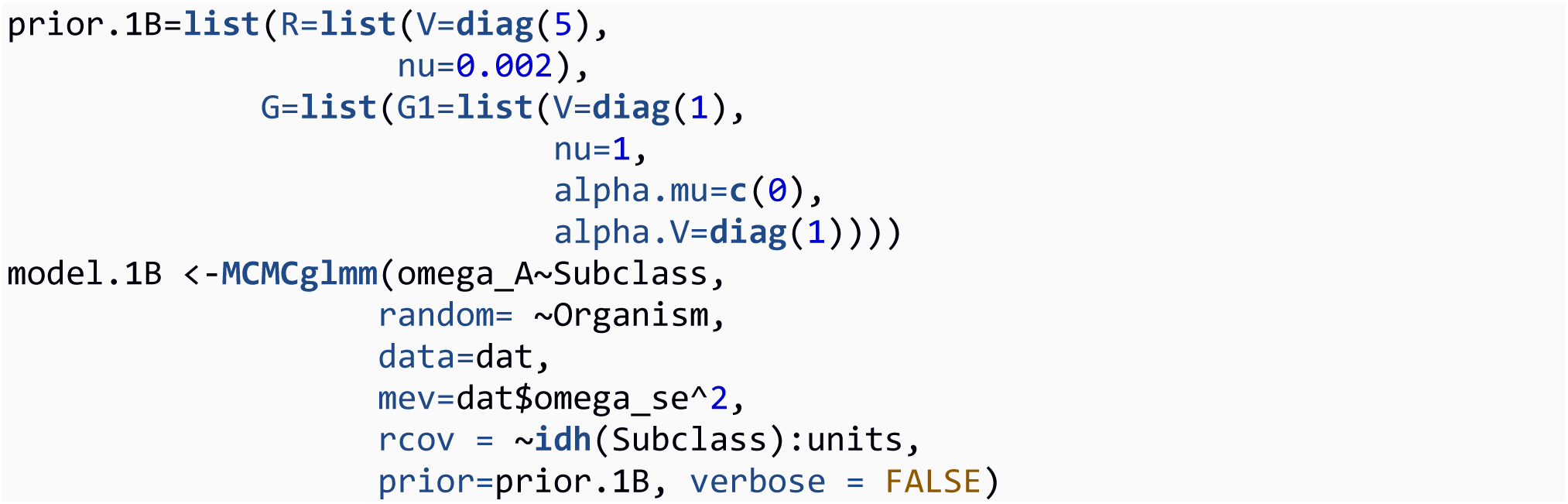

Model 1B is similar to model 1A, except the RNAi class has now been divided into four subpathways (miRNA, siRNA, piRNA, viRNA). The summary of the model output is the following:

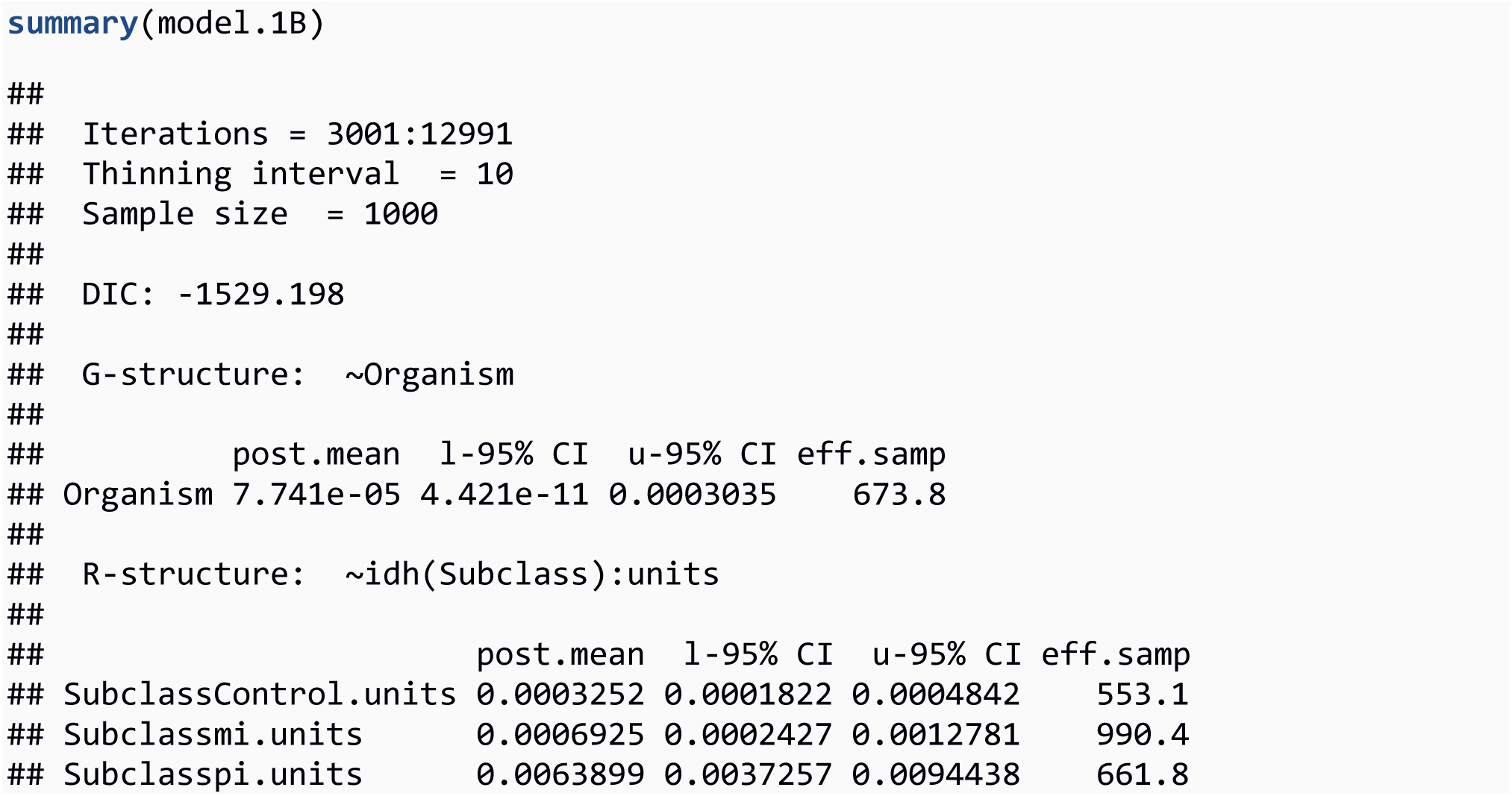
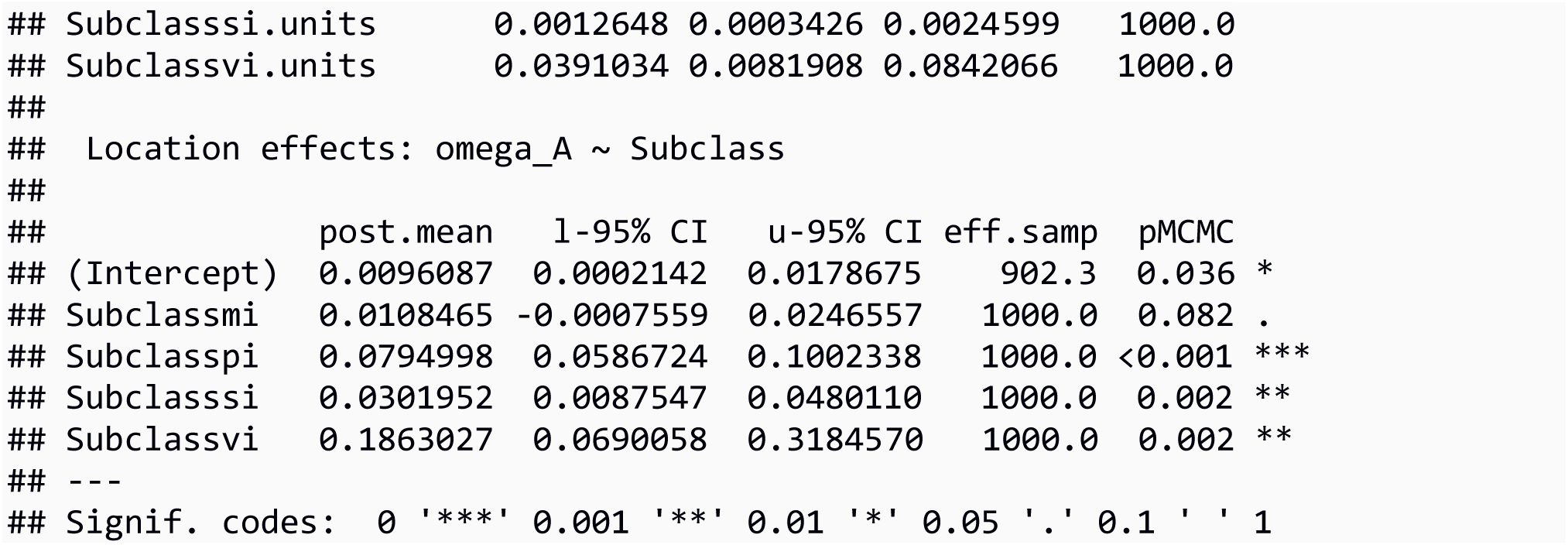

The subclass effects (parameterised as “mi”, “pi”, “si”, and “vi”), along with their pMCMC values are listed under location effects. Differences in residual variances between subpathways can be tested in a similar manner to model 1A. We tested whether certain subpathways were greater than others as previously done with variance components in Model 1A, except using the posteriors for the fixed effects stored in the Sol object. For example:

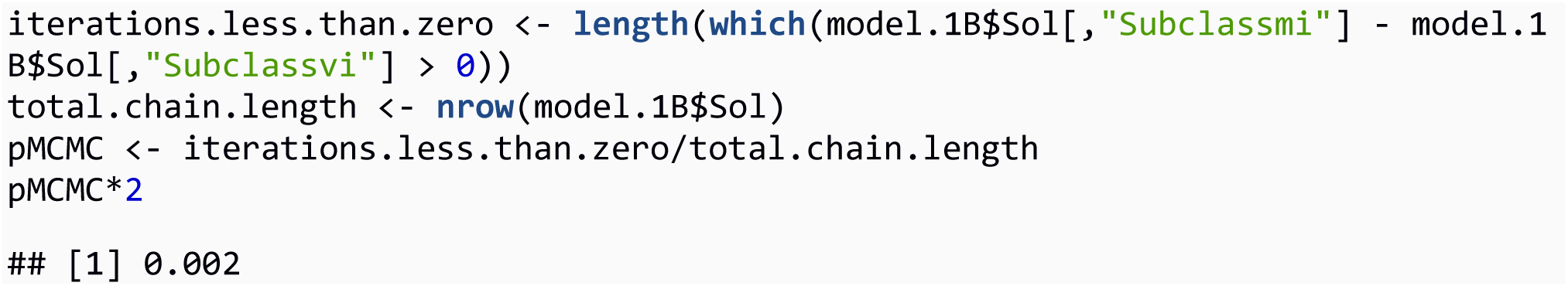

We conclude that the viRNA pathway has a significantly greater rate of adaptive protein evolution (MCMCp = 0.002).

Model 1C: Comparison of RNAi subpathways, with the piRNA split into effectors, biogenesis factors, and transcriptional silencing factors.

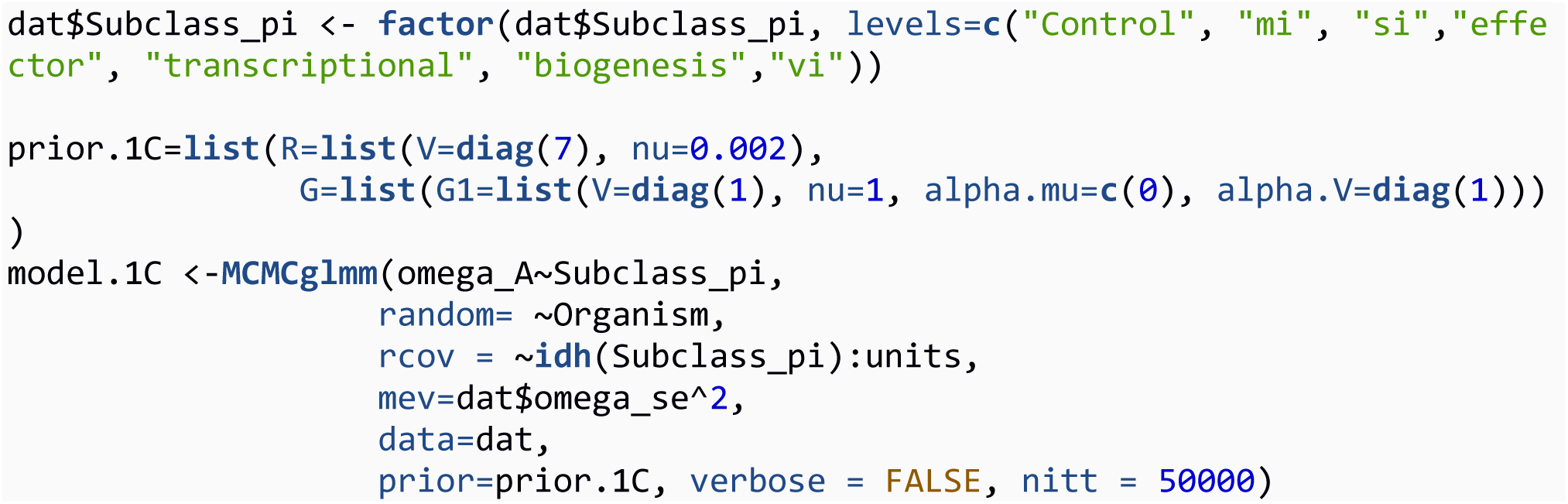

Again, Model 1C is structurally identical to Model 1A and Model 1B, with the only difference being the number of factor levels which genes are grouped into.

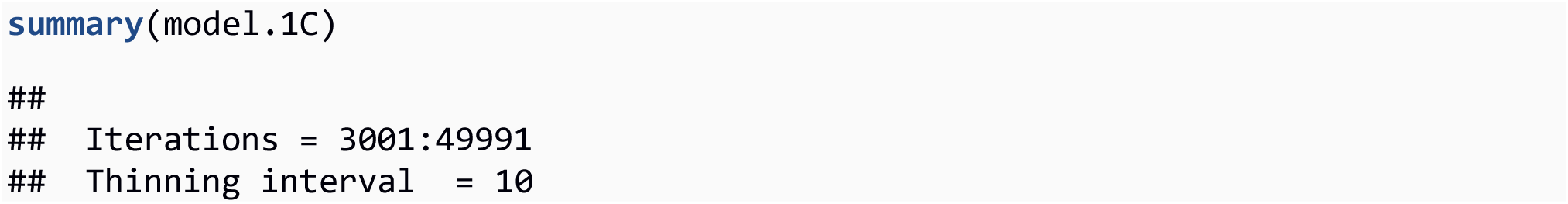
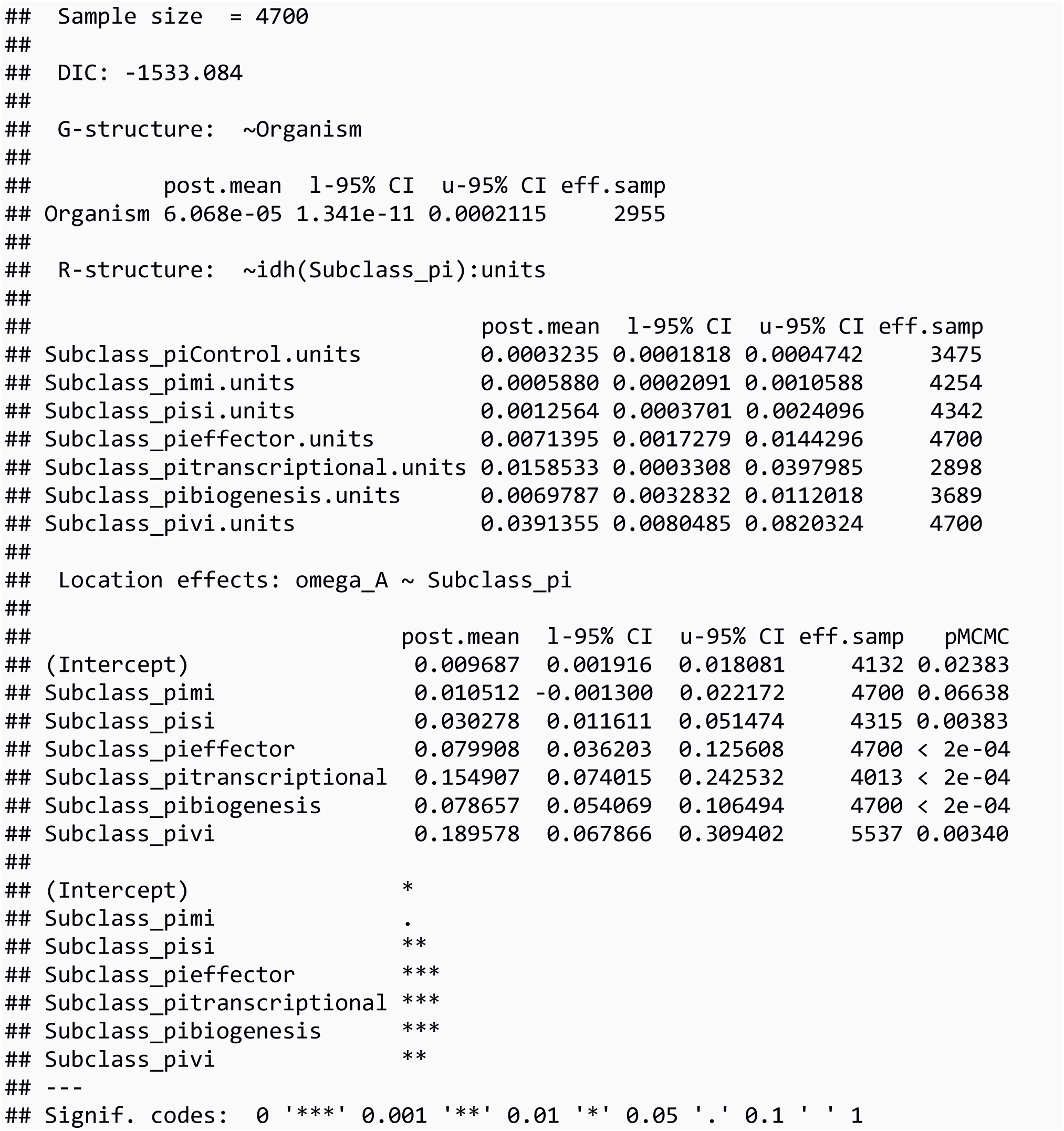

All three piRNA pathways are significantly greater than control genes. Significance between subpathways and variance components was assessed as above.

Model 2A: Comparison of RNAi homologues (with subpathway as a fixed effect)

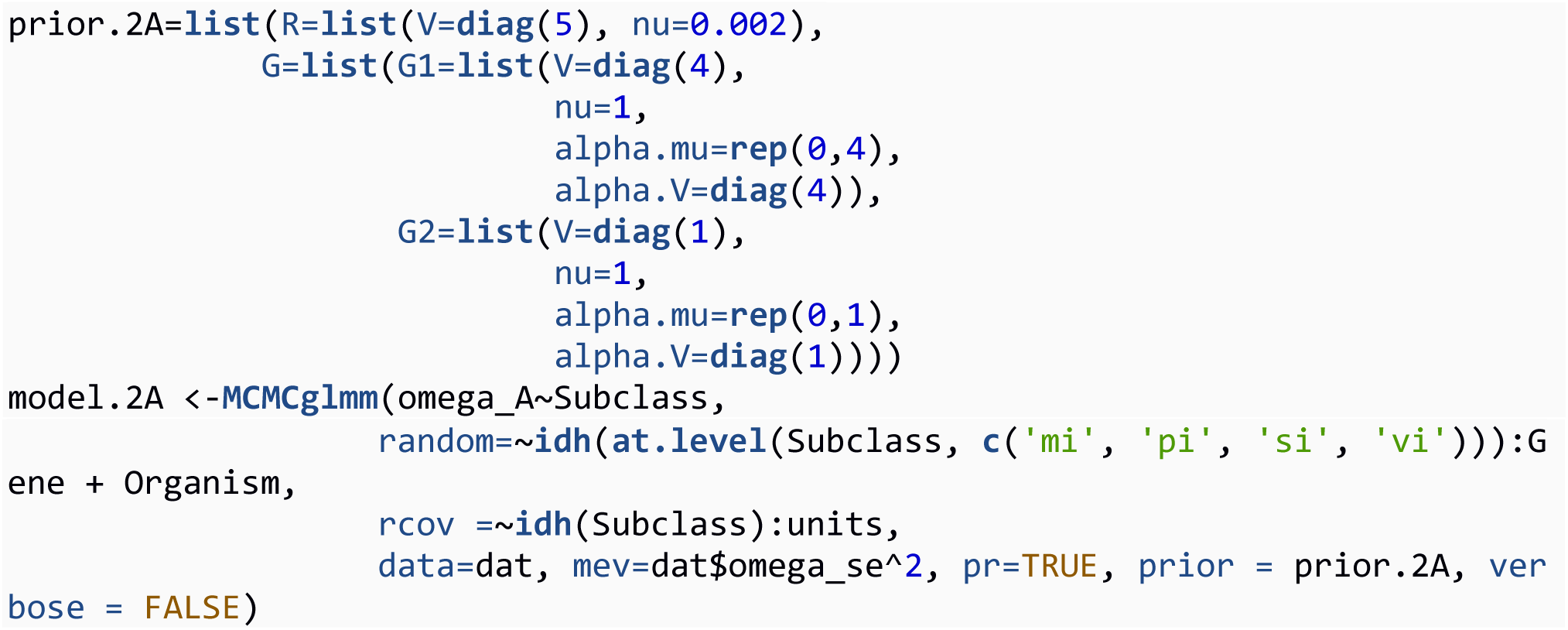

The second model is similar to the first, but for two differences. First, the effect of each RNAi homologue is estimated across species, specifying idh(at.level(Subclass, c(‘mi’, ‘pi’, ‘si’, ‘.vi’))) so that the homologue effect is not estimated for the (nonhomologous) control genes. Second, we specify pr=TRUE, so that the random effects are stored along with the fixed effects in the model output.

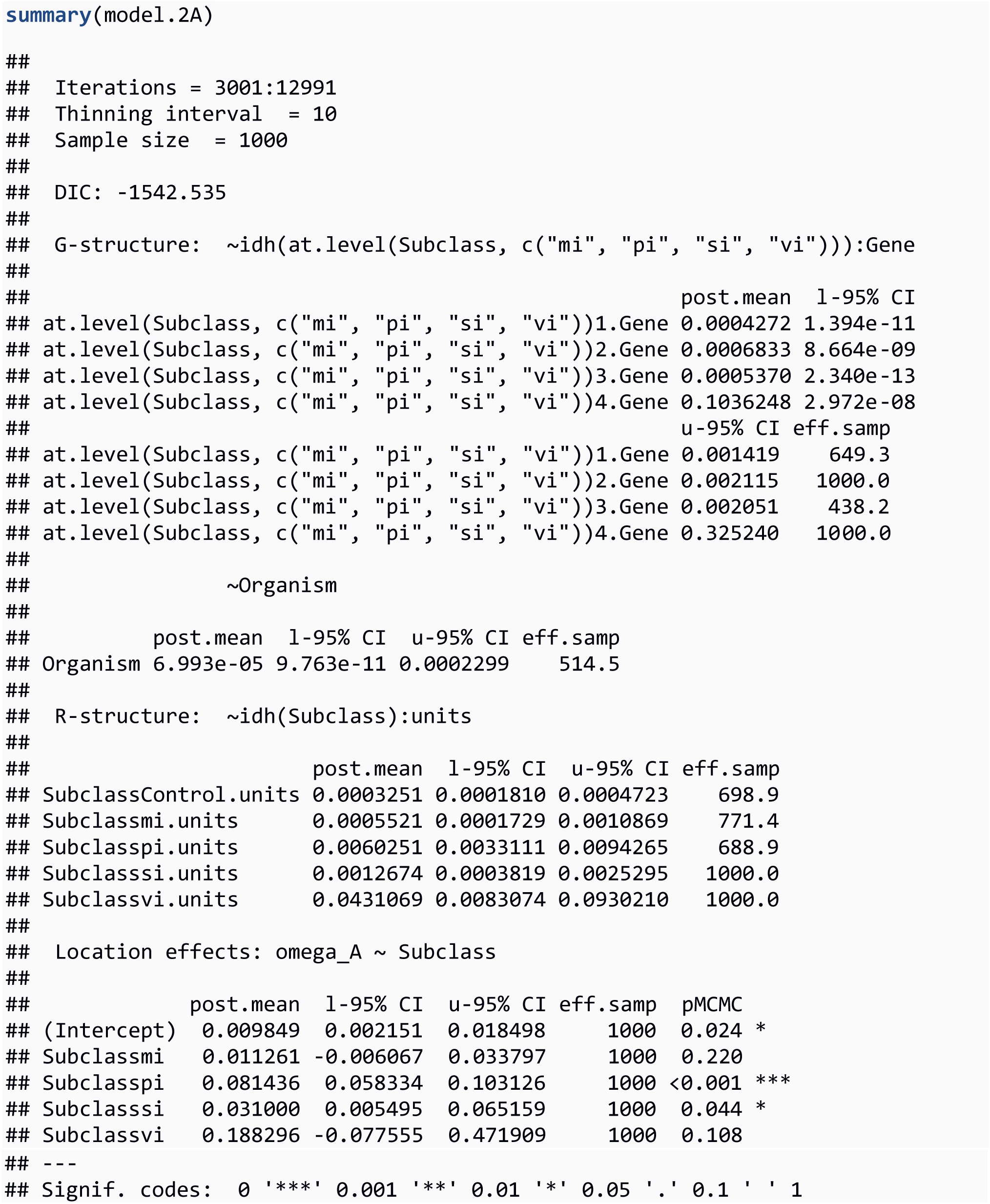

The posterior distribution of the fixed and random effects of the model are stored in the object named model.2A$Sol, with columns pertaining to each distribution. For example:

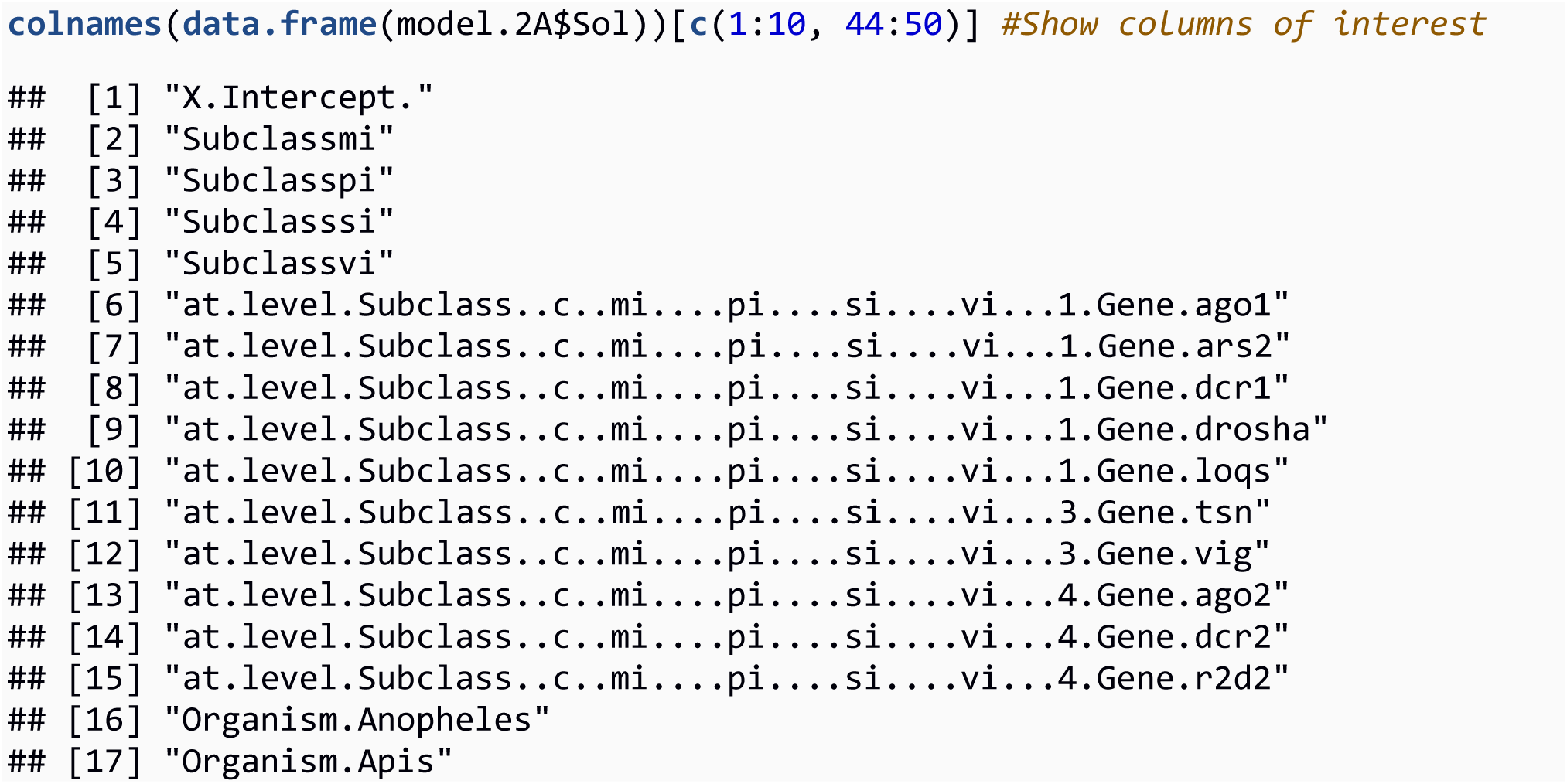

We obtain the posterior distribution for ωA of an individual homologue by adding the posterior distributions for the intercept, subclass, and homologue. For example, the ωA posterior for Argonaute-2 is:

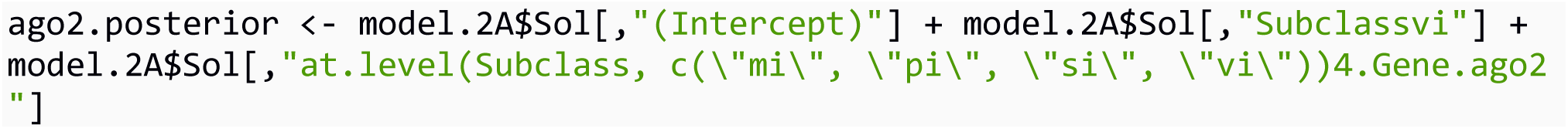

We obtain 95% HPD confidence intervals using the command HPDinterval():

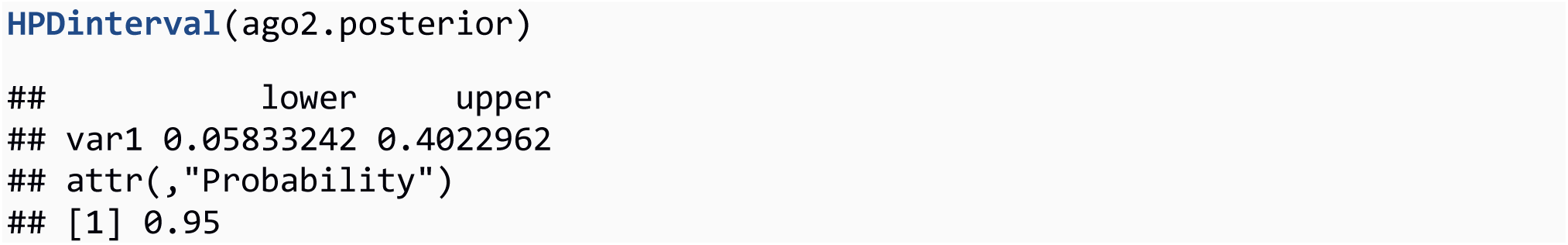

This show the lower 95% HPD interval (0.07) is greater than 0. We test whether this is greater than control genes by subtracting the posterior of ωA estimates of Argonaute-2 from the control gene class posterior, and see the proportion of MCMC intervals where it overlaps zero.

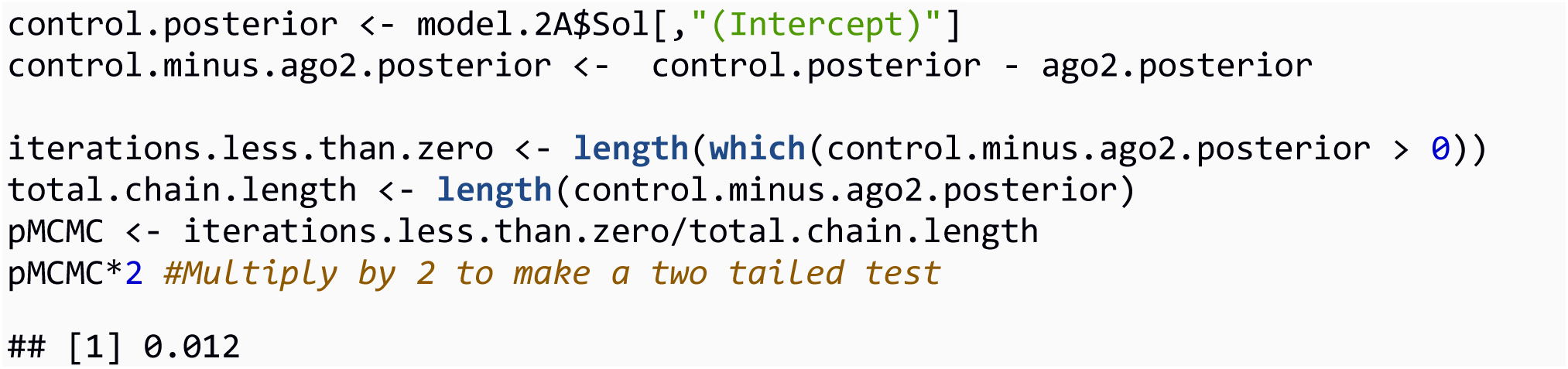

We conclude Argonaute-2 has a greater adaptive rate than control genes (pMCMC = 0.012).

Model 2B: Comparison of RNAi genes

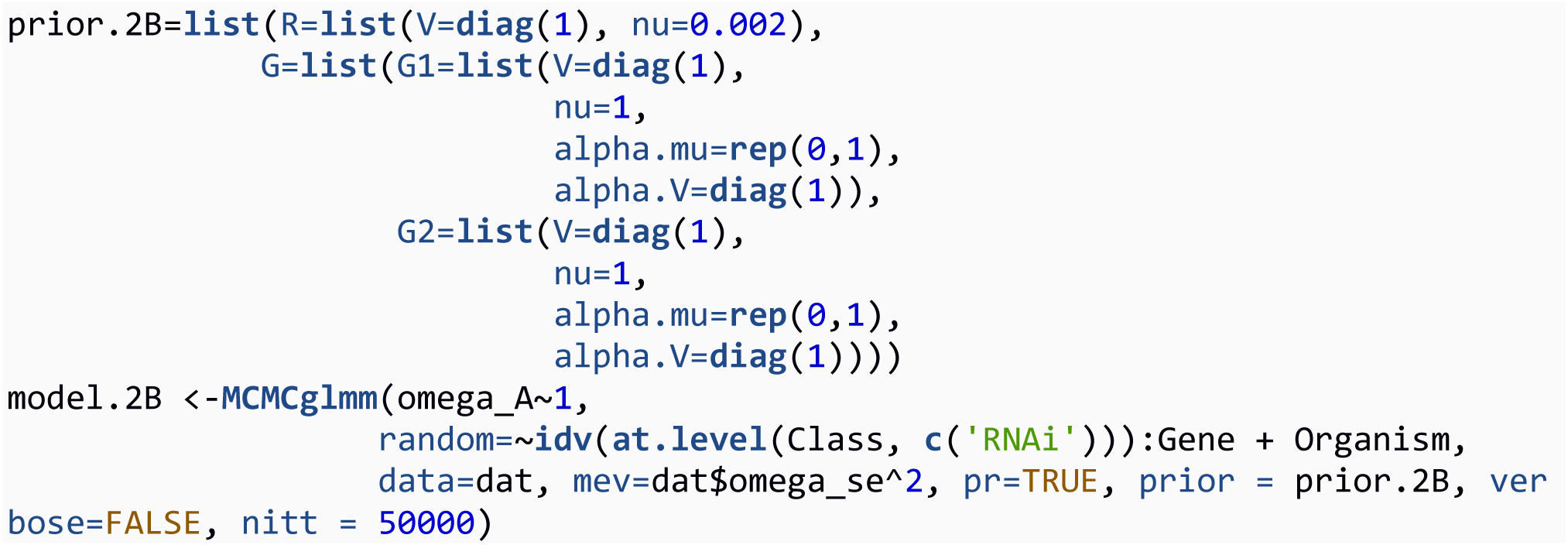

We also parameterise the second model without assigning genes to subpathways, and remove the subclass fixed effect and subclass-specific error-variances. The following is the output of the model:

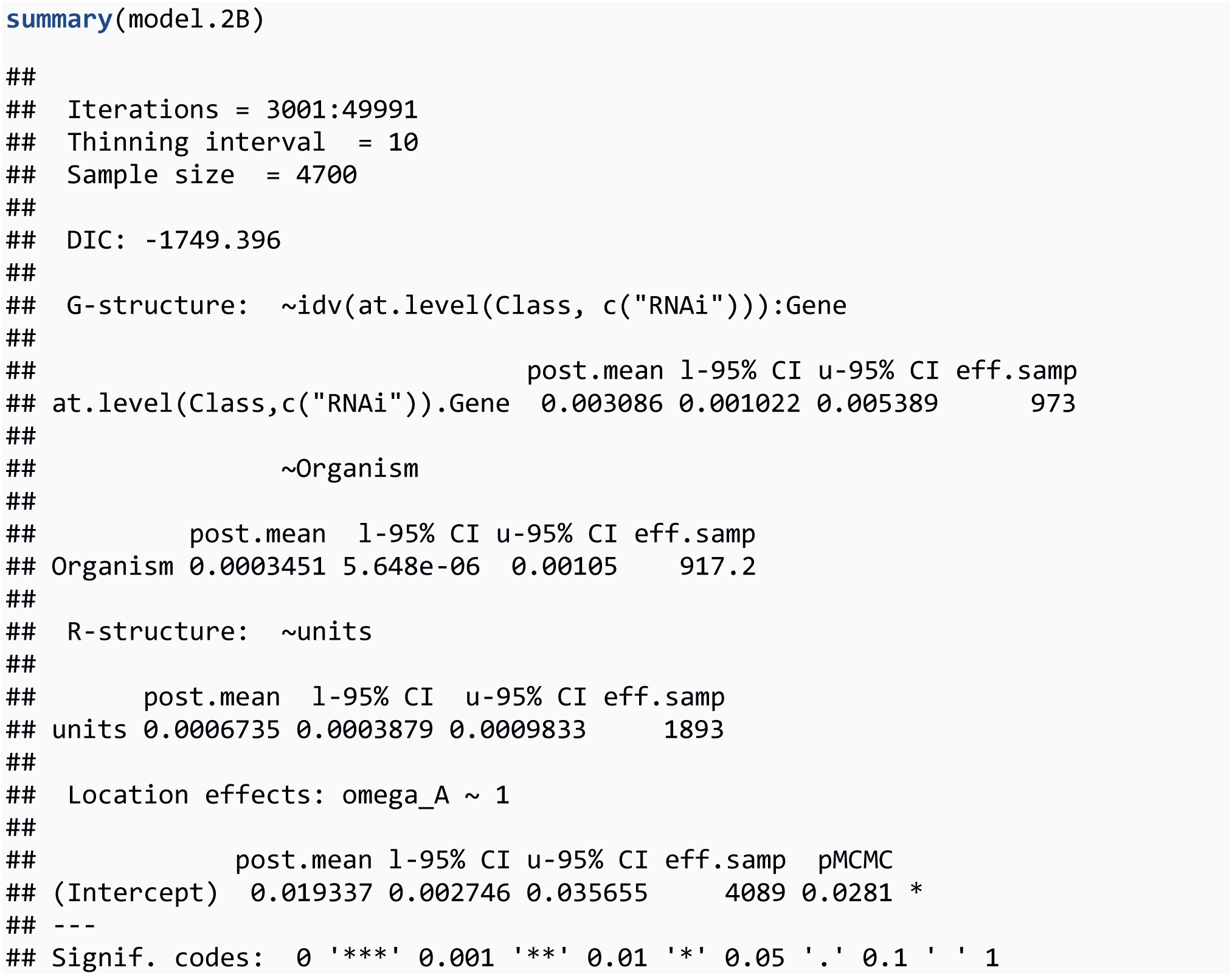

Without a subclass effect, we assess whether a gene has a significantly elevated rate of adaptive amino acid evolution by comparing the posterior distributions of each gene effect to zero. For example, for Argonaute-2:

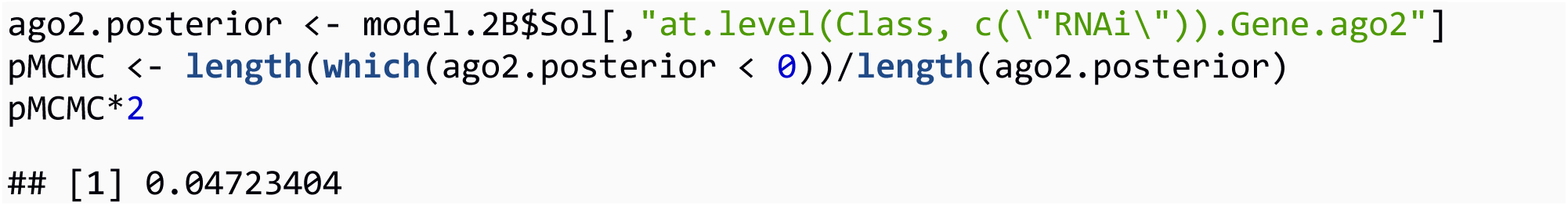

We conclude that Argonaute-2 has significantly elevated ωA (pMCMC = 0.047).

SnIPRE-like analysis

Model 3A: SnIPRE-like analysis, with subpathway as a fixed effect

Set up data

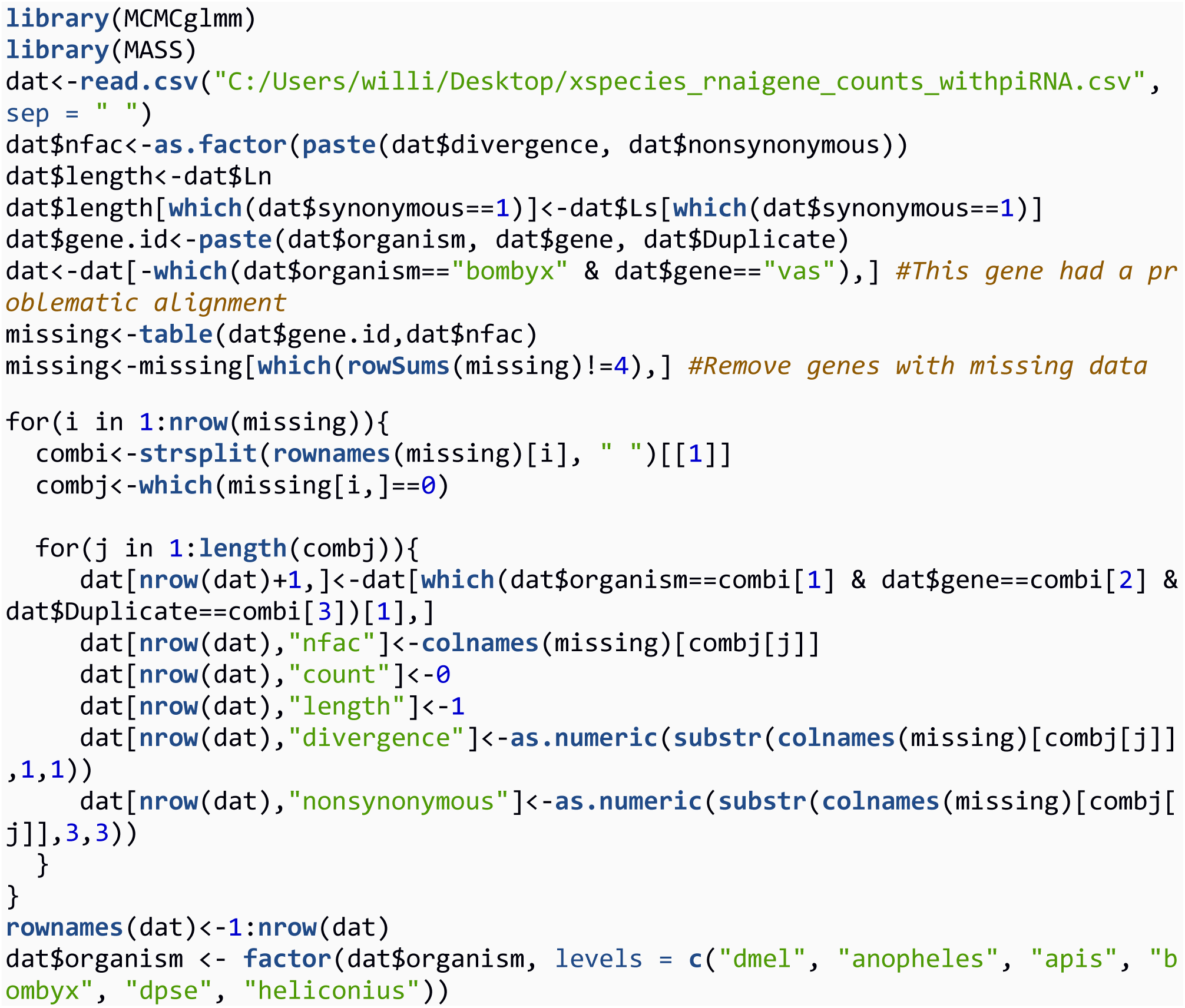

The data look like the following. Each gene in each species is represented by 4 rows of count data, one for each of the MK observations of polymorphism and divergence by synonymous and nonsynonymous mutation types.

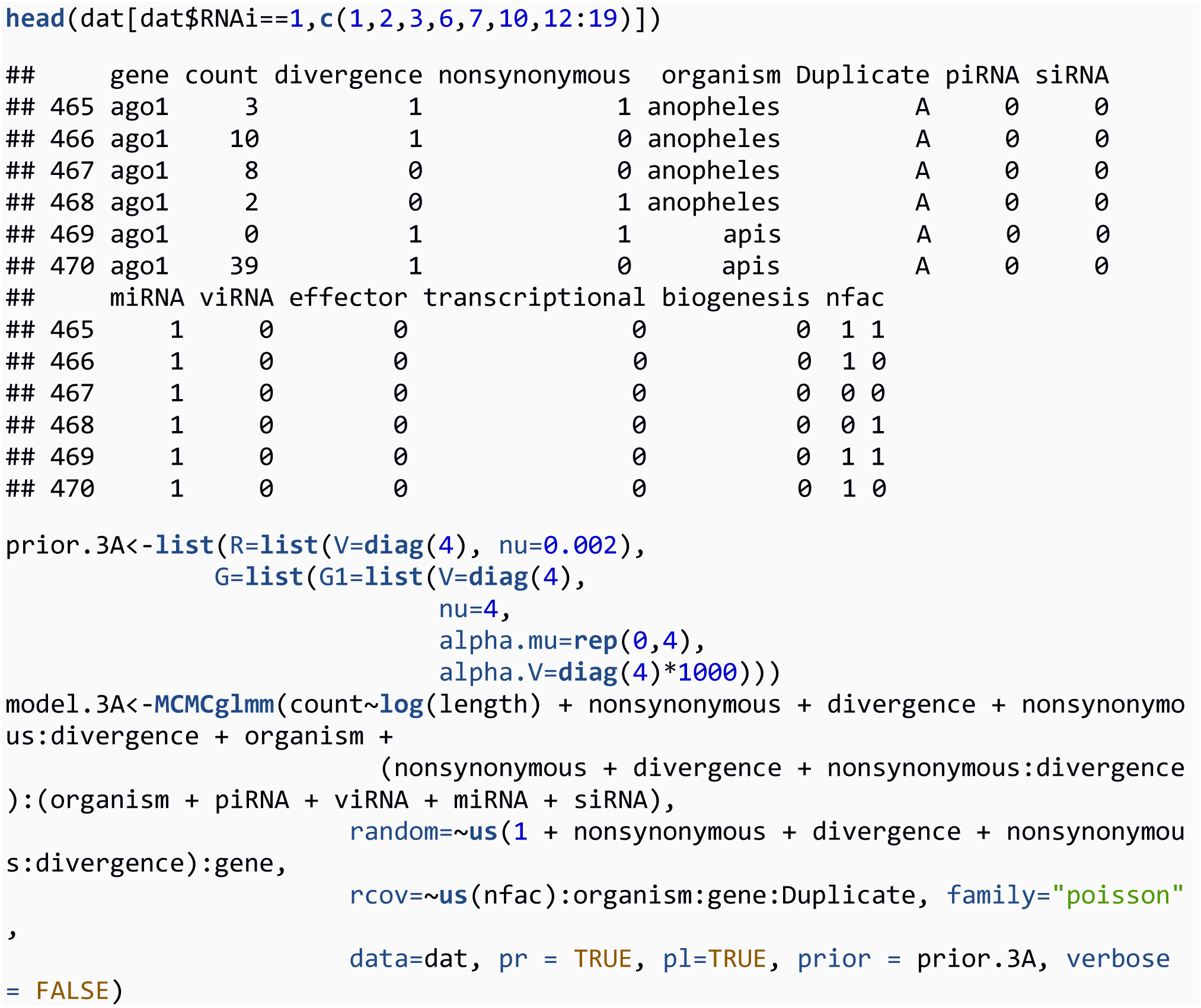

For the SnIPRE-like analysis, we model the counts of each type of mutation (Pn, Ps, Dn, Ds) in each gene in each organism as a poisson response variable. Following Eilertson et al (2012), we set the fixed effects to the length of the gene, the type of mutation (by fitting either a nonsynonymous or divergence effect), and the interaction between nonsynonymous and divergence effects. Because we are interested in estimating effects of certain pathways across species, we also fit nonsynonymous, divergence and nonsynonymous-by-divergence effects separately for each gene class and organism with the term nonsynonymous+divergence+nonsynonymous:divergence):(organism+piRNA+siRNA+miRNA+viRN

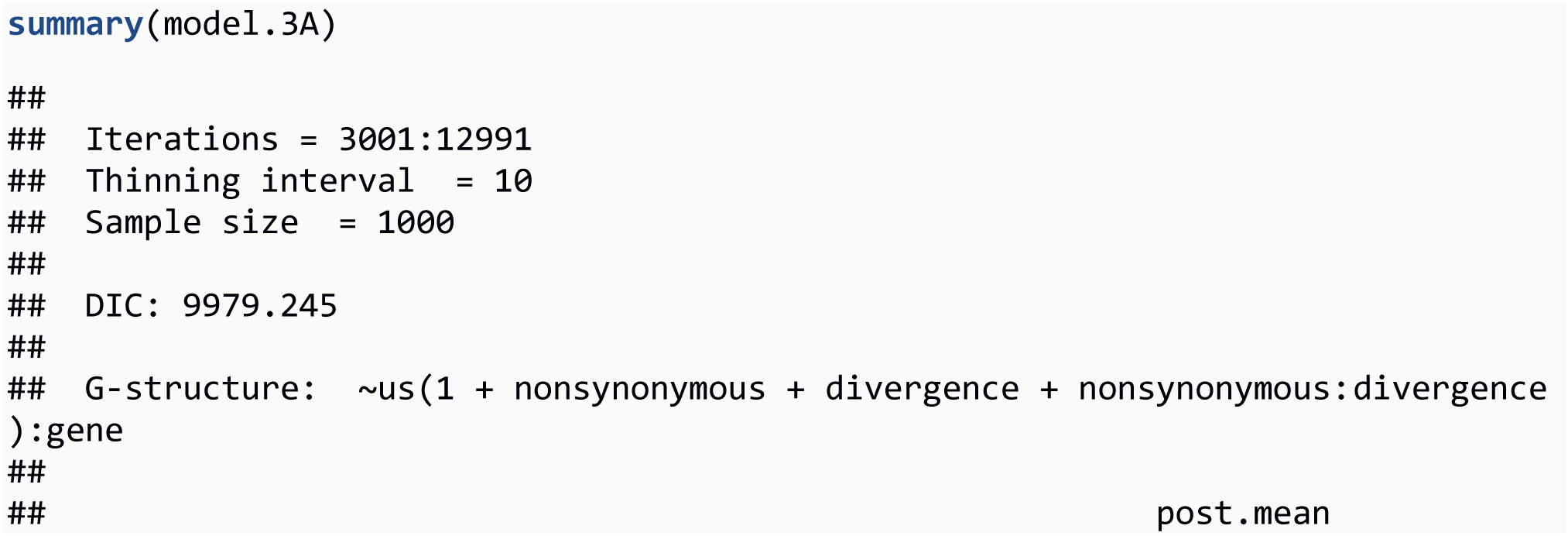
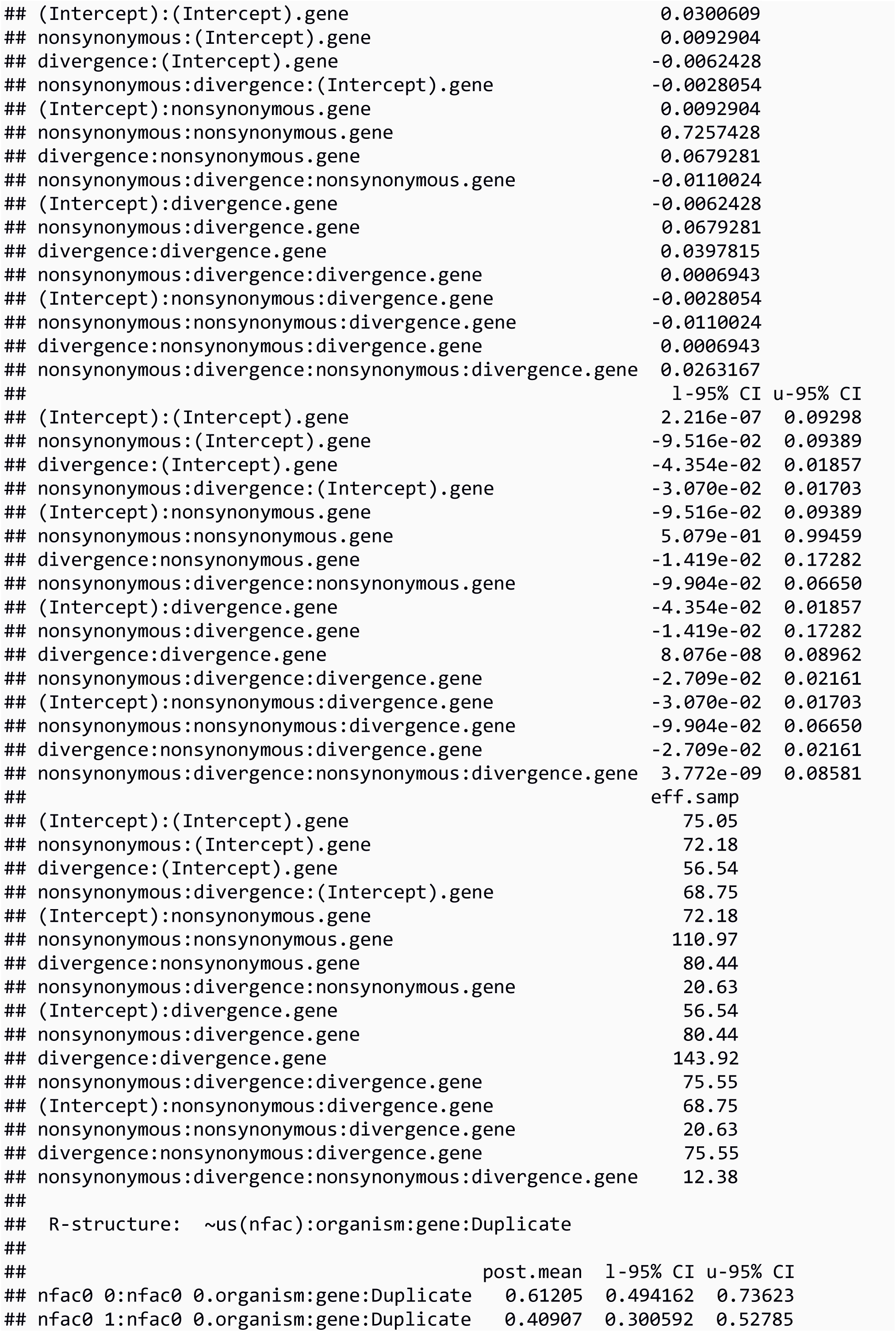
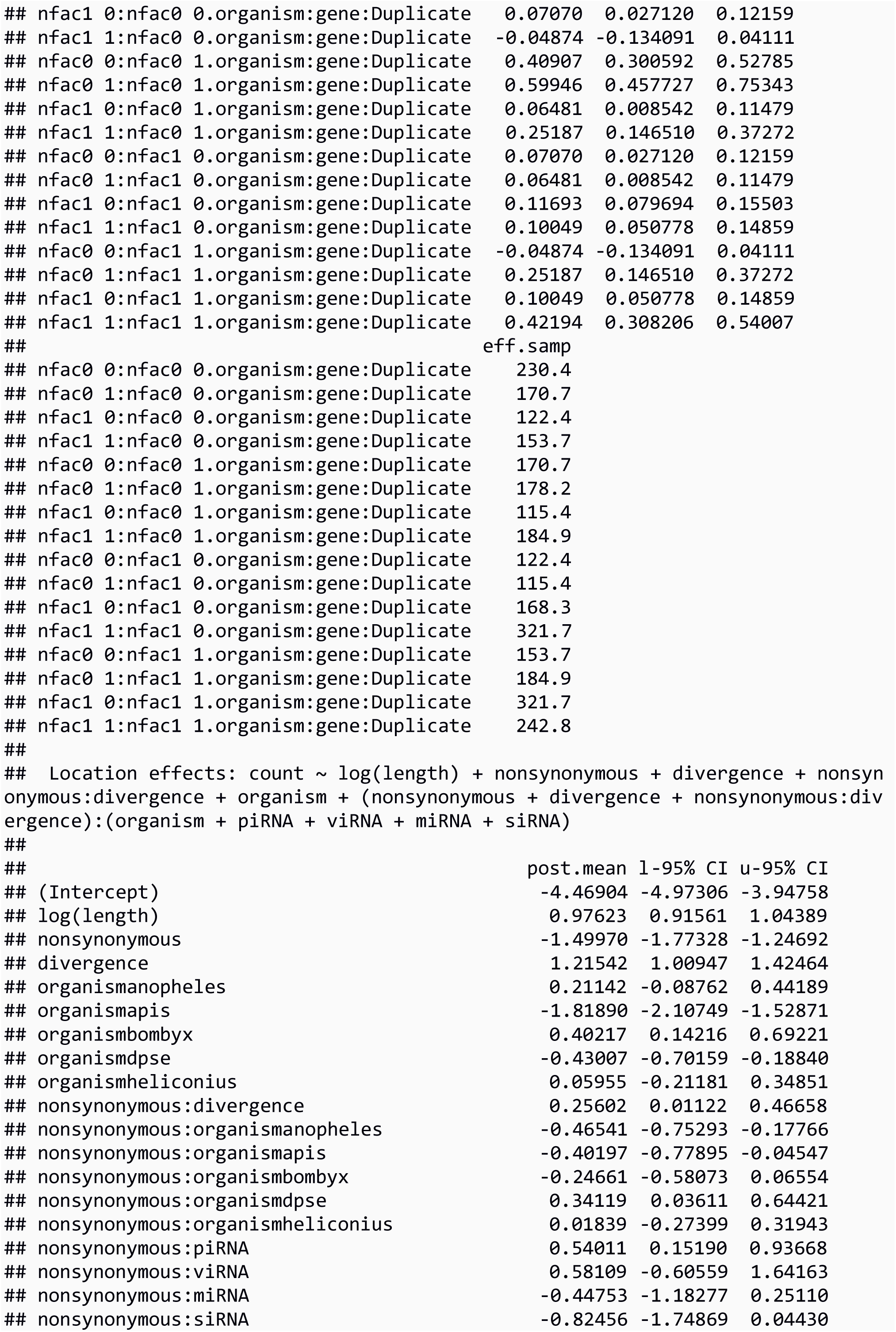
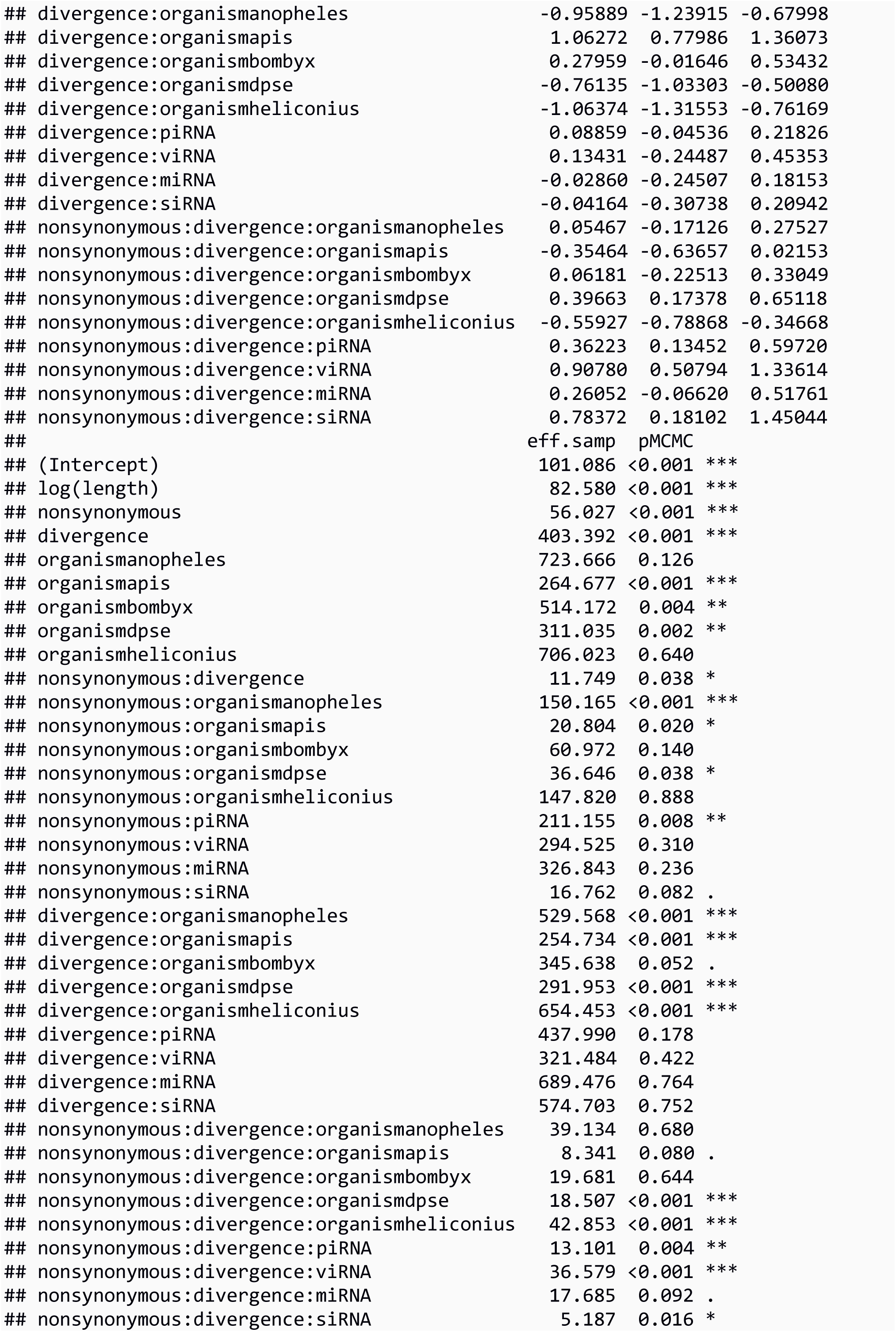
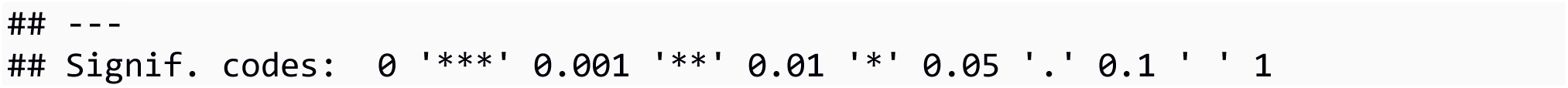

The model output shows organisms (e.g. nonsynonymous:divergence:organismapis effect) and subpathways (e.g. nonsynonymous:divergence:viRNA effect) differ in their genome-wide level of positive selection. We test whether a homologue has an increased selection effect (e.g. Figure 4, Figure S4) by comparing the posterior distributions of the selection effect for control genes and the homologues. For example:

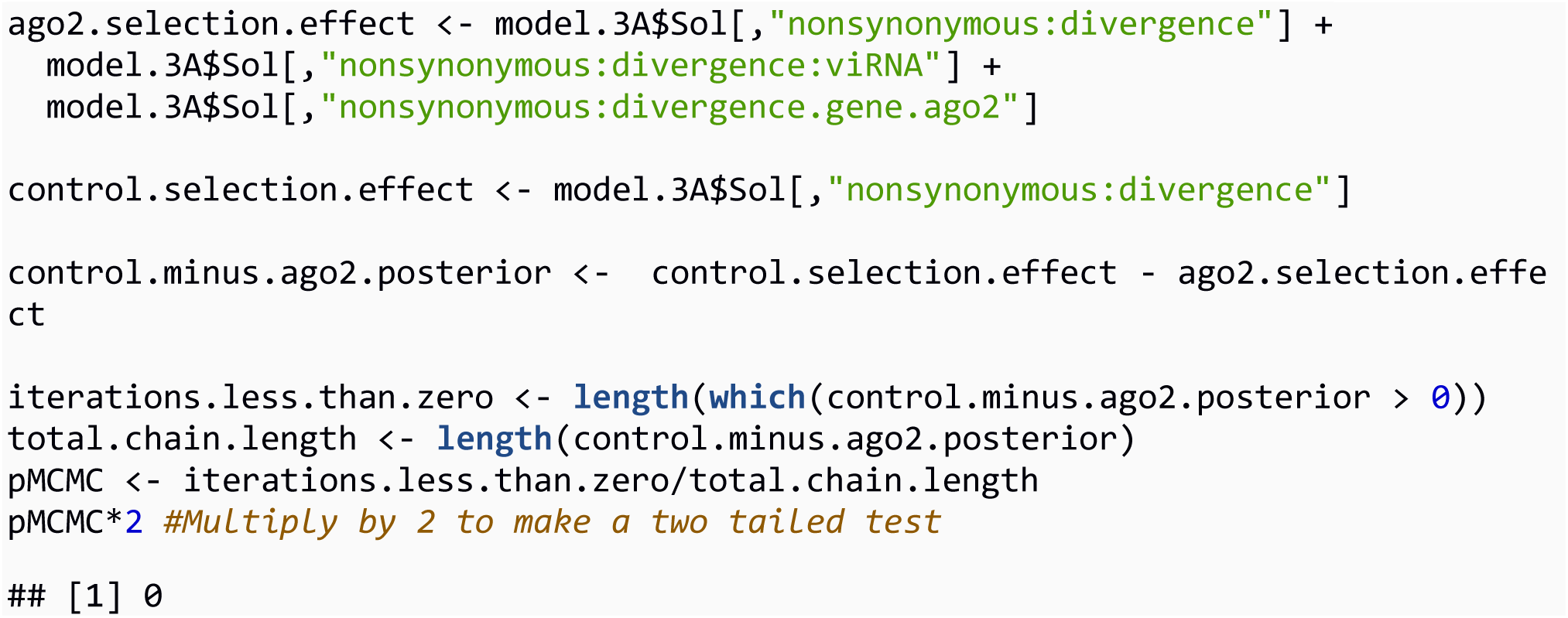

We conclude that the Ago2 selection effect is greater than control genes (MCMCp < 0.001).

In addition to homologue-specific random effects, we include gene-specific random effects in the SnIPRE model. The homologue-specific random effects are coded as “random=~us(1 + nonsynonymous+divergence+nonsynonymous:divergence):gene”, very similar to the random effect structure of Eilertson et al (2012), except we have an average of 24 observations per gene (6 species by 4 types of mutation). Instead of using the per-gene random effect structure from Eilertson et al (2012), we reparameterise the model so that residuals for each class of mutation would be estimated for each gene, and (by specifying pl=TRUE), the posterior distribution of each of these residuals stored. We code the different classes of mutation in the nfac column, where “0 0” denotes synonymous polymorphism, “0 1” denotes nonsynonymous polymorphism, and so on. The unstructured covariance matrix between mutation classes (us(nfac)) is estimated for each gene (organism:gene:Duplicate), from which we extract the gene-level selection effect. The posterior distributions of these residuals are saved in the model.3A$Liab data structure, with columns in the same order as the rows in our original data table:

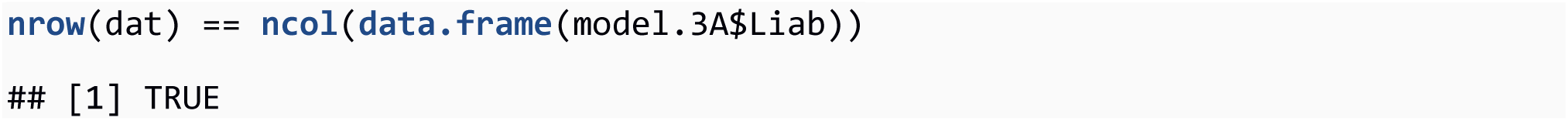

To extract the gene specific selection effect from each residual, we create the mapping matrix (X), the model matrix for all nonsynonymous:divergence effects (X.matrix, a template for which fixed effects link with the rows of data), the columns of the model.3A fixed effects which match X.matrix (X.model.hit), and a list of unique genes (unique.genes):

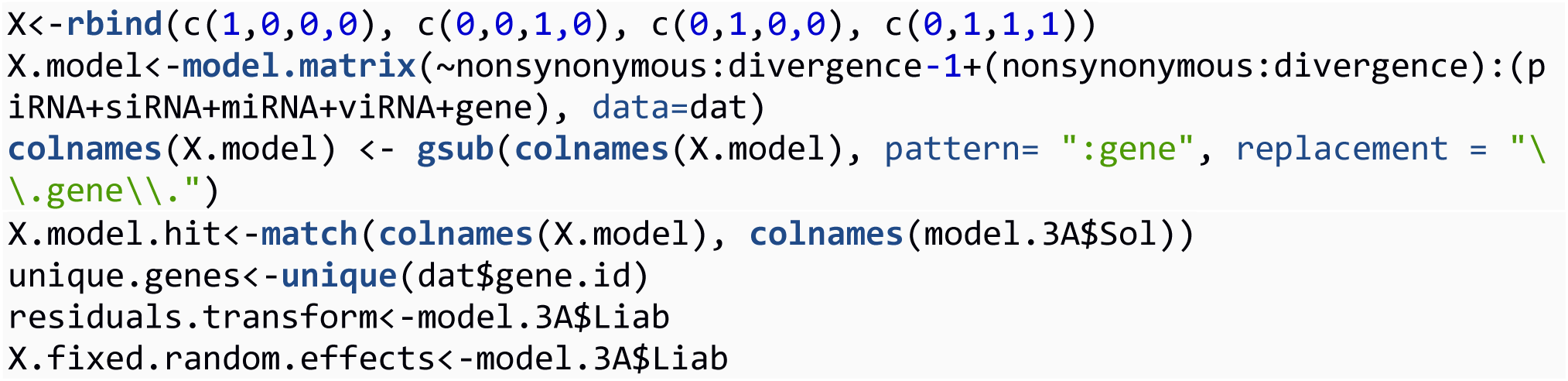

We loop through rows of the stored latent variables (model.3A$Liab), with each row being an iteration of the MCMC chain, and subtract the fixed and random effects corresponding to each data point ((model.3A$X%*%model.3A$Sol[i,1:ncol(model.3A$X)])) from the ωA estimate (model.3A$Z%*%model.3A$Sol[i,(ncol(model.3A$X)+1):ncol(model.3A$Sol)])), resulting in residuals for each gene in each species. Then, for each gene (gene.id - a combination of gene name, organism, and duplicate) in each iteration, we map the residuals onto the design matrix X to solve for the random effects of each observation. (residuals.transform). Finally, we obtain the posterior distribution of the selection effect for a particular gene (selection.effects) by adding the random nonsynonymous:divergence:gene effect (residuals.transform) to the fixed and random nonsynonymous:divergence effects (X.fixed.random.effects).

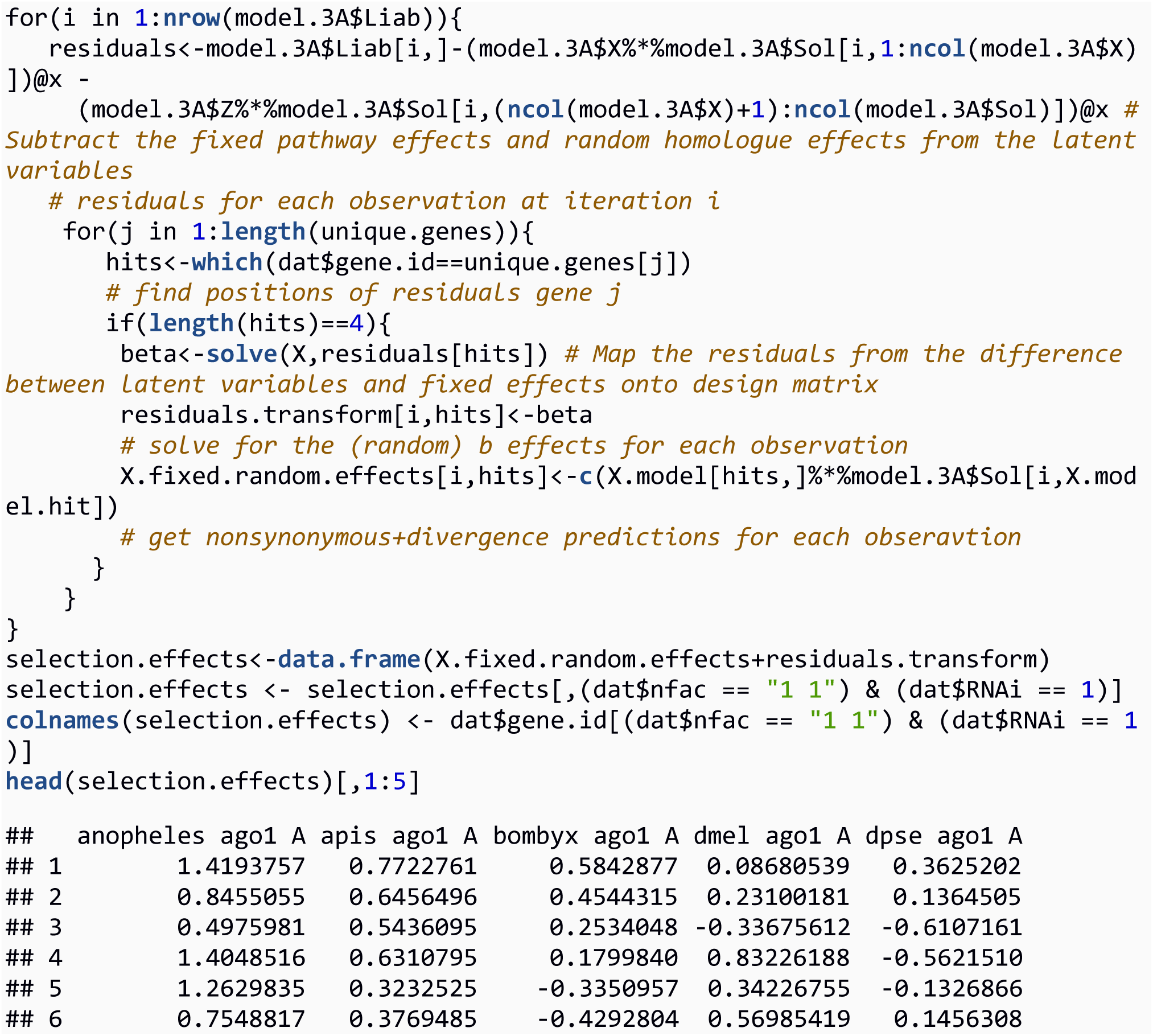

We calculate a “species-corrected” selection effect where nonsynonymous:divergence:organism effects are excluded in order to visualise differences between subpathways (e.g. Figure 3). We add these effects later when assessing positive selection in individual genes within a species. Also, in this example, we have only solved for the gene-level nonsynonymous:divergence random effects, however, the other gene-level effects could be obtained in a similar way.

SnIPRE was originally intended to identify genes in a single organism which shows signs of elevated positive selection. To do this, we add the organism specific selection effect to the “species-corrected” selection effect we have already obtained, and ask whether the selection effect overlaps zero. For example, to estimate the selection effect for the genes in Apis mellifera, we add the nonsynonymous:divergence:apis posterior to the columns of selection.effects which belong to Apis.

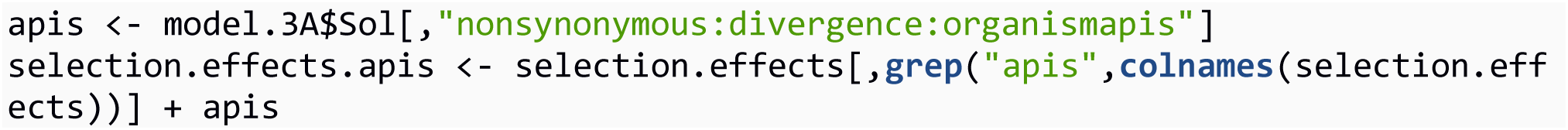

Then, using HPDinterval() and colMeans(), we can get the selection effect for each gene, along with the upper and lower 95% highest posterior density intervals.

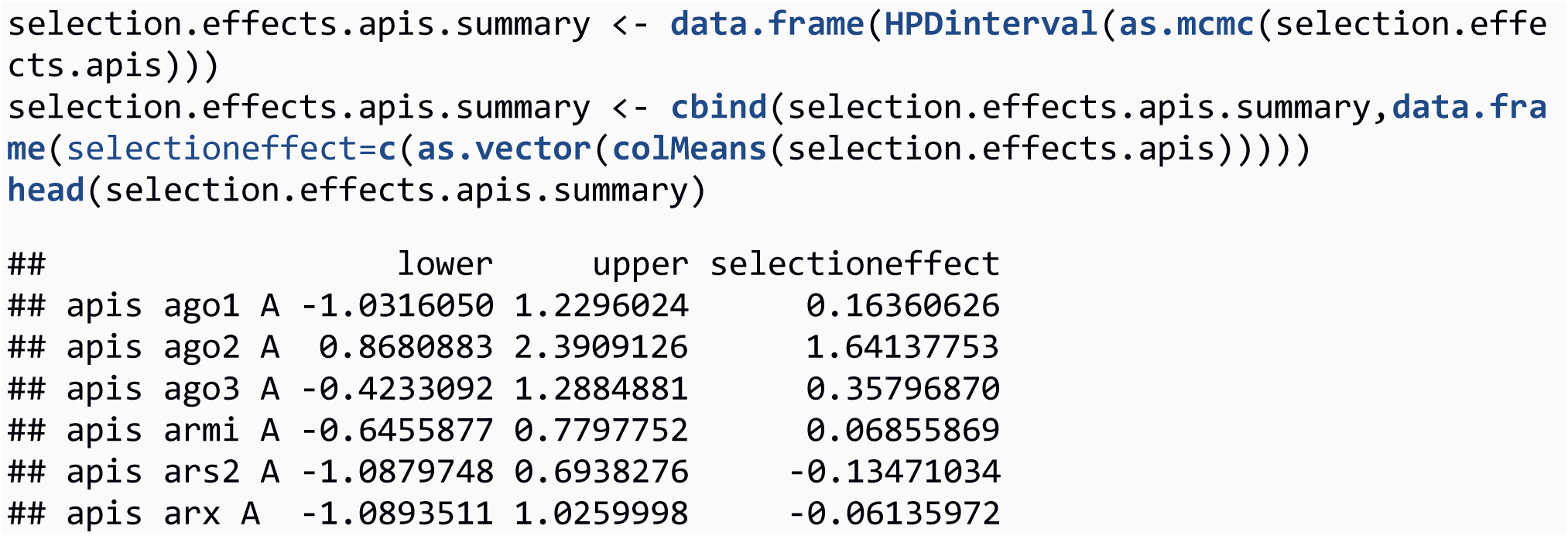

Finally, we identify genes with significantly positive selection effects.

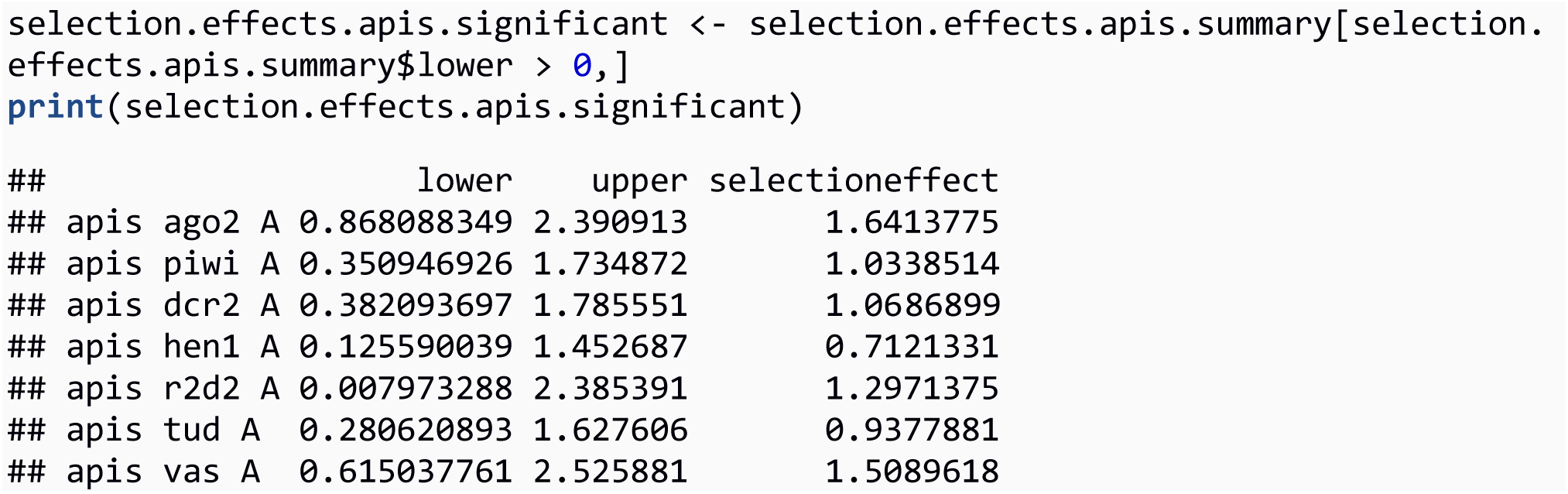

Model 3B: SnIPRE-like analysis, with piRNA split into biogenesis factors, effectors, and transcriptional silencing

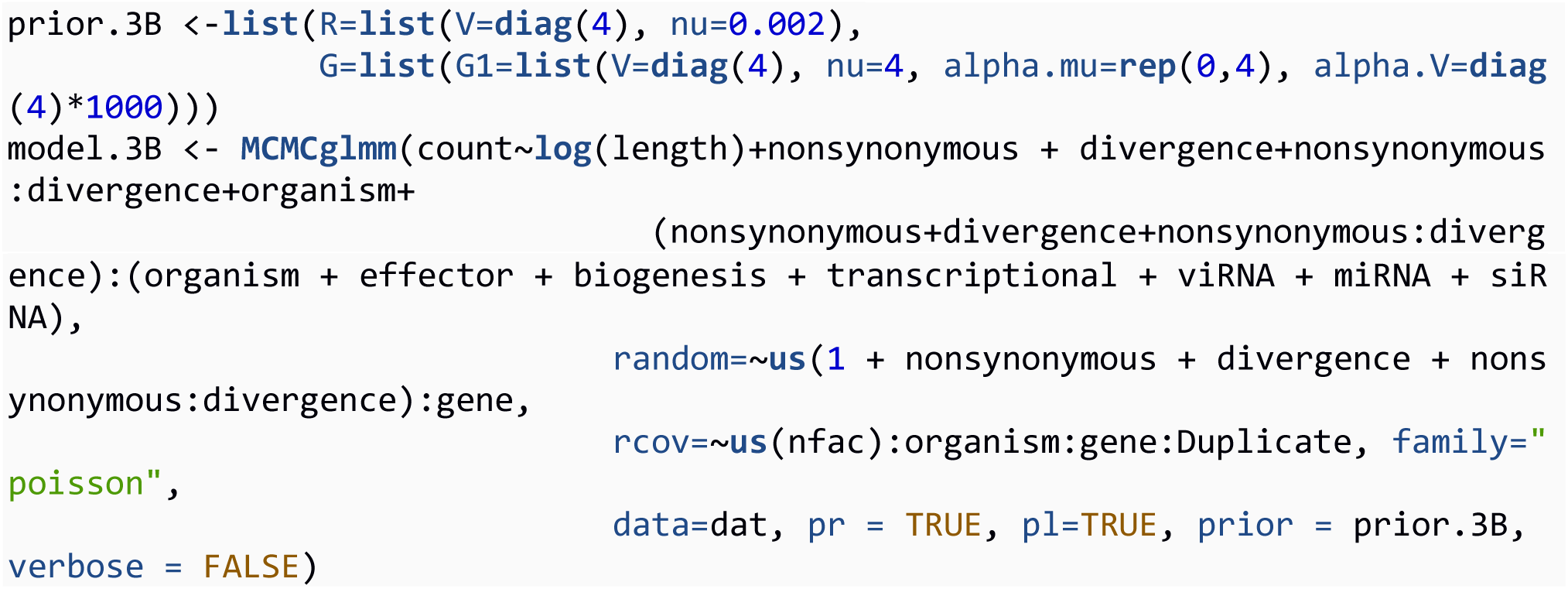

We also fit the SnIPRE model (Model 3A) with the piRNA pathway split into different functional categories, akin to the Model 1B and 1C. We only used this model to estimate the selection effects associated with the piRNA categories (transcriptional silencing, effectors, and biogenesis machinery).

Model 3C: SnIPRE-like analysis, without subpathway as a fixed effect

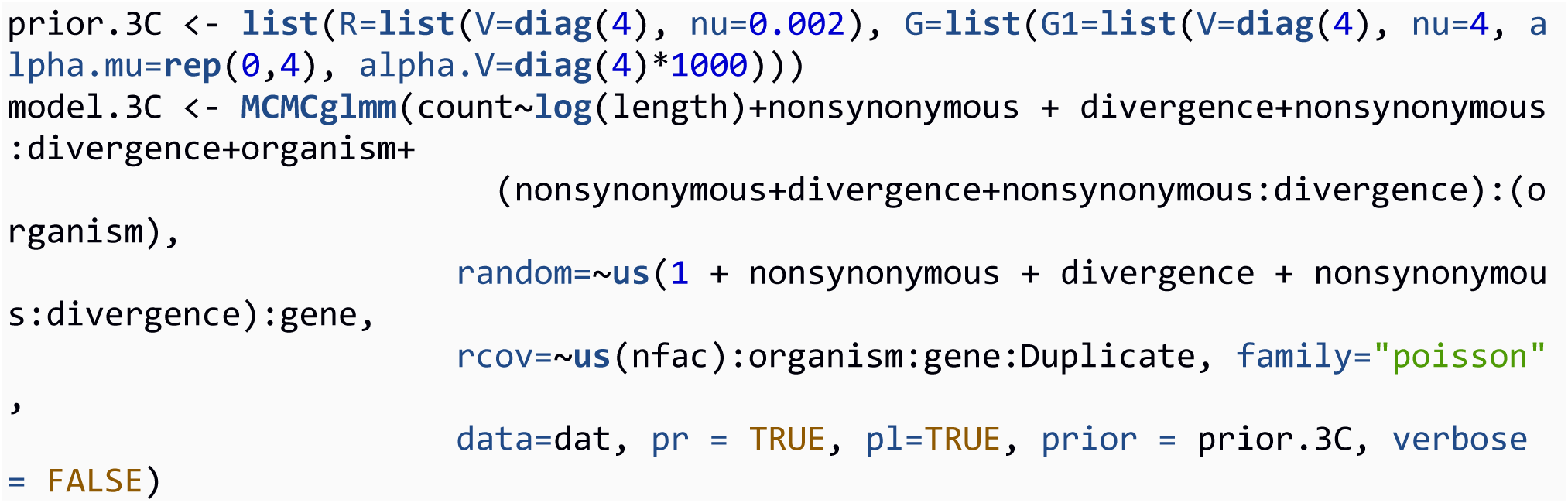

Finally, we fit the SniPRE model (Model 5A) without assuming genes belong to any particular subpathway,similar to the difference between Model 2A and 2B. Selection effects were then calculated in the same way, excluding the addition of a subpathway fixed effect.

## S2 Text: Supplementary R code for models

To assess significance in the SweeD analyses, we used ms (Hudson, 2002) to perform 1000 coalescent simulations for each gene region of interest in each species, given the observed number of segregating sites, reported recombination rate, and a previously published estimate of the demographic history of that species. When population scaled recombination rate estimates were not available, we used estimates of *N_e_* to scale per-base rate estimates. Although the details of the demographic scenarios we modelled are unlikely to impact substantially upon our qualitative comparisons of between sweep frequency in different types of gene, we attempted to use null models consistent with the published literature. The demographic scenarios modelled for each species are illustrated in Figure S1. For *D. melanogaster*, recombination rates from the *Drosophila* recombination rate calculator were used with a constant *N_e_* for African populations of 1.15x10^6^ (Charlesworth, 2009). Some genes (*ael, AGO3, pasha,* and *Rm62*) are reported to lie in areas with zero recombination (Fiston-Lavier, et al., 2010), so we set the recombination rate in these genes at the lowest non-zero rate observed. For *D. pseudoobscura*, we simulated a population expansion (Haddrill, et al., 2010; Larracuente & Clark, 2014), and used the population scaled rates of recombination and gene conversion from Larracuente and Clark (2014). For *Anopheles gambiae,* we used demographic history parameters from Crawford and Lazzaro et al (2010) for the Cameroon population, and the recombination rates for each individual chromosome arm (1 cM/Mb for the X, 1.3 cM/Mb for 3L and 2R, 1.6 cM/Mb for 3R, and 2 cM/Mb for 2L) from Pombi et al (2006) and Stump et al (2007). Effective population size (*N_e_*) was set to 2.4x10^6^ estimated using the *D. melanogaster* mutation rate of Keightley et al (2014) and the Watterson’s theta (θ_W_) estimate in Crawford and Lazzaro (2010). For *H. melpomene,* we simulated three Costa Rican populations corresponding to *H. melpomene, H. cydno, and H. pachinus,* using the migration rates provided in Table 2 of Kronforst et al (2006). We used a constant recombination rate of 7.51 cM/Mb across the entire genome with an *N_e_* of 2.1x10^6^ for *H. melpomene*, 3.3x10^6^ for cydno, and 2.7x10^6^ for *H. pachinus*. For *B. mandarina,* we modelled the “gene-flow at bottleneck” scenario (Yang et al, 2014), with an *N_e_* of 500,000 for *B. mandarina* and 73,000 for *B. mori*, and a recombination rate of 2.97 cM/Mb (Yamamoto et al, 2008; Yang et al, 2014). For *A. mellifera,* four subpopulations were modelled using *N_e_* values in Table 1 of Wallberg et al (2014), following Figure 1F in Wallberg et al (2014) when modelling past subpopulation size changes. These subpopulations share migrants, and migration rates were estimated based on *F*_ST_ values between subpopulations reported in Whitfield et al (2006). A recombination rate of 19 cM/Mb is assumed to be constant across the genome (Beye et al, 2006). For *C. briggsae*, coalescent simulations and SweeD analyses were carried out on the 25 “tropical” samples in order to avoid modelling complicated demographic scenarios. These are expected to have an effective population size of 60,000, and to have undergone a recent bottleneck 0.916 *N_e_* generations in the past (Cutter et al, 2006; Denver et al, 2009), assuming a 60-day generation time (Barrière & Félix, 2005). We used recombination rates for *C. briggsae* from Ross et al (2011), which are estimated to be 9.97 x 10^−8^ per bp per generation in autosomes and 4.6 x 10^−8^ per bp per generation on the X chromosome (Ross et al, 2011). Finally, for *P. pacificus*, four subpopulations were modelled corresponding to clade A1, A2, C, and 9 individuals whose clade was unknown (Rödelsperger, et al., 2014) which coalesced 0.849 *N_e_* generations in the past (McGaughran et al, 2013). *N_e_* was estimated by calculating θ_W_ for each contig and assuming a mutation rate of2×10^−9^ (Weller et al, 2014).To minimise differences between the real data and simulations, sites were randomly chosen to be folded, ancestrally invariant, or fixed for a derived substitution, in each case matching the numbers observed in the real data before the SweeD analysis.

